# Dimensionality reduction distills complex evolutionary relationships in seasonal influenza and SARS-CoV-2

**DOI:** 10.1101/2024.02.07.579374

**Authors:** Sravani Nanduri, Allison Black, Trevor Bedford, John Huddleston

## Abstract

Public health researchers and practitioners commonly infer phylogenies from viral genome sequences to understand transmission dynamics and identify clusters of genetically-related samples. However, viruses that reassort or recombine violate phylogenetic assumptions and require more sophisticated methods. Even when phylogenies are appropriate, they can be unnecessary or difficult to interpret without specialty knowledge. For example, pairwise distances between sequences can be enough to identify clusters of related samples or assign new samples to existing phylogenetic clusters. In this work, we tested whether dimensionality reduction methods could capture known genetic groups within two human pathogenic viruses that cause substantial human morbidity and mortality and frequently reassort or recombine, respectively: seasonal influenza A/H3N2 and SARS-CoV-2. We applied principal component analysis (PCA), multidimensional scaling (MDS), t-distributed stochastic neighbor embedding (t-SNE), and uniform manifold approximation and projection (UMAP) to sequences with well-defined phylogenetic clades and either reassortment (H3N2) or recombination (SARS-CoV-2). For each low-dimensional embedding of sequences, we calculated the correlation between pairwise genetic and Euclidean distances in the embedding and applied a hierarchical clustering method to identify clusters in the embedding. We measured the accuracy of clusters compared to previously defined phylogenetic clades, reassortment clusters, or recombinant lineages. We found that MDS embeddings accurately represented pairwise genetic distances including the intermediate placement of recombinant SARS-CoV-2 lineages between parental lineages. Clusters from t-SNE embeddings accurately recapitulated known phylogenetic clades, H3N2 reassortment groups, and SARS-CoV-2 recombinant lineages. We show that simple statistical methods without a biological model can accurately represent known genetic relationships for relevant human pathogenic viruses. Our open source implementation of these methods for analysis of viral genome sequences can be easily applied when phylogenetic methods are either unnecessary or inappropriate.

## 1. Introduction

Tracking the evolution of human pathogenic viruses in real time enables epidemiologists to respond quickly to emerging epidemics and local outbreaks [1]. Real-time analyses of viral evolution typically rely on phylogenetic methods that can reconstruct the evolutionary history of viral populations from their genome sequences and estimate states of inferred ancestral viruses from the resulting trees including their most likely genome sequence, time of circulation, and geographic location [2, 3, 4]. Importantly, these methods assume that the sequence diversity of sampled tips accrued through clonal evolution, that is, the occurrence of mutations on top of an inherited genomic background, that is further inherited by descendant pathogens. In practice, the evolutionary histories of many human pathogenic viruses violate this assumption through processes of reassortment or recombination, as seen in seasonal influenza [5, 6] and seasonal coronaviruses [7], respectively. Researchers account for these evolutionary mechanisms by limiting their analyses to individual genes [8, 9], combining multiple genes despite their different evolutionary histories [10], or developing more sophisticated models to represent the joint likelihoods of multiple co-evolving lineages with ancestral reassortment or recombination graphs [11, 12]. However, several key questions in genomic epidemiology do not require inference of ancestral relationships and states, and therefore may be amenable to non-phylogenetic approaches for summarizing genetic relationships. For example, genomic epidemiologists commonly need to 1) visualize the genetic relationships among closely related virus samples [13, 14], 2) identify clusters of closely-related genomes that represent regional outbreaks or new variants of concern [15, 16, 17, 18], and 3) place newly sequenced viral genomes in the evolutionary context of other circulating samples [19, 20, 21]. Given that these common use cases rely on genetic distances between samples, tree-free statistical methods that operate on pairwise distances could be sufficient to address each case. As these tree-free methods lack a formal biological model of evolutionary relationships, they make weak assumptions about the input data and therefore should be applicable to pathogen genomes that violate phylogenetic assumptions. Furthermore, methods that describe genetic relationships with network-like visualizations may feel more familiar to public health practitioners who are accustomed to viewing contact tracing networks alongside genomic information in tools like MicrobeTrace [14] or MicroReact [13] and for viral pathogens like HIV [22, 23] and SARS-CoV-2 [24, 25]. For this reason, reduced dimensionality representations of genomic relationships may be more easily applied for public health action.

Common statistical approaches to analyzing variation from genome alignments start by transforming alignments into either a matrix that codes biallelic nucleotide variants as binary integers (0 for reference alleles and 1 for alternate alleles) [26] or a distance matrix representing the pairwise distances between sequences [27]. The first of these transformations is the first step prior to performing a principal component analysis (PCA) to find orthogonal representations of the inputs that explain the most variance [28]. The second transformation calculates the number of mismatches between each pair of aligned genome sequences, also known as the Hamming distance, to create a distance matrix. Most phylogenetic methods begin by building a distance matrix for all sequences in a given multiple sequence alignment. Dimensionality reduction algorithms such as multidimensional scaling (MDS) [29], t-distributed stochastic neighbor embedding (t-SNE) [30], and uniform manifold approximation and projection (UMAP) [31] accept such distance matrices as an input and produce a corresponding low-dimensional representation or “embedding” of those data. Both types of transformation allow us to reduce high-dimensional genome alignments (*M ⇥ N* values for *M* genomes of length *N*) to low-dimensional embeddings where clustering algorithms and visualization are more tractable. Additionally, distance-based methods can reflect the presence or absence of insertions and deletions in an alignment that many phylogenetic methods ignore.

Each of the embedding methods mentioned above has been applied previously to genomic data to visualize relationships between individuals and identify clusters of related genomes. Although PCA is a generic linear algebra algorithm that optimizes for an orthogonal embedding of the data, the principal components from single nucleotide polymorphisms (SNPs) represent mean coalescent times and therefore recapitulate broad phylogenetic relationships [26]. PCA has been applied to SNPs of human genomes [32, 33, 26, 34] and to multiple sequence alignments of viral genomes [35]. MDS attempts to embed input data into a lower-dimensional representation such that each pair of data points are as close in the embedding as they are in the original high-dimensional space. MDS has been applied to multiple gene segments of seasonal influenza viruses to visualize evolutionary relationships between segments [27] and to individual influenza gene segments to reveal low-dimensional trajectories of genetic clusters [36, 37]. Both t-SNE and UMAP build on manifold learning methods like MDS to find low-dimensional embeddings of data that place similar points close together and dissimilar points far apart [38]. These methods have been applied to SNPs from human genomes [39] and single-cell transcriptomes [40, 41].

Although these methods are commonly used for qualitative studies of evolutionary relationships, few studies have attempted to quantify patterns observed in the resulting embeddings, investigate the value of applying these methods to viruses that reassort or recombine, or identify optimal method parameters for application to viruses. Recent studies disagree about whether methods like PCA, t-SNE, and UMAP produce meaningful global structures [38] or arbitrary patterns that distort high-dimensional relationships [42]. To address these open questions, we tuned and validated the performance of PCA, MDS, t-SNE, and UMAP with genomes from simulated influenza-like and coronavirus-like populations and then applied these methods to natural populations of seasonal influenza virus A/H3N2 and SARS-CoV-2. These natural viruses are highly relevant as major causes of global human mortality, common subjects of real-time genomic epidemiology, and representatives of reassortant and recombinant human pathogens. For each combination of virus and embedding method, we quantified the relationship between pairwise genetic and Euclidean embedding distances, identified clusters of closely-related genomes in embedding space, and evaluated the accuracy of clusters compared to genetic groups defined by experts and clusters defined directly from pairwise genetic distances. Finally, we tested the ability of these methods to capture patterns of reassortment between seasonal influenza A/H3N2 hemagglutinin (HA) and neuraminidase (NA) segments and recombination in SARS-CoV-2 genomes. These results demonstrate the interpretability of embeddings from each method and inform our recommendations for future applications of these methods to specific problems in genomic epidemiology.

## 2. Results

### 2.1. The ability of embedding methods to produce global structures for simulated viral populations varies little across method parameters

To understand how well PCA, MDS, t-SNE, and UMAP could represent genetic relationships between samples of human pathogenic viruses under well-defined evolutionary conditions, we simulated influenza-like and coronavirus-like populations as previously described [43, 12] and created embeddings for each population across a range of method parameters. For both influenza- and coronavirus-like population types, we simulated five independent replicates for 60 years, filtered out the first 10 years of each population as a burn-in period, and analyzed the remaining 50 years. We simulated influenza-like populations with a mutation rate of 0.00382 substitutions per site per year to match the natural H3N2 HA rate [43], and we sampled 10 HA sequences per week. We simulated coronavirus-like populations with a mutation rate of 0.0008 substitutions per site per year [44] and a recombination rate of 10*^-^*^5^ events per site per year [12]. We sampled 15 full-length coronavirus sequences approximately every two weeks.

We maximized the local and global interpretability of each method’s embeddings by identifying parameters that maximized a linear relationship between genetic distance and Euclidean distance in low-dimensional space (see Methods). Specifically, we selected parameters that minimized the median of the mean absolute error (MAE) between observed pairwise genetic distances of simulated genomes and predicted genetic distances for those genomes based on their Euclidean distances in each embedding. For methods like PCA and MDS where increasing the number of components available to the embedding could lead to overfitting, we selected the maximum number of components beyond which the median MAE did not decrease by more than 1 nucleotide.

For influenza-like populations, the optimal parameters were 2 components for PCA, 3 components for MDS, perplexity of 200 and learning rate of 100 for t-SNE, and nearest neighbors of 100 and minimum distance of 0.1 for UMAP. As expected, increasing the number of components for PCA and MDS gradually decreased the median MAEs of their embeddings (Supplementary Fig. S1 A and B). However, beyond 2 and 3 components, respectively, the reduction in error did not exceed 1 nucleotide. This result suggests that there were diminishing returns for the increased complexity of additional components. Both t-SNE and UMAP embeddings produced a wide range of errors (the majority between 10 and 20 average mismatches) across all parameter values (Supplementary Fig. S1 C and D). Embeddings from t-SNE appeared robust to variation in parameters, with a slight improvement in median MAE associated with perplexity of 200 and little benefit to any of the learning rate values (Supplementary Fig. S1 C). Similarly, UMAP embeddings were robust across the range of tested parameters, with the greatest benefit coming from setting the nearest neighbors greater than 25 and no benefit from changing the minimum distance between points (Supplementary Fig. S1 D).

The optimal parameters for coronavirus-like populations were similar to those for the influenza-like populations. The optimal parameters were 2 components for PCA, 3 for MDS, perplexity of 100 and learning rate of 100 for t-SNE, and nearest neighbors of 50 and minimum distance of 0.05 for UMAP. As with influenza-like populations, both PCA and MDS showed diminishing benefits of increasing the number of components (Supplementary Fig. S2 A and B). Similarly, we observed little improvement in MAEs from varying t-SNE and UMAP parameters (Supplementary Fig. S2 C and D). The most noticeable improvement came from setting t-SNE’s perplexity to 100 (Supplementary Fig. S2 C). These results indicate the limits of t-SNE and UMAP to represent global genetic structure from these data.

We inspected representative embeddings based on the optimal parameters above for the first four years of influenza- and coronavirus-like populations. Simulated sequences from the same time period tended to map closer in embedding space, indicating the maintenance of “local” genetic structure in the embeddings (Fig. 1). Most embeddings also represented some form of global structure, with later generations mapping closer to intermediate generations than earlier generations. MDS maintained the greatest continuity between generations for both population types (Supplementary Fig. S3). In contrast, PCA, t-SNE, and UMAP all demonstrated tighter clusters of samples separated by potentially arbitrary space. These qualitative results matched our expectations based on how well each method maximized a linear relationship between genetic and Euclidean distances during parameter optimization (Supplementary Fig. S1 and Supplementary Fig. S2).

**Fig. 1.**
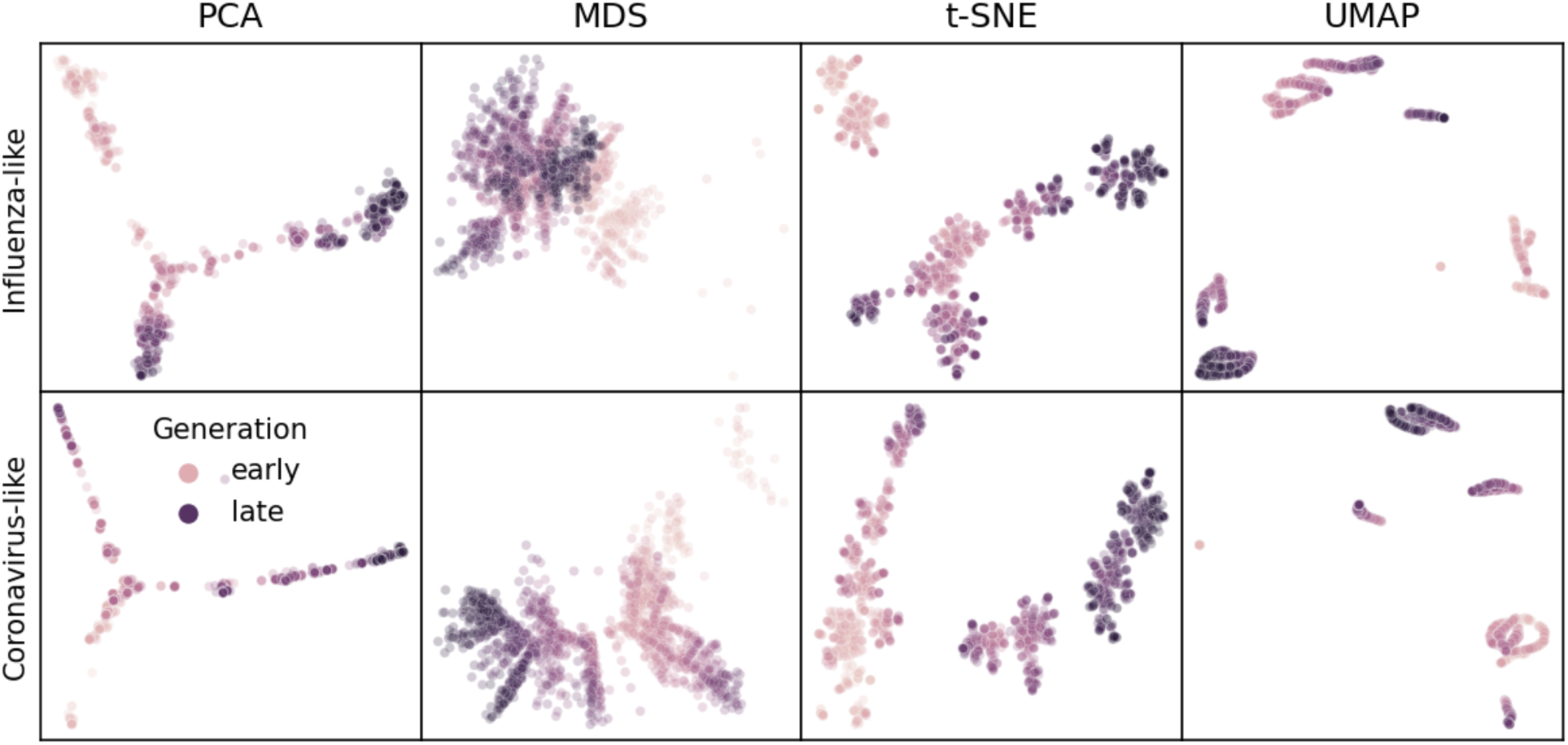
Representative embeddings for simulated populations using optimal parameters per pathogen (rows) and embedding method (columns). Each panel shows the embedding for sequences from the first four years of a single replicate population for the corresponding pathogen type. Each point represents a simulated viral sequence colored by its generation with darker values representing later generations. Supplementary Fig. S3 shows the full MDS embedding for all components.

### 2.2. Embedding clusters recapitulate phylogenetic clades for seasonal influenza H3N2

Seasonal influenza H3N2’s hemagglutinin (HA) sequences provide an ideal positive control to test embedding methods and clustering in low-dimensional space. H3N2’s HA protein evolves rapidly, accumulating amino acid mutations that enable escape from adaptive immunity in human populations [45]. These mutations produce distinct phylogenetic clades that represent potentially different antigenic phenotypes. The World Health Organization (WHO) Global Influenza Surveillance and Response System regularly sequences genomes of circulating influenza lineages [46] and submits these sequences to public INSDC databases like NCBI’s GenBank [47]. These factors, coupled with HA’s relatively short gene size of 1,701 nucleotides, facilitate real-time genomic epidemiology of H3N2 [48] and rapid analysis by the embedding methods we wanted to evaluate. We analyzed H3N2 HA sequences from two consecutive time periods including an “early” dataset from 2016–2018 and a “late” dataset from 2018–2020. For each dataset, we created embeddings with all four methods, identified clusters in the embeddings with HDBSCAN, and calculated the accuracy of clusters relative to expert-defined genetic groups (see Methods). As a point of comparison, we also identified HDBSCAN clusters from pairwise genetic distances between sequences and compared the accuracy of these clusters to clusters from embeddings and known genetic groups. We used the early dataset to identify cluster parameters that minimized the distance between clusters and known genetic groups. We tested these optimal parameters with the late dataset. This approach allowed us to maximize cluster accuracy against the background of embedding method parameters that we already optimized to maximize the interpretability of visualizations.

We first applied each embedding method to the early H3N2 HA sequences (2016–2018) and compared the placement of these sequences in the embeddings to their corresponding clades in the phylogeny. All four embedding methods qualitatively recapitulated the 10 Nextstrain clades observed in the phylogeny (Fig 2 and Supplementary Fig. S4). Samples from the same clade generally grouped tightly together. Most embedding methods also delineated larger phylogenetic clades, placing clades A1, A2, A3, A4, and 3c3.A into separate locations in the embeddings. Despite maintaining local and broader global structure, not all embeddings captured intermediate genetic structure. For example, all methods placed A1b and its descendant clades, A1b/135K and A1b/135N, into tight clusters together. The t-SNE embedding created separate clusters for each of these clades, but these clusters all placed so close together in the embedding space that, without previously defined clade labels, we would have visually grouped these samples into a single cluster. These results qualitatively replicate the patterns we observed in embeddings for simulated influenza-like populations where genetically similar sequences placed closer together and all methods except MDS produced tight local clusters of similar sequences (Fig 1).

**Fig. 2.**
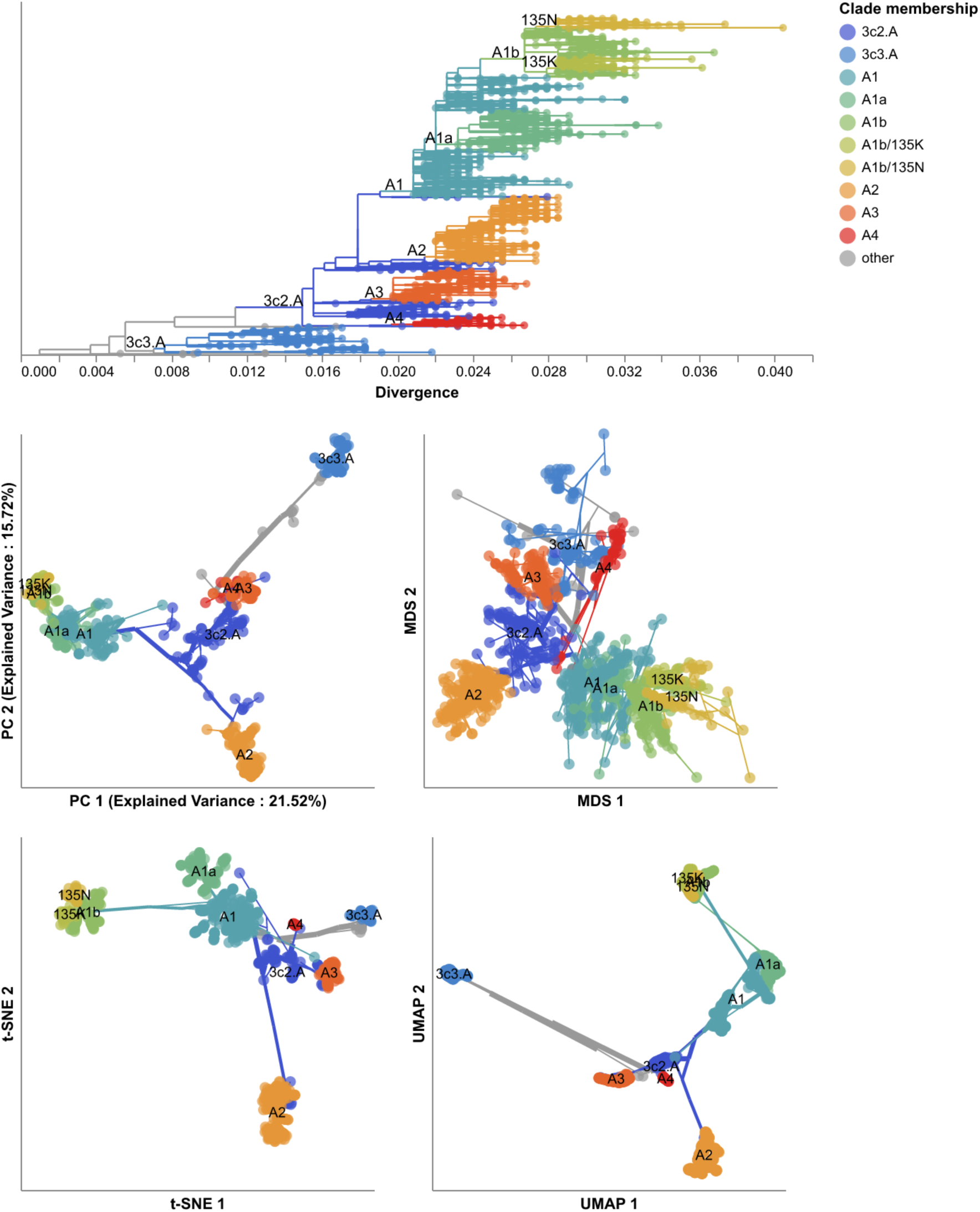
Phylogeny of early (2016–2018) influenza H3N2 HA sequences plotted by nucleotide substitutions per site on the x-axis (top) and low-dimensional embeddings of the same sequences by PCA (middle left), MDS (middle right), t-SNE (bottom left), and UMAP (bottom right). Tips in the tree and embeddings are colored by their Nextstrain clade assignment. Line segments in each embedding reflect phylogenetic relationships with internal node positions calculated from the mean positions of their immediate descendants in each dimension (see Methods). Line colors represent the clade membership of the most ancestral node in the pair of nodes connected by the segment. Line thickness in the embeddings scales by the square root of the number of leaves descending from a given node in the phylogeny. Clade labels appear in the tree at the earliest ancestral node of the tree for each clade. Clade labels appear in each embedding at the average position on the x and y axis for sequences in a given clade.

To quantify the apparent maintenance of local and global structure by all four embedding methods, we calculated the relationship between pairwise genetic and Euclidean distance of samples in each embedding. All methods maintained a linear pairwise relationship for samples that differed by no more than *⇡*10 nucleotides (Fig 3). Only MDS consistently maintained that linearity as genetic distance increased (Pearson’s *R*^2^ = 0.94). We observed a less linear relationship for samples with more genetic differences in PCA (Pearson’s *R*^2^ = 0.75), t-SNE (Pearson’s *R*^2^ = 0.38), and UMAP (Pearson’s *R*^2^ = 0.52) embeddings. While PCA and UMAP Euclidean distances increased monotonically with genetic distance, t-SNE embeddings placed some pairs of samples with intermediate distances of 35-45 nucleotides closer together than pairs of samples with lower genetic distances.

**Fig. 3.**
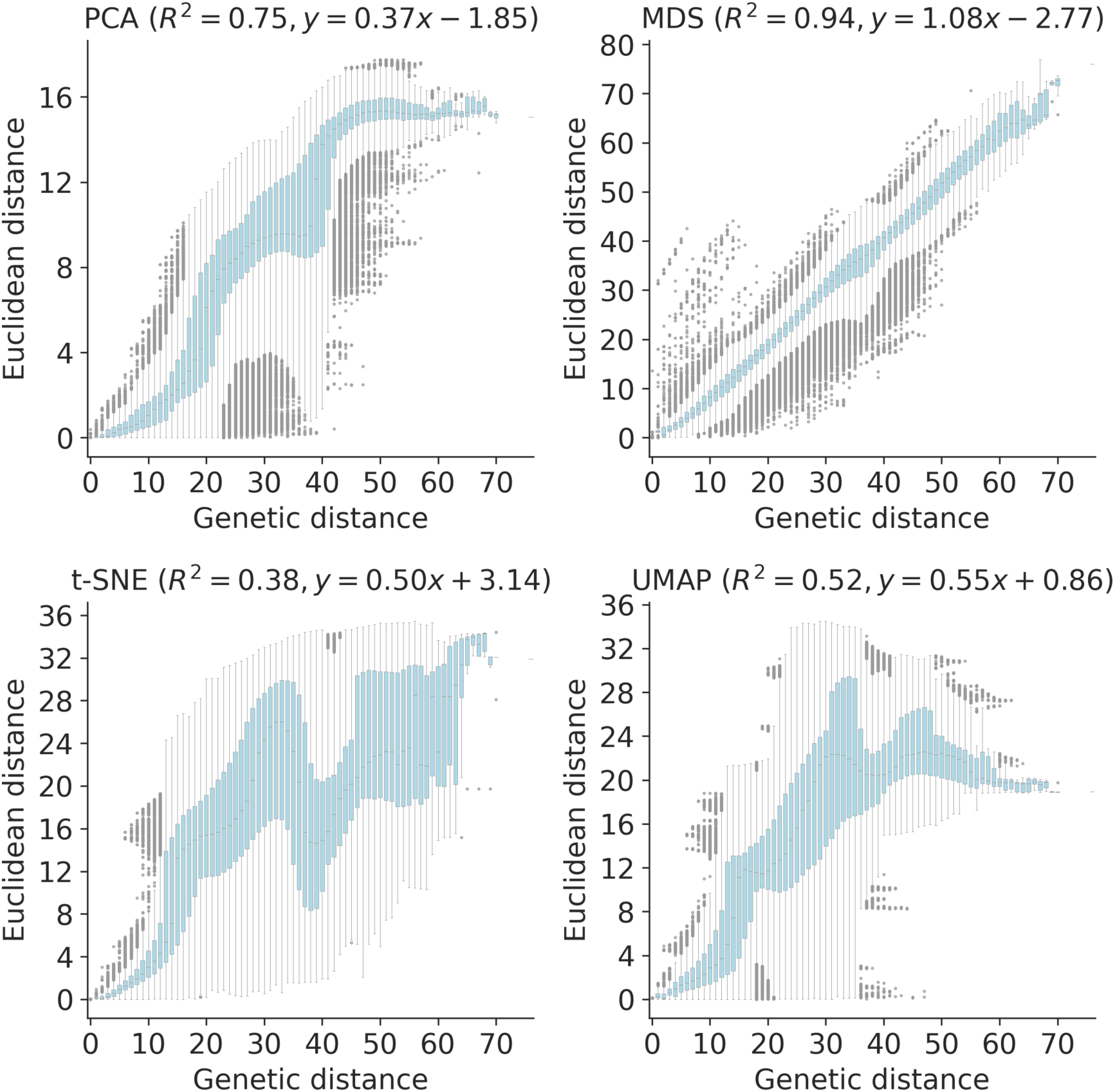
Relationship between pairwise genetic and Euclidean distances in embeddings of early (2016–2018) influenza H3N2 HA sequences by PCA (upper left), MDS (upper right), t-SNE (lower left), and UMAP (lower right). Each boxplot represents the distribution of pairwise Euclidean distances at a given genetic distance. Panel titles include Pearson’s *R*^2^ values and linear regression coefficients between the plotted distances.

Next, we found clusters for early H3N2 HA embeddings and pairwise genetic distances and calculated the distance from these clusters to previously defined genetic groups. We assigned cluster labels to each sample with the hierarchical clustering algorithm, HDBSCAN [49]. We calculated distances between clusters and known genetic groups with the normalized variation of information (VI) metric [50] which produces a value of 0 for identical groups and 1 for maximally different groups (see Methods).

HDBSCAN does not require an expected number of clusters as input, but it does provide a parameter for the minimum distance required between clusters. We optimized this minimum distance threshold by minimizing the VI distance between known genetic groups and clusters produced with different threshold values (Supplementary Table S1). Clusters from all four embedding methods captured broad phylogenetic clades (A1, A1b, A2, A3, 3c2.A, and 3c3.A) but failed to distinguish between A1b and its descendants (Fig 4). The 15 clusters from t-SNE and 7 from UMAP most accurately captured expert clade assignments (normalized VI=0.09), followed by PCA’s 7 clusters (normalized VI=0.10) and MDS’s 9 clusters (normalized VI=0.11). PCA and UMAP clusters failed to distinguish between A4 and its ancestral clade of 3c2.A. Although all methods produced clusters that were generally supported by cluster-specific mutations (Supplementary Table S3 and Supplementary Fig. S5), only MDS and t-SNE produced monophyletic clusters (Supplementary Table S2). The average pairwise genetic diversity within and between clusters matched the diversity within Nextstrain clades (Supplementary Fig. S6). These results indicate that all embedding methods could potentially be well-suited for clustering and classification of H3N2 HA sequences. Clusters based on genetic distances were the farthest from Nextstrain clades (normalized VI=0.17, Supplementary Table S1) and had higher average genetic diversity than clusters from embedding methods (Supplementary Fig. S6). These results suggest that applying dimensionality reduction methods prior to clustering could improve cluster accuracy.

**Fig. 4.**
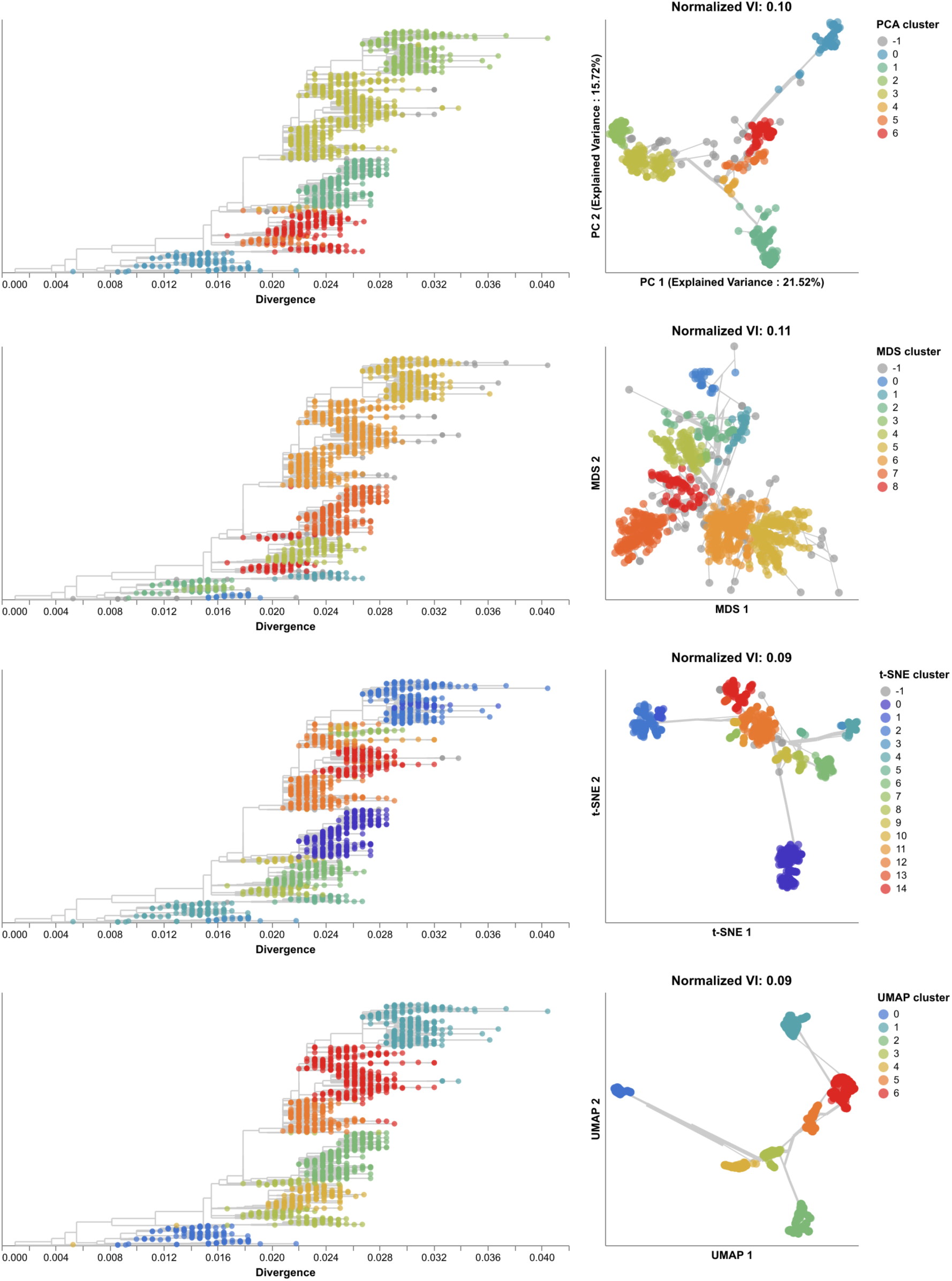
Phylogenetic trees (left) and embeddings (right) of early (2016–2018) influenza H3N2 HA sequences colored by HDBSCAN cluster. Normalized VI values per embedding reflect the distance between clusters and known genetic groups (Nextstrain clades). Line segments in each embedding reflect phylogenetic relationships with internal node positions calculated from the mean positions of their immediate descendants in each dimension (see Methods). Line thickness in the embeddings scales by the square root of the number of leaves descending from a given node in the phylogeny.

To understand whether these embedding methods and optimal cluster parameters could effectively cluster previously unseen sequences, we applied each method to the late H3N2 HA dataset (2018–2020), identified clusters per embedding, and calculated the VI distance between clusters and previously defined clades. The late dataset included 9 clades with at least 10 samples (Supplementary Fig. S7). These clades had a greater average between-clade distance than clades in the early dataset (Supplementary Fig. S6). Clusters from PCA (N=5), MDS (N=6), t-SNE (N=7), and UMAP (N=8) were similarly accurate, with normalized VI distances of 0.07, 0.08, 0.05, and 0.09, respectively (Fig. 5, Supplementary Fig. S8, and Supplementary Table S1). Genetic distance clusters (N=4) were farthest from Nextstrain clades (normalized VI=0.12, Supplementary Table 1). MDS split clade A3’s 17 samples into two widely separated groups in its Euclidean space. On further inspection of this clade in the tree, we found 7 homoplasies (mutations that also occur elsewhere in the tree) on the branch leading to A3 and 10 homoplasies on the branch leading to one of A3’s subclades. Previous work with MDS embeddings of HA sequences has shown that MDS’s global optimization algorithm is sensitive to homoplasies [37]. In this dataset, clade A3 represented an extreme example where MDS could not cluster a clade that had many homoplasies and few samples. In contrast, PCA, t-SNE, and UMAP correctly clustered A3 samples together in their embeddings, showing the robustness of these methods to homoplasies. Accordingly, clusters from all methods except MDS were monophyletic (Supplementary Table S2). The majority of clusters from all methods were supported by cluster-specific mutations (Supplementary Table S3 and Supplementary Fig. S5). The average of pairwise nucleotide differences within clusters generally matched the diversity within Nextstrain clades (Supplementary Fig. S6). As with the early H3N2 HA dataset, clusters from genetic distances between late H3N2 HA sequences had the highest within and between group pairwise nucleotide differences.

**Fig. 5.**
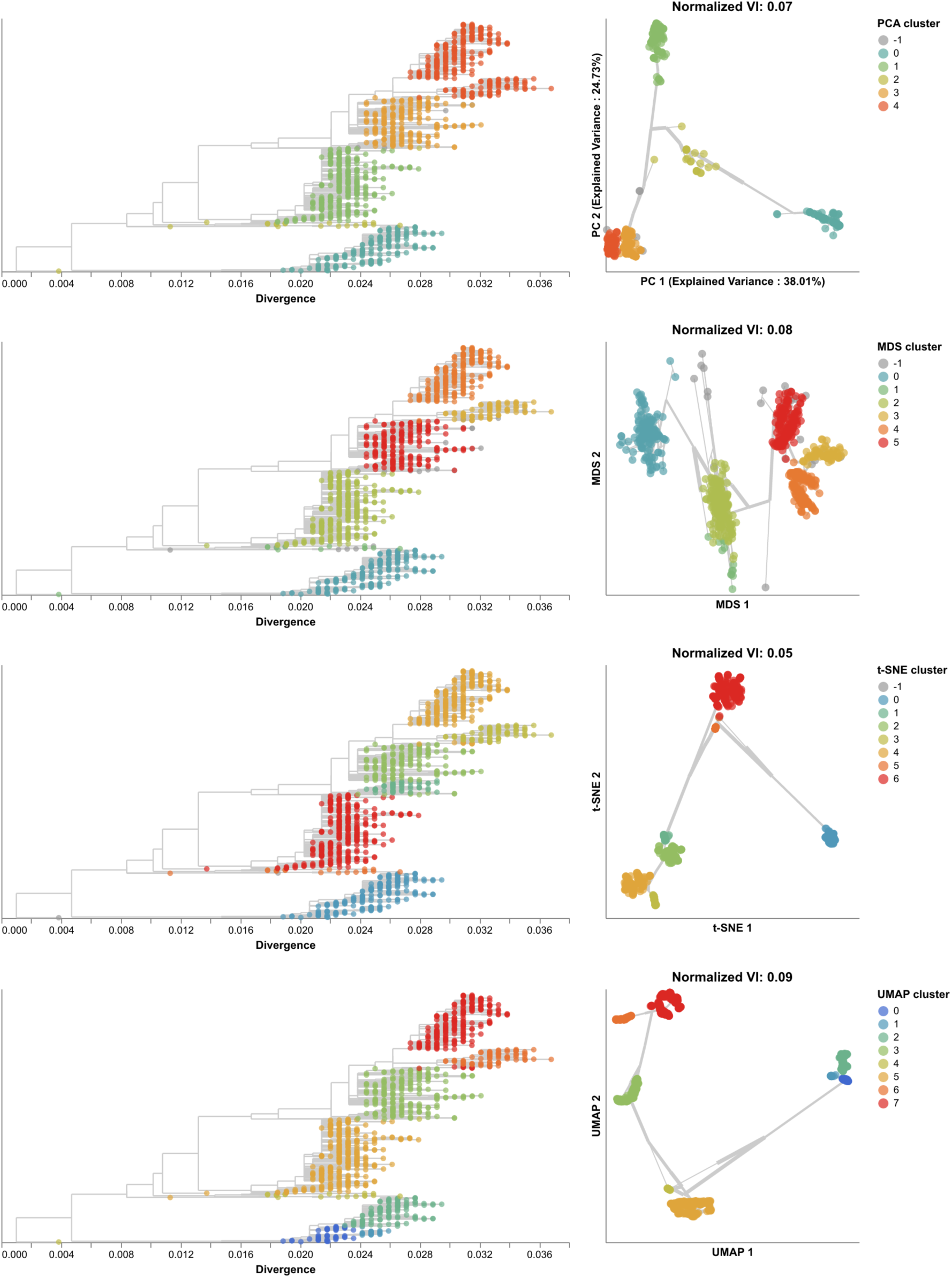
Phylogenetic trees (left) and embeddings (right) of late (2018–2020) H3N2 HA sequences colored by HDBSCAN cluster. Normalized VI values per embedding reflect the distance between clusters and known genetic groups (Nextstrain clades). Line segments in each embedding reflect phylogenetic relationships with internal node positions calculated from the mean positions of their immediate descendants in each dimension (see Methods). Line thickness in the embeddings scales by the square root of the number of leaves descending from a given node in the phylogeny.

Cluster accuracies were robust to changes in sampling density under the same even geographic and temporal sampling scheme, with MDS clusters producing the lowest median distance to Nextstrain clades (Supplementary Fig. S9A). However, biased sampling toward the USA and clade 3c3.A decreased cluster accuracy for t-SNE and UMAP (Supplementary Fig. S9B). Under even sampling, genetic distance clusters were less accurate than clusters from embeddings until the number of input sequences exceeded 1500. However, genetic distance clusters were consistently accurate under biased sampling. These results show that all four methods can produce clusters that accurately capture known genetic groups when applied to previously unseen H3N2 HA samples with unbiased sampling. Clusters from PCA, MDS, and genetic distances are better choices when the composition of sequences is biased strongly by geography or time.

### 2.3. Joint embeddings of hemagglutinin and neuraminidase sequences identify seasonal influenza virus H3N2 reassortment events

Given that clusters from embedding methods could recapitulate expert-defined clades, we measured how well the same methods could capture reassortment events between multiple gene segments as detected by biologically-informed computational models. Evolution of HA and NA surface proteins contributes to the ability of influenza viruses to escape existing immunity [45] and HA and NA genes frequently reassort [5, 6, 51]. Therefore, we focused our reassortment analysis on HA and NA sequences, sampling 1,607 viruses collected between January 2016 and January 2018 with sequences for both genes. We inferred HA and NA phylogenies from these sequences and applied TreeKnit to both trees to identify maximally compatible clades (MCCs) that represent reassortment events [11]. Of the 208 reassortment events identified by TreeKnit, 15 (7%) contained at least 10 samples representing 1,049 samples (65%, Supplementary Fig. S10).

We created PCA, MDS, t-SNE, and UMAP embeddings from the HA alignments and from merged HA and NA alignments. We identified clusters in both HA-only and HA/NA embeddings and calculated the VI distance between these clusters and the MCCs identified by TreeKnit. We expected that clusters from HA-only embeddings could only reflect reassortment events when the HA clade involved in reassortment happened to carry characteristic nucleotide mutations. We expected that the VI distances for clusters from HA/NA embeddings would improve on the baseline distances calculated with the HA-only clusters.

All embedding methods produced more accurate clusters from the HA/NA alignments than the HA-only alignments (Fig. 6, Supplementary Fig. S11, and Supplementary Table S1). HA/NA clusters from t-SNE were closest to reassortment events identified by TreeKnit (normalized VI=0.06) and almost twice as close to those reassortment events than the HA-only clusters (normalized VI=0.11). For the other three embedding methods, including NA with HA reduced the distance between clusters and reassortment events by a similar absolute values of 0.05, 0.07, and 0.03 for PCA, MDS, and UMAP, respectively. Clusters from genetic distances improved the most by the addition of NA from a normalized VI of 0.2 to 0.11 (Supplementary Table 1). Embeddings with both genes produced more clusters for all methods than the HA-only embeddings with 3 additional clusters in PCA (Supplementary Fig. S12), 9 in MDS (Supplementary Fig. S13), 2 in t-SNE (Supplementary Fig. S14 and Supplementary Fig. S15), 1 in UMAP (Supplementary Fig. S16), and 16 in genetic distance clusters (Supplementary Table 1). All embeddings of HA/NA alignments produced distinct clusters for the known reassortment event within clade A2 [51] as represented by MCCs 14 and 11. Other pairs of larger reassortment events that occurred in the same part of the HA tree like MCCs 9 and 12 or MCCs 5 and 10 mapped farther apart in all HA/NA embeddings compared to HA-only embeddings (Supplementary Fig. S11). We noted that some of the additional clusters in HA/NA embeddings likely also reflected genetic diversity in NA that was independent of reassortment between HA and NA. These results suggest that a single embedding of multiple gene segments could identify biologically meaningful clusters within and between all genes.

**Fig. 6.**
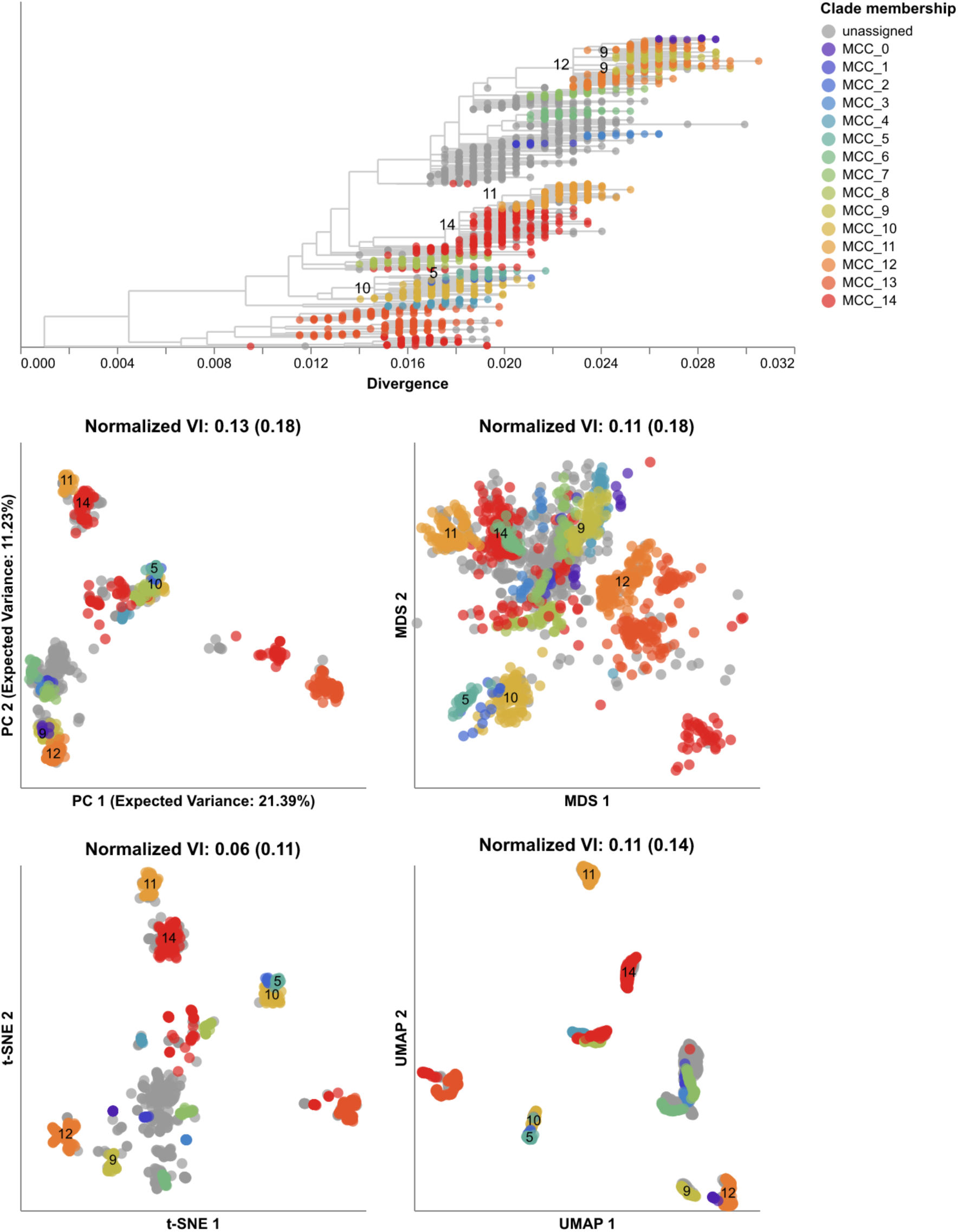
Phylogeny of early (2016–2018) influenza H3N2 HA sequences plotted by nucleotide substitutions per site on the x-axis (top) and low-dimensional embeddings of the same HA sequences concatenated with matching NA sequences by PCA (middle left), MDS (middle right), t-SNE (bottom left), and UMAP (bottom right). Tips in the tree and embeddings are colored by their TreeKnit Maximally Compatible Clades (MCCs) label which represents putative HA/NA reassortment groups. Tips from MCCs with fewer than 10 sequences are colored as “unassigned”. The first normalized VI values per embedding reflect the distance between HA/NA clusters and known genetic groups (MCCs). VI values in parentheses reflect the distance between HA-only clusters and known genetic groups. MCC labels appear in the tree and each embedding for larger pairs of reassortment events. MCC 9 represents two Nextstrain clades, so its labels appear twice in the tree. MCCs 14 and 11 represent a previously published reassortment event within Nextstrain clade A2 [51]. Labels for MCC 14 represent the subset of its sequences from clade A2.

### 2.4. SARS-CoV-2 clusters recapitulate broad genetic groups corresponding to Nextstrain clades

SARS-CoV-2 poses a greater challenge to embedding methods than seasonal influenza, with an unsegmented genome an order of magnitude longer than influenza’s HA or NA [52], a mutation rate in the spike surface protein subunit S1 that is four times higher than influenza H3N2’s HA rate [53], and increasingly common recombination [54, 55]. However, multiple expert-based clade definitions exist for SARS-CoV-2, enabling comparison between clusters from embeddings and known genetic groups. These definitions span from broad genetic groups named by the WHO as “variants of concern” (e.g., “Alpha”, “Beta”, etc.) [56] or systematically defined by the Nextstrain team [57, 58, 59] to smaller, emerging genetic clusters defined by Pango curators [19]. As with seasonal influenza, we defined an early SARS-CoV-2 dataset spanning from January 2020 to January 2022, embedded genomes with the same four methods, and identified HDBSCAN clustering parameters that minimized the VI distance between embedding clusters and previously defined genetic groups as defined by Nextstrain clades and Pango lineages (see Methods). To test these optimal cluster parameters, we produced clusters from embeddings of a late SARS-CoV-2 dataset spanning from January 2022 to November 2023 and calculated the VI distance between those clusters and known genetic groups. Unlike the seasonal influenza analysis, we counted insertion and deletion (“indel”) events in pairwise genetic distances for SARS-CoV-2, to improve the resolution of distance-based embeddings.

All embedding methods placed samples from the same Nextstrain clades closer together and closely related Nextstrain clades near each other (Fig. 7). For example, the most genetically distinct clades like 21J (Delta) and 21L (Omicron) placed farthest from other clades, while both Delta clades (21I and 21J) placed close together (Fig. 7, Supplementary Fig. S17). MDS placed related clades closer together on a continuous scale, while PCA, t-SNE, and UMAP produced more clearly separate groups of samples. We did not observe the same clear grouping of Pango lineages. For example, the Nextstrain clade 21J (Delta) contained 11 Pango lineages that all appeared to map into the same overlapping space in all four embeddings (Supplementary Fig. S18). These results suggest that distance-based embedding methods can recapitulate broader genetic groups of SARS-CoV-2, but that these methods lack the resolution of finer groups defined by Pango nomenclature.

**Fig. 7.**
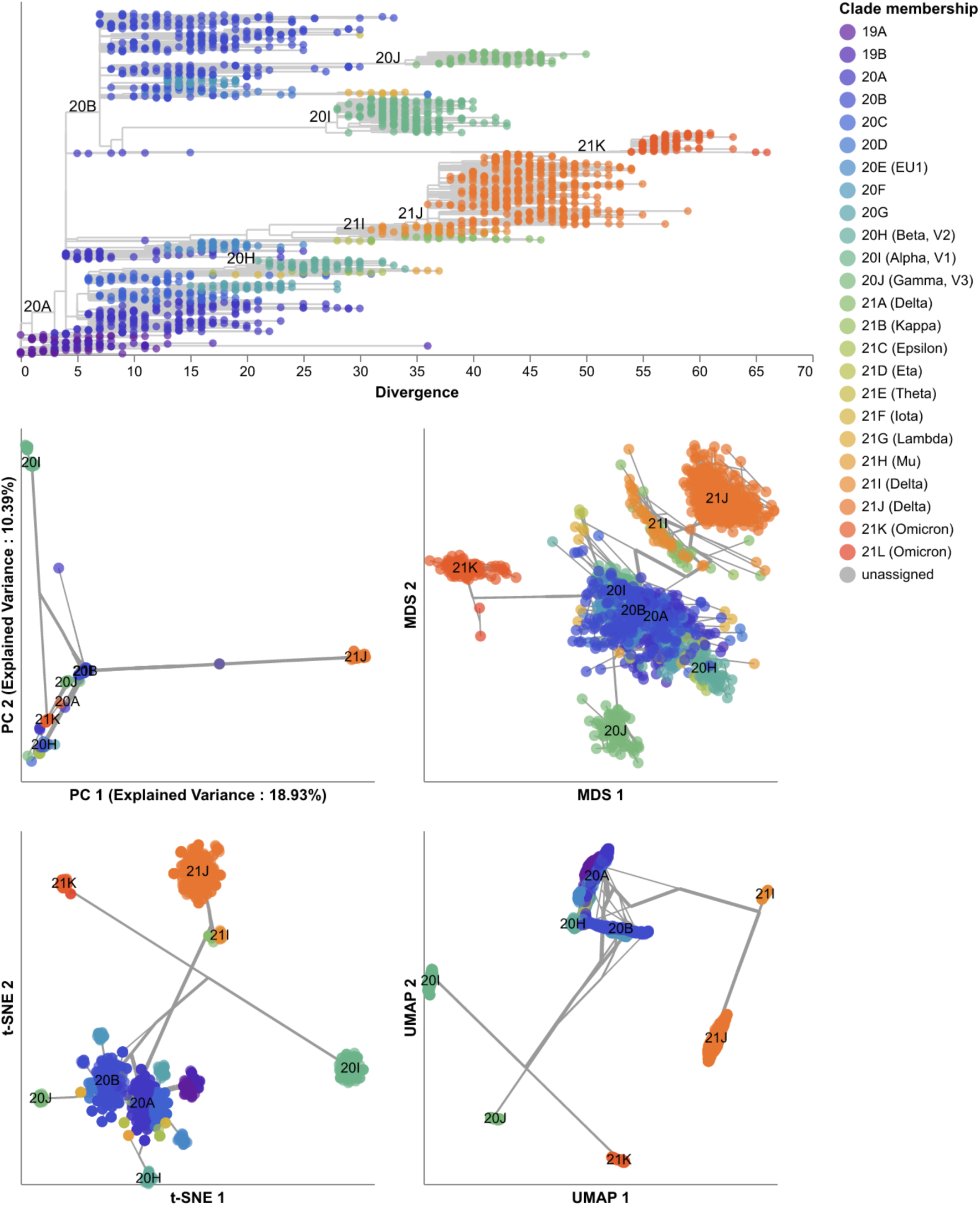
Phylogeny of early (2020–2022) SARS-CoV-2 sequences plotted by number of nucleotide substitutions from the most recent common ancestor on the x-axis (top) and low-dimensional embeddings of the same sequences by PCA (middle left), MDS (middle right), t-SNE (bottom left), and UMAP (bottom right). Tips in the tree and embeddings are colored by their Nextstrain clade assignment. Line segments in each embedding reflect phylogenetic relationships with internal node positions calculated from the mean positions of their immediate descendants in each dimension (see Methods). Line thickness in the embeddings scales by the square root of the number of leaves descending from a given node in the phylogeny. Clade labels in the tree and embeddings highlight larger clades.

We quantified the maintenance of local and global structure in early SARS-CoV-2 embeddings by fitting a linear model between pairwise genetic and Euclidean distances of samples. PCA produced the weakest linear relationship (Pearson’s *R*^2^ = 0.35, Fig. 8). MDS created a strong linear mapping across the range of observed genetic distances (Pearson’s *R*^2^ = 0.92). Both t-SNE and UMAP maintained intermediate degrees of linearity (Pearson’s *R*^2^ = 0.55 and *R*^2^ = 0.59, respectively). These embeddings placed the most genetically similar samples close together and the most genetically distinct farther apart. However, these embeddings did not consistently place pairs of samples with intermediate genetic distances at an intermediate distance in Euclidean space. The linear relationship for genetically similar samples in t-SNE and UMAP remained consistent up to a genetic distance of approximately 30 nucleotides.

**Fig. 8.**
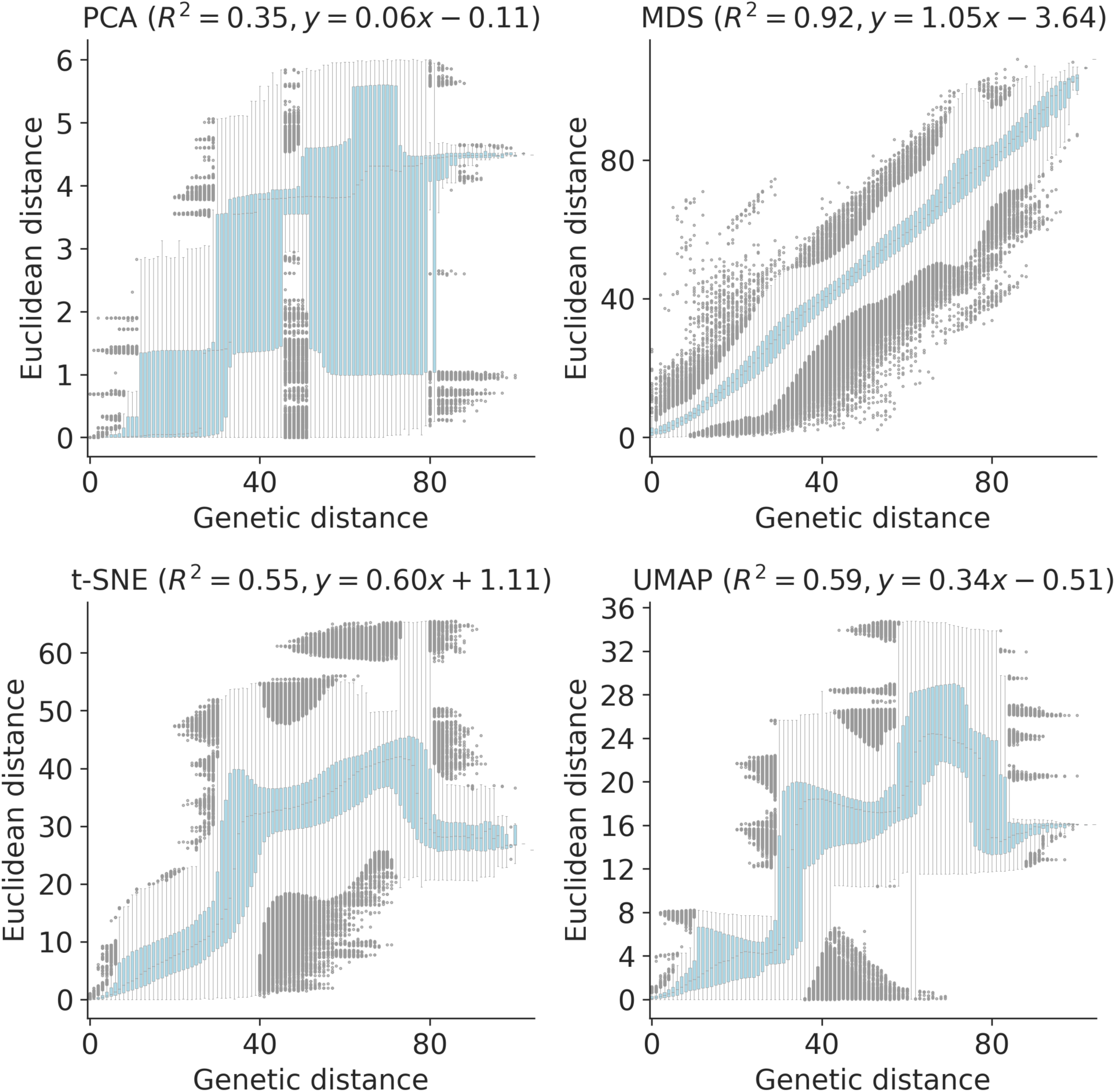
Relationship between pairwise genetic and Euclidean distances in embeddings for early (2020–2022) SARS-CoV-2 sequences by PCA (upper left), MDS (upper right), t-SNE (lower left), and UMAP (lower right). Each boxplot represents the distribution of pairwise Euclidean distances at a given genetic distance. Panel titles include Pearson’s *R*^2^ values and linear regression coefficients between the plotted distances.

We identified clusters in embeddings and pairwise genetic distances from early SARS-CoV-2 data using cluster parameters that minimized the normalized VI distance between clusters and known genetic groups. Since Nextstrain clades and Pango lineages represented different resolutions of genetic diversity, we identified optimal distance thresholds per lineage definition. However, we found that the optimal thresholds were the same for both lineage definitions, except for genetic distance clusters which had a slightly higher optimal threshold for Pango lineages than Nextstrain clades (Supplementary Table S1). The 19 clusters from t-SNE were closest to the 24 Nextstrain clades (normalized VI=0.09) followed by MDS, UMAP, genetic distances, and PCA with distances of 0.15, 0.16, 0.17, and 0.23, respectively (Fig. 9 and Supplementary Table S1). Clusters from PCA, t-SNE, and UMAP all represented completely monophyletic groups (Supplementary Table S2). Clusters from all methods were generally supported by cluster-specific mutations including 2 of 3 (67%) PCA clusters, 9 of 16 (56%) MDS clusters, 15 of 19 (79%) t-SNE clusters, 5 of 6 (83%) UMAP clusters, and 7 of 9 (78%) genetic distance clusters (Supplementary Table S3 and Supplementary Fig. S5). Clusters from t-SNE had the most similar within-group distances to Nextstrain clades (Supplementary Fig. S19).

**Fig. 9.**
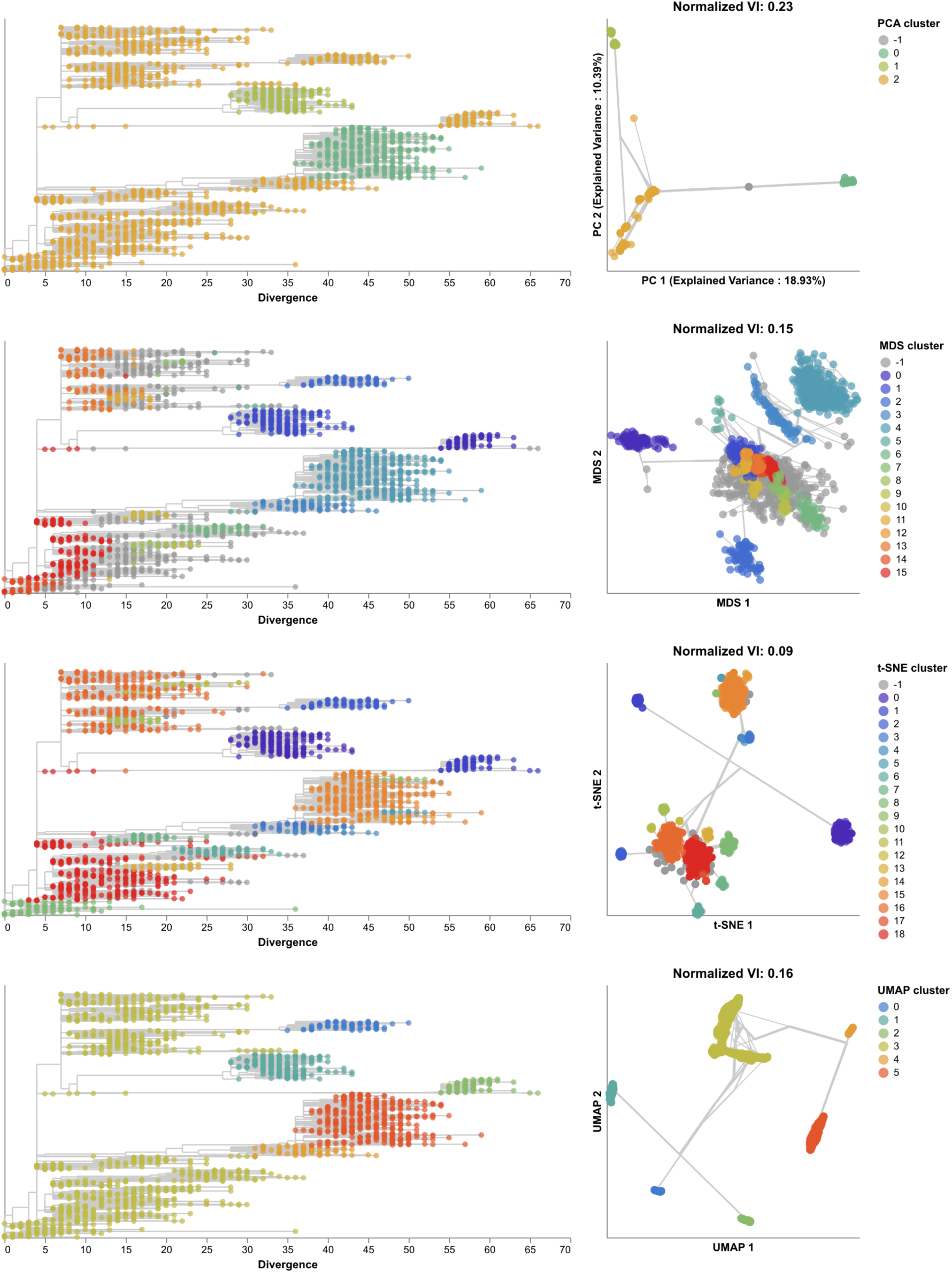
Phylogenetic trees (left) and embeddings (right) of early (2020–2022) SARS-CoV-2 sequences colored by HDBSCAN cluster. Normalized VI values per embedding reflect the distance between clusters and known genetic groups (Nextstrain clades). Line segments in each embedding reflect phylogenetic relationships with internal node positions calculated from the mean positions of their immediate descendants in each dimension (see Methods). Line thickness in the embeddings scales by the square root of the number of leaves descending from a given node in the phylogeny.

Clusters from all methods were farther from the 35 Pango lineages (Supplementary Fig. S20 and Supplementary Table S1). The 19 clusters from t-SNE were the closest (normalized VI=0.14) followed by MDS, UMAP, genetic distances, and PCA with distances of 0.23, 0.25, 0.26, and 0.32, respectively (Supplementary Table S1). We found that within-cluster distances for t-SNE were lower on average than within-lineage Pango distances, suggesting that Pango lineages were not as tightly scoped as we originally expected (Supplementary Fig. S19). These results confirm quantitatively that these embeddings methods can accurately capture broader genetic diversity of SARS-CoV-2, but most methods cannot distinguish between fine resolution genetic groups defined by Pango lineage nomenclature.

To test the optimal cluster parameters identified above, we applied embedding methods to late SARS-CoV-2 data and compared clusters from these embeddings to the corresponding Nextstrain clades and Pango lineages. Compared to the 17 Nextstrain clades defined in this time period (Supplementary Fig. S21), the closest clusters were from t-SNE (normalized VI=0.09 with 66 clusters) and UMAP (normalized VI=0.09 with 13 clusters, Fig. 10 and Supplementary Table S1). We attributed t-SNE’s additional clusters to recombinant lineages that were genetically distinct but which received a generic “recombinant” label in Nextstrain’s clade definitions instead of a unique clade name (Supplementary Fig. S21). Although we did not consider these non-monophyletic recombinant samples when calculating VI distances between clusters and Nextstrain clades, these samples appear in each embedding where they could form their own distinct clusters. Only t-SNE, UMAP, and genetic distance clusters were fully monophyletic (Supplementary Table S2). Genetic distance, PCA, and t-SNE clusters were best supported by cluster-specific mutations with 16 of 17 clusters (94%), 6 of 7 clusters (86%), and 51 of 66 clusters (77%), respectively (Supplementary Table S3 and Supplementary Fig. S5). Clusters from t-SNE had the lowest average within-group distances (Supplementary Fig. S19). We observed similar absolute and relative distances to Nextstrain clades across methods at different sampling densities under an even geographic and temporal sampling scheme with t-SNE and UMAP consistently producing the most accurate clusters (Supplementary Fig. S23A). In the presence of geographic and genetic bias associated with randomly sampling the late SARS-CoV-2 data, UMAP produced the most accurate clusters across all sampling densities while t-SNE clusters became less accurate as the number of biased sequences increased (Supplementary Fig. S23B). In contrast to the same analysis for H3N2 HA populations, clusters from genetic distances were consistently farther from Nextstrain clades than clusters from embeddings across all sampling densities and biases.

**Fig. 10.**
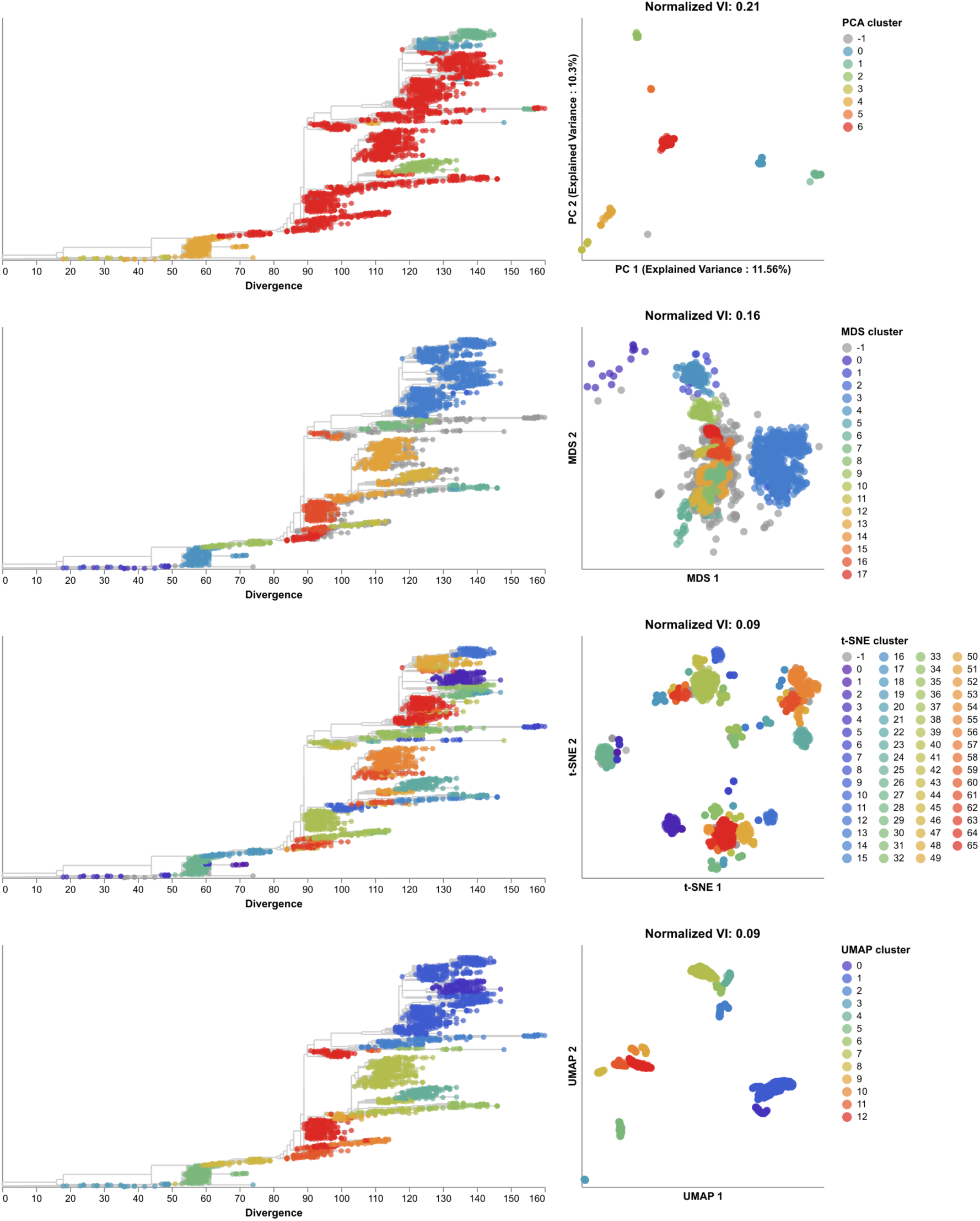
Phylogenetic trees (left) and embeddings (right) of late (2022–2023) SARS-CoV-2 sequences colored by HDBSCAN cluster. Normalized VI values per embedding reflect the distance between clusters and known genetic groups (Nextstrain clades).

All methods produced less accurate representations of the 137 Pango lineages (Supplementary Figs. S22 and S24 and Supplementary Table 1). However, t-SNE clusters were nearly as accurate with a normalized VI of 0.14, suggesting that t-SNE’s numerous additional clusters likely did represent many of the recombinant Pango lineages in the dataset that all received a “recombinant” Nextstrain clade label. Of the 80 recombinant Pango lineages that also had a t-SNE cluster, 79 (99%) mapped to a single t-SNE cluster (Supplementary Fig. S25). Of the 52 t-SNE clusters with recombinant Pango lineages, 43 (83%) mapped to a single Pango lineage. Clusters from other methods were at least twice as far from Pango lineages as t-SNE’s clusters, suggesting that these other methods poorly captured recombinant lineages. With the exception of t-SNE’s performance, these results replicate the patterns we observed with early SARS-CoV-2 data where clusters from embeddings more effectively represented broader genetic diversity than the finer resolution diversity denoted by Pango lineages. Unlike the Pango lineages in the early SARS-CoV-2 data, the lineages from the later data exhibited fewer pairwise genetic distances between samples in each lineage than samples in Nextstrain clades or any embedding cluster (Supplementary Fig. S19).

To understand whether t-SNE clusters could capture Pango-resolution genetic groups within a single Nextstrain clade, we evenly sampled approximately 2,000 sequences from a dominant Nextstrain clade with many Pango lineages, 21J (Delta), and identified clusters from a t-SNE embedding of those data. Within the 1,992 sequences sampled from 21J (Delta), we found 38 Pango lineages after collapsing lineages with fewer than 10 sequences into their parent lineages. We found 28 t-SNE clusters representing 1,806 sequences (91%) with 186 sequences (9%) assigned to the unclustered “-1” label (Supplementary Fig. S26). The VI distance between Pango lineages and all clusters (including the unclustered group) was 0.17 (Supplementary Table 1). This distance was consistent with the distance of 0.14 between Pango lineages and t-SNE clusters from both the full early and late SARS-CoV-2 datasets. The VI distance between Pango lineages and clusters without the unclustered sequences was 0.13, confirming that one quarter of the distance between t-SNE clusters and Pango lineages above came from unclustered sequences. Of the 38 Pango lineages with a t-SNE cluster, 30 lineages (79%) had a single corresponding t-SNE cluster, seven lineages (18%) had two or three t-SNE clusters, and one lineage (B.1.617.2) had five t-SNE clusters (Supplementary Fig. S26). Of the 28 t-SNE clusters, 21 clusters (75%) had a single corresponding Pango lineage, 6 (21%) mapped to two or three Pango lineages, and 1 (cluster 27) mapped to 18 Pango lineages with most sequences from B.1.617.2 and AY.4. These results suggest that clusters from t-SNE embeddings can capture more Pango-resolution genetic groups by analyzing sequences within a specific Nextstrain clade.

### 2.5. Distance-based embeddings reflect SARS-CoV-2 recombination events

Finally, we tested the ability of sequence embeddings to place known recombinant lineages of SARS-CoV-2 between their parental lineages in Euclidean space. We reasoned that each recombinant lineage, *X*, should always place closer to its parental lineages *A* and *B* than the parental lineages place to each other. Based on this logic, we calculated the average Euclidean distance between pairs of samples in lineages *A* and *B*, *A* and *X*, and *B* and *X* for each embedding method (see Methods). We identified recombinant lineages that mapped closer to both of their parental lineages and those that mapped closer to at least one of the parental lineages. We identified 66 recombinant lineages for which that lineage and both of its parental lineages had at least 10 genomes (Supplementary Table S4). MDS embeddings most consistently placed recombinant lineages between the parental lineages with correct placement of 60 (91%) lineages (e.g., Supplementary Fig. S27). The t-SNE, UMAP, and PCA embeddings correctly placed 55 (83%), 50 (76%), and 45 (68%) lineages, respectively. Additionally, all 66 recombinant lineages placed closer to at least one parent in all embeddings except for one lineage in the PCA embeddings.

## 3. Discussion

### 3.1. Tree-free dimensionality reduction methods can provide valuable biological insights

We applied four standard dimensionality reduction methods to simulated and natural genome sequences of two relevant human pathogenic viruses and found that the resulting embeddings could reflect pairwise genetic relationships between samples and capture previously identified genetic groups. From our analysis of simulated influenza- and coronavirus-like sequences, we found that each method produced consistent embeddings of genetic sequences for two distinct pathogens, 50 years of evolution, and a wide range of practical method parameters. Of the four methods, MDS most accurately reflected pairwise genetic distances between simulated samples in its embeddings. From our analysis of natural populations of seasonal influenza H3N2 HA and SARS-CoV-2 sequences, we confirmed that MDS most reliably reflected pairwise genetic distances. We found that clusters from t-SNE embeddings most accurately recapitulated previously defined genetic groups at the resolution of WHO variants and Nextstrain clades and consistently produced clusters that corresponded to monophyletic groups in phylogenies and were robust to the presence of homoplasies. Clusters from t-SNE embeddings of H3N2 HA and NA sequences most accurately matched reassortment clades identified by a biologically-informed model based on ancestral reassortment graphs. MDS embeddings consistently placed known recombinant lineages of SARS-CoV-2 between their parental lineages, while t-SNE clusters most accurately captured recombinant lineages. All of the embedding methods and the HDBSCAN clustering method rely on pairwise comparisons between all samples making them robust to individual outliers caused by sequencing errors. Furthermore, distance-based methods like MDS, t-SNE, and UMAP easily ignore missing characters in individual sequences. These results show that tree-free dimensionality reduction methods can provide valuable biological insights for human pathogenic viruses through easily interpretable visualizations of genetic relationships and the ability to account for genetic variation that tree-based phylogenetic methods cannot use, including indels, reassortment, and recombination.

### 3.2. Recommendations for application of methods to new pathogens

From these results, we can also make the following recommendations about how to apply these methods to other viral pathogens. First, evenly sample the available genome sequences across time and geography, to minimize bias in embeddings. Then, choose which embedding method to use based on the question under investigation. For analyses that require the most accurate low-dimensional representation of pairwise genetic distances across local and global scales, use MDS with 3 dimensions. For analyses that need to find clusters of closely related samples, use t-SNE with a perplexity of 100 (or less, if using fewer than 100 samples) and a learning rate that scales with the number of samples in the data. In all cases, plot the relationship between pairwise genetic distances and Euclidean distances in each embedding. These plots reveal the range of genetic distances that an embedding can represent linearly and act as a sanity check akin to plotting the temporal signal present in samples prior to inferring a time-scaled phylogeny [60, 4]. Before finding clusters in the t-SNE embedding, determine the minimum genetic distance desired between clusters, and use the pairwise genetic and Euclidean distance plot to find the corresponding Euclidean distance to use as a threshold for HDBSCAN. While HDBSCAN clusters require this pathogen-specific tuning, the linear relationship between Euclidean and genetic distance remains robust to changes in method parameters.

The computational complexity of the original MDS and t-SNE algorithms scales by the cube and square of the number of input samples (*N*), respectively [61, 30]. However, more efficient alternate implementations exist for both methods that can scale by *N* log *N* and operate on millions of samples [61, 62, 63, 64, 65]. For this reason, the primary practical limitation to scaling MDS and t-SNE to increasingly more pathogen sequences is the time and space required to calculate pairwise genetic distances for all samples. Implementing more efficient pairwise distance calculations remains a future direction for this area of research.

### 3.3. Limitations of methods and analysis

Despite the promise of these simple methods to answer important public health questions about human pathogenic viruses, these methods and our analyses suffer from inherent limitations. The lack of an underlying biological model is both a strength and the clearest limitation of the dimensionality reduction methods we considered here. For example, embeddings of SARS-CoV-2 genomes that represent the complete circulating diversity at a given time cannot capture the same fine-grained genetic resolution as Pango lineage annotations. Only t-SNE clusters of SARS-CoV-2 genomes within a single Nextstrain clade get close to defining Pango-resolution genetic groups. Each method provides only a few parameters to tune its embeddings and these parameters have little effect on the qualitative outcome. PCA is sensitive to missing characters in individual sequences and must treat each gap character from a single deletion event as an independent mutation instead of a single variant. In maintaining a linear relationship between Euclidean and genetic distances, MDS sacrifices the ability to form more accurate genetic clusters for viruses with large genomes like SARS-CoV-2 and struggles to correctly cluster samples from the same genetic group with numerous recurrent mutations. Neither t-SNE nor UMAP maintain a linear relationship between pairwise Euclidean and genetic distances across the observed range of genetic distances. As a result, viewers cannot know that samples mapping far apart in a t-SNE or UMAP embedding are as genetically distant as they appear. Given these limitations of these methods, we do not expect them to replace biologically-informed methods that provide more meaningful parameters to tune their algorithms. Instead, these methods provide an easy first step to produce interpretable visualizations and clusters of genome sequences, prior to analysis with more sophisticated methods with biological models.

We note that our analysis reflects a small subset of human pathogenic viruses and dimensionality reduction methods. We focused on analysis of two respiratory RNA viruses that contribute substantially to seasonal human morbidity and mortality, but numerous alternative pathogens would also have been relevant subjects. For example, HIV represents a canonical example of a highly recombinant and bloodborne virus, while Zika, dengue, and West Nile viruses represent pathogens with multiple host species in a transmission chain. Similarly, we selected only four dimensionality reduction methods from myriad options that are commonly applied to genetic data [66]. We chose these methods based on their wide use and availability in tools like scikit-learn [67] and to limit the dimensionality of our analyses.

### 3.4. Current applications and future directions for applying dimensionality reduction methods to viral pathogens

Following the recommendations we outlined above, researchers can immediately put these methods into practice with viral pathogens. We provide optimal settings for each pathogen and embedding method in this study and open source tools to apply these methods to other pathogens through the `pathogen-embed` toolkit. We provide this toolkit through the standard Python package repository PyPI, Bioconda, and Nextstrain-managed Docker and Conda environments. These tools integrate easily into existing workflows for the genomic epidemiology of viruses and their results can be visualized with Nextstrain. Alternately, researchers may choose to apply similar existing tools developed for analysis of metagenomic or bacterial populations [68, 69, 70, 71, 72] to the analysis of viral populations.

In the short term, these methods can identify biologically-relevant clusters from viral sequence alignments. For example, researchers can apply t-SNE and HDBSCAN clustering to alignments of unsegmented viruses like Zika or Ebola that lack existing clade definitions to identify candidate phylogenetic groups. Similarly, researchers can jointly build embeddings from alignments of segmented viruses like influenza and identify clusters corresponding to putative reassortment groups. For example, the Nextstrain team now routinely produces t-SNE embeddings and clusters from HA and NA sequences for weekly seasonal influenza analyses at https://nextstrain.org/seasonal-flu/ to track reassortment events. Researchers can also quickly apply these methods in response to outbreaks like the recent H5N1 avian influenza outbreak in cattle in the United States [73]. As a proof of concept, we applied t-SNE to all eight gene segments of recent H5N1 sequences, identified clusters with HDBSCAN, and confirmed the previously reported reassortment groups with PB2/NP and the other gene segments in the cattle outbreak. Researchers can easily visualize their embeddings in standard visualization tools for genomic epidemiology including Nextstrain`’`s Auspice [74], MicrobeTrace [14], or MicroReact [13].

Some limitations noted above suggest future directions for this line of research. In the long term, researchers may benefit from analyzing viral genomes with a broader range of dimensionality reduction methods including neural network models [75, 42]. Biologically-informed versions of these methods could support finer-grained cluster identification and more intuitive parameters for users to adjust to suit their pathogens. We also expect that researchers will benefit from applying the methods we describe here to a broader range of virus families including those that lack standard phylogenetic clade definitions or that undergo too much recombination to be appropriately analyzed with standard phylogenetic methods. Finally, the combination of dimensionality reduction methods and clustering with HDBSCAN provides the foundation for future methods to automatically identify reassortant and recombinant lineages. For example, representing viral genomes with low-dimensional MDS embeddings could simplify the problem of identifying recombinant lineages to a matter of classifying groups by their Euclidean distances.

In conclusion, we showed that simple dimensionality reduction methods operating on pairwise genetic differences can capture biologically-relevant clusters of phylogenetic clades, reassortment events, and patterns of recombining lineages for human pathogenic viruses. The conceptual and practical simplicity of these tools should enable researchers and public health practitioners to more readily visualize and compare samples for human pathogenic viruses when phylogenetic methods are either unnecessary or inappropriate.

## 4. Methods

### 4.1. Embedding methods

We selected four standard and common dimensionality reduction (or “embedding”) methods to apply to human pathogenic viruses: PCA, MDS, t-SNE, and UMAP. PCA operates on a matrix with samples in rows, “features” in columns, and numeric values in each cell [28]. To apply PCA to multiple sequence alignments, we encoded each nucleotide value as a vertex in a simplex as previously described [76] with “A” encoded as (1, −1, −1), “C” as (−1, 1, −1), “G” as (−1, −1, 1), and “T” as (1, 1, 1). We encoded all other characters as (0, 0, 0). This encoding places each nucleotide at an equal distance from the other nucleotides in the simplex and produces an input matrix for PCA of size *N ⇥* 3*L* for *N* sequences of length *L*. We applied scikit-learn’s PCA implementation to the resulting numerical matrix with the “full” singular value decomposition solver and 10 components [67]. To minimize the effects of missing data on the PCA embeddings, we dropped all columns with “N” or “-” characters from concatenated H3N2 HA/NA alignments and SARS-CoV-2 alignments prior to producing PCA embeddings.

The remaining three methods operate on a distance matrix. We constructed a distance matrix from a multiple sequence alignment by calculating the pairwise Hamming distance between nucleotide sequences. By default, the Hamming distance only counted mismatches between pairs of standard nucleotide values (A, C, G, and T), ignoring other values including gaps. We implemented an optional mode that additionally counted each occurrence of consecutive gap characters in either input sequence as individual insertion/deletion (“indel”) events.

We applied scikit-learn’s MDS implementation to a given distance matrix, with an option to set the number of components in the resulting embedding [67]. Similarly, we applied scikit-learn’s t-SNE implementation, with options to set the “perplexity” and the “learning rate”. The perplexity controls the number of neighbors the algorithm uses per input sample to determine an optimal embedding [30]. This parameter effectively determines the balance between maintaining “local” or “global” structure in the embedding [41]. The learning rate controls how rapidly the t-SNE algorithm converges on a specific embedding [77, 30] and should scale with the number of input samples [78]. We initialized t-SNE embeddings with the first two components of the corresponding PCA embedding, as previously recommended to obtain more accurate global structure [41, 38]. Finally, we applied the `umap-learn` Python package written by UMAP`’`s authors, with options to set the number of `“`nearest neighbors`”` and the `“`minimum distance`”` [31]. As with t-SNE`’`s perplexity parameter, the nearest neighbors parameter determines how many adjacent samples the UMAP algorithm considers per sample to find an optimal embedding. The minimum distance sets the lower limit for how close any two samples can map next to each other in a UMAP embedding. Lower minimum distances allow tighter groups of samples to form. For both t-SNE and UMAP, we used the default number of components of 2.

### 4.2. Simulation of influenza-like and coronavirus-like populations

Given the relative lack of prior application of dimensionality reduction methods to human pathogenic viruses, we first attempted to understand the behavior and optimal parameter values for these methods when applied to simulated viral populations with well-defined evolutionary parameters. To this end, we simulated populations of influenza-like and coronavirus-like viruses using SANTA-SIM [79]. These simulated populations allowed us to identify optimal parameters for each embedding method, without overfitting to the limited data available for natural viral populations. For each population type described below, we simulated five independent replicates with fixed random seeds for 60 years, filtered out the first 10 years of each population as a burn-in period, and analyzed the remaining years.

We simulated influenza-like populations as previously described with 1,700 bp hemagglutinin sequences [43]. As in that previous study, we scaled the number of simulated generations per real year to 200 per year to match the observed mutation rate for natural H3N2 HA sequences, we sampled 10 genomes every 4 generations for 12,000 generations (or 60 years of real time), and filtered out the first 10 years of each simulation’s data as a burn-in period.

We simulated coronavirus-like populations as previously described for human seasonal coronaviruses with genomes of 21,285 bp [12]. For the current study, we assigned 30 generations per real year to obtain mutation rates similar to the 8 *⇥* 10*^-^*^4^ substitutions per site per year estimated for SARS-CoV-2 [44]. To account for the effect of recombination on optimal method parameters, we simulated populations with a recombination rate of 10*^-^*^5^ events per site per year based on human seasonal coronaviruses for which recombination rates are well-studied [12, 80]. We calibrated the overall recombination probability in SANTA-SIM such that the number of observed recombination events per year matched the expected number for human seasonal coronaviruses (0.3 per year) [12]. To assist with this calibration of recombination events per year, we modified the SANTA-SIM source code to emit a boolean status of “is recombinant” for each sampled genome. This change allowed us to identify recombinant genomes by their metadata in downstream analyses and calculate the number of recombination events observed per year. For each replicate population, we sampled 15 genomes every generation for 1,800 generations (or 60 years of real time) and filtered out the first 10 years of each simulation’s data as a burn-in period.

### 4.3. Optimization of embedding method parameters

We identified optimal parameter values for each embedding method with time series cross-validation of embeddings based on simulated populations [81]. To increase the interpretability of embedding space, we defined parameters as “optimal” when they maximized the linear relationship between pairwise genetic distance of viral genomes and the corresponding Euclidean distance between those same genomes in an embedding. This optimization approach allowed us to also determine the degree to which each method could recapitulate this linear relationship.

For each simulated population replicate, we created 10 training and test datasets that each consisted of 4 years of training data and 4 years of test data preceded by a 1-year gap from the end of the training time period. These settings produced training/test data with 2000 samples each for influenza-like populations and 1800 samples each for coronavirus-like populations. For each combination of training/test dataset, embedding method, and method parameters, we applied the following steps. We created an embedding from the training data with the given parameters, fit a linear model to estimate pairwise genetic distance from pairwise Euclidean distance in the embedding, created an embedding from the test data, estimated the pairwise genetic distance for genomes in the test data based on their Euclidean distances and the linear model fit to the training data, and calculated the mean absolute error (MAE) between estimated and observed genetic distances in the test data. We summarized the error for a given population type, method, and method parameters across all population replicates and training/test data by calculating the median of the MAE. For all method parameters except those controlling the number of components used for the embedding, we selected the optimal parameters as those that minimized the median MAE for a given embedding method. Since increasing the number of components used by PCA and MDS allows these methods to overfit to available data, we selected the optimal number of components for these methods as the number beyond which the median MAE did not decrease by at least 1 nucleotide. This approach follows the same concept from the MDS algorithm itself where optimization occurs iteratively until the algorithm reaches a predefined error threshold [29].

With the approach described above, we tested each method across a range of relevant parameters with all combinations of parameter values. For PCA and MDS, we tested the number of components between 2 and 10. For t-SNE, we tested perplexity values of 15, 30, 100, 200, and 300, and we tested learning rates of 100, 200, and 500. For UMAP, we tested nearest neighbor values of 25, 50, and 100, and we tested values for the minimum distance that points can be in an embedding of 0.05, 0.1, and 0.25.

### 4.4. Selection of natural virus population data

We selected recent publicly available genome sequences and metadata for seasonal influenza H3N2 HA and NA genes and SARS-CoV-2 genomes from INSDC databases [47]. For both viruses, we divided the available data into “early” and “late” datasets to use as training and test data, respectively, for identification of virus-specific clustering parameters.

For analyses that focused only on H3N2 HA data, we defined the early dataset between January 2016 and January 2018 and the late dataset between January 2018 to January 2020. These datasets reflected two years of recent H3N2 evolution up to the time when the SARS-CoV-2 pandemic disrupted seasonal influenza circulation. For both early and late datasets, we evenly sampled 25 sequences per country, year, and month. We excluded outliers which were sequences either labeled as environmental samples, containing over 100 gap characters within the HA sequence, or flagged by TreeTime [4] for having a phylogenetic divergence that exceeded four times the interquartile interval of residuals from a root-to-tip regression for all sequences in the same tree. With this sampling scheme, we selected 1,523 HA sequences for the early dataset and 1,073 for the late dataset. For analyses that combined H3N2 HA and NA data, we defined a single dataset between January 2016 and January 2018, keeping 1,607 samples for which both HA and NA have been sequenced.

For SARS-CoV-2 data, we defined the early dataset between January 1, 2020 and January 1, 2022 and the late dataset between January 1, 2022 and November 3, 2023. For the early dataset, we evenly sampled 1,736 SARS-CoV-2 genomes by geographic region, year, and month, excluding known outliers that had been previously identified by the Nextstrain team during weekly phylogenetic surveillance since January 2020 (https://github.com/nextstrain/ncov/blob/master/defaults/exclude.txt). For the late dataset, we used the same even sampling by space and time to select 1,309 representative genomes. In addition to these genomes, we identified all recombinant lineages in the official Pango designations as of November 3, 2023 (https://github.com/cov-lineages/ `pango-designation/raw/1bf4123/pango_designation/alias_key.json`) for which the recombinant lineage and both of its parental lineages had at least 10 genome records each. We sampled at most 10 genomes per lineage for all distinct recombinant and parental lineages for a total of 1,157. With these additional genomes, the late SARS-CoV-2 dataset included 2,464 total genomes.

### 4.5. Evaluation of linear relationships between genetic distance and Euclidean distance in embeddings

To evaluate the biological interpretability of distances between samples in low-dimensional embeddings, we plotted the pairwise Euclidean distance between samples in each embedding against the corresponding genetic distance between the same samples. We calculated Euclidean distance using all components of the given embedding (e.g., 2 components for PCA, t-SNE, and UMAP and 3 components for MDS). For each embedding, we fit a linear model between Euclidean and genetic distance and calculated the squared Pearson’s correlation coefficient, *R*^2^. The distance plots provide a qualitative assessment of each embedding’s local and global structure relative to a biologically meaningful scale of genetic distance, while the linear models and correlation coefficients quantify the global structure in the embeddings.

### 4.6. Phylogenetic analysis

For each natural population described above, we created an annotated phylogenetic tree. For seasonal influenza H3N2 HA and NA sequences, we aligned sequences with MAAFT (version 7.486) [82, 83] using the `augur align` command (version 22.0.3) [84]. For SARS-CoV-2 sequences, we used existing reference-based alignments provided by the Nextstrain team (https://docs.nextstrain.org/projects/ncov/en/latest/reference/remote inputs.html) and generated with Nextalign (version 2.14.0) [21]. We inferred each phylogeny with IQ-TREE (version 2.1.4-beta) [85] using the `augur tree` command with its default IQ-TREE parameters of `-ninit 2 -n 2 -me 0.05` and a general time reversible (GTR) model. These are the same parameters we use to build SARS-CoV-2 and seasonal influenza phylogenies for https://nextstrain.org. We named internal nodes of the resulting divergence tree with TreeTime (version 0.10.1) [4] using the `augur refine` command. We visualized phylogenies with Auspice [74], after first converting the trees to Auspice JSON format with `augur export`. To visualize phylogenetic relationships in the context of each pathogen embedding, we calculated the mean Euclidean position of each internal node in each dimension of a given embedding (e.g., MDS 1) based on the Euclidean positions of that node`’`s immediate descendants and plotted line segments on the embedding connecting each node of the tree with its immediate parent to represent branches in the phylogeny. We only plotted these phylogenetic relationships on embeddings for pathogen datasets that lacked reassortment and recombination including early and late H3N2 HA and early SARS-CoV-2 datasets.

### 4.7. Definitions of genetic groups by experts or biologically-informed models

We annotated phylogenetic trees with genetic groups previously identified by experts or assigned by biologically-informed models. For seasonal influenza H3N2, the World Health Organization assigns “clade” labels to clades in HA phylogenies that appear to be genetically or phenotypically distinct from other recently circulating H3N2 samples. We used the latest clade definitions for H3N2 maintained by the Nextstrain team as part of their seasonal influenza surveillance efforts [48].

As seasonal influenza clades only account for the HA gene and lack information about reassortment events, we assigned joint HA and NA genetic groups using a biologically-informed model, TreeKnit (version 0.5.6) [11]. TreeKnit infers ancestral reassortment graphs from two gene trees, finding groups of samples for which both genes share the same history. These groups, also known as maximally compatible clades (MCCs), represent samples whose HA and NA genes have evolved together. TreeKnit attempts to resolve polytomies in one tree using information present in the other tree(s). Input trees for TreeKnit must contain the same samples and root on the same sample. Because of these TreeKnit expectations, we inferred HA and NA trees with IQ-TREE with a custom argument to collapse near-zero-length branches (`-czb`). We rooted the resulting trees on the same sample that we used as an alignment reference, A/Beijing/32/1992, and pruned this sample prior to downstream analyses. We applied TreeKnit to the rooted HA and NA trees with a gamma value of 2.0 and the `--better-MCCs` flag, as previously recommended for H3N2 analyses [11]. Finally, we filtered the MCCs identified by TreeKnit to retain only those with at least 10 samples. We also omitted the root MCC which consistently included the most recent common ancestor of both HA and NA trees and did not represent a reassortment event, as described in TreeKnit`’`s definition of MCCs [11].

For SARS-CoV-2, we used both coarser “Nextstrain clades” [57, 58, 59] and more granular Pango lineages [19] provided by Nextclade as “Nextclade pango” annotations. Nextstrain clade definitions represent the World Health Organization’s variants of concern along with post-Omicron phylogenetic clades that have reached minimum global and regional frequencies and growth rates. Pango lineages represent expert-curated lineages (https://github.com/cov-lineages/pango-designation) and must contain at least 5 samples with an unambiguous evolutionary event. Additionally, Pango lineages produced by recombination receive a lineage name prefixed by an “X”, while Nextstrain clades do not explicitly reflect recombination events.

Since Pango lineages can represent much smaller genetic groups than are practically useful, we collapsed lineages with fewer than 10 samples in our analysis into their parental lineages using the pango aliasor tool (https://github.com/corneliusroemer/pangoaliasor). Specifically, we counted the number of samples per lineage, sorted lineages in ascending order by count, and collapsed each lineage with a count less than 10 into its parental lineage in the count-sorted order. This approach allowed small lineages to aggregate with other small parental lineages and meet the 10-sample threshold. We used these “collapsed Nextclade Pango” lineages for subsequent analyses.

### 4.8. Clustering of samples in embeddings

To understand how well embeddings of genetic data could capture previously defined genetic groups, we applied an unsupervised clustering algorithm, HDBSCAN [49], to each embedding. In addition to the four embedding methods, we identified HDBSCAN clusters from a fifth “method” of precomputed pairwise genetic distances. This genetic distance method allowed us to understand how clusters differed between low- and high-dimensional inputs. HDBSCAN identifies initial clusters from high-density regions in the input space and merges these clusters hierarchically. This algorithm allowed us to avoid defining an arbitrary or biased expected number of clusters *a priori*. HDBSCAN provides parameters to tune the minimum number of samples required to seed an initial cluster (“min samples”), the minimum size for a final cluster (“min size”), and the minimum distance between initial clusters below which those clusters are hierarchically merged (“distance threshold”). We hardcoded the min samples to 5 to minimize the number of spurious initial clusters and min size to 10 to reflect our interest in genetic groups with at least 10 samples throughout our analyses. HDBSCAN calculates the distance between clusters on the Euclidean scale of each embedding or on the scale of precomputed nucleotides differences from each genetic distance matrix. To account for variation in method-specific distances, we performed a coarse grid search of distance threshold values for each virus type and method.

We performed the grid search on the early datasets for both seasonal influenza H3N2 HA and SARS-CoV-2. For each dataset and method, we applied HDBSCAN clustering with a distance threshold between 0 and 20 inclusive with steps of 0.5 between values. For a given threshold, we obtained sets of samples assigned to HDBSCAN clusters from the embedding. We evaluated the accuracy of these clusters with variation of information (VI) which calculates the distance between two sets of clusters of the same samples [50]. When two sets of clusters are identical, VI equals 0. When the sets are maximally different, VI is log *N* where *N* is the total number of samples. To make VI values comparable across datasets, we normalized each value by dividing by log *N*, following the pattern used to validate TreeKnit’s MCCs [11]. Unlike other standard metrics like accuracy, sensitivity, or specificity, VI distances do not favor methods that tend to produce more, smaller clusters. For each virus dataset and method, we identified the distance threshold that minimized the normalized VI between HDBSCAN clusters and genetic groups defined by experts or biologically-informed models (“Nextstrain clade” for seasonal influenza and both “Nextstrain clade” and “Pango lineage” for SARS-CoV-2). HDBSCAN allows samples to not belong to a cluster and assigns these samples a numeric label of −1. We intentionally included all unassigned samples in the normalized VI calculation thereby penalizing cluster parameters that increased the number of unassigned samples by increasing their VI values. Since Nextstrain clade assignments could include non-monophyletic labels like “unassigned” and “recombinant” to represent samples that did not map into a single clade, we ignored these labels in our VI distance calculations to avoid rewarding clusters that placed such non-monophyletic samples into the same group. Finally, we used these optimal distance thresholds to identify clusters in out-of-sample data from the late datasets for both viruses and calculate the normalized VI between those clusters and previously defined genetic groups.

### 4.9. Evaluating robustness of embedding cluster accuracy

The cluster accuracies we estimated for late H3N2 HA and SARS-CoV-2 datasets represented a single VI measurement for a single pathogen dataset. To understand how robust these accuracies were across different datasets, we generated alternate random samples from both late pathogen datasets using two different sampling schemes and a range of total sequences sampled. Specifically, we sampled 500, 1000, 1500, 2000, or 2500 total sequences for five replicates per pathogen (random seeds of 0, 1, 2, 3, and 4) with either even sampling by geography and time or random sampling. For the relatively smaller influenza data, we evenly sampled by country, year, and month. For the larger SARS-CoV-2 data, we evenly sampled by region, year, and month. Even sampling attempted to minimize geographic and temporal biases in the original data. Random sampling uniformly selected samples in a way that reflected the bias in the data. Influenza data were heavily biased toward samples from the USA and clade 3c3.A, while SARS-CoV-2 data were biased toward Europe and North America and Nextstrain clades 21K, 21L, and 22B. For each replicate from each sampling scheme and total number of sequences, we embedded the corresponding sequences with each method, identified clusters in embeddings, and calculated the VI distance between those clusters and Nextstrain clade assignments. We plotted the distribution of the resulting VI distances, to estimate the robustness of these values to sampling bias and density (Supplementary Figs. S9 and S23).

### 4.10. Evaluating the monophyletic nature of embedding clusters

To quantify the degree to which embedding clusters represented monophyletic groups in a pathogen phylogeny, we counted the number of times clusters from each embedding method appeared in different parts of the tree. Specifically, we applied `augur traits` with TreeTime (version 0.10.1) [4, 84] to infer cluster labels for internal nodes of the phylogeny for each pathogen dataset and embedding method. Using a preorder traversal of the tree, we identified each transition between different cluster labels assigned to pairs of ancestral and derived internal nodes. Since the `“`unclustered`”` cluster label of `“`-1`”` produced by HDBSCAN could occur in both ancestral and derived nodes and lead to overcounting transitions, we only logged transitions with this label in the ancestral state (e.g., transition from cluster −1 to cluster 0 but not cluster 0 to cluster −1). For each embedding, we counted the number of distinct clusters, total transitions, and excess transitions beyond the expected single transition between pairs of clusters. Embeddings with no excess transitions between clusters represented monophyletic groups.

### 4.11. Identification of cluster-specific mutations

To better understand the genetic basis of embedding clusters, we identified cluster-specific mutations for all HDBSCAN clusters. First, we found all mutations between each sample’s sequence and the reference sequence used to produce the alignment, considering only A, C, G, T, and gap characters. Within each cluster, we identified mutations that occurred in at least 10 samples and in at least 50% of samples in the cluster. We recorded the resulting mutations per cluster in a table with columns for the embedding method, the position of the mutation, the derived allele of the mutation, and a list of the distinct clusters the mutation appeared in. From this table, we could identify mutations the only occurred in specific clusters and mutations that distinguished sets of clusters from each other.

### 4.12. Assessment of HA/NA reassortment in seasonal influenza populations

To assess the ability of embedding methods to detect reassortment in seasonal influenza populations, we applied each method to either HA alignments only or concatenated alignments of HA and NA sequences from the same samples, performed HDBSCAN clustering with the optimal distance threshold for the given method, and calculated the normalized VI between the resulting clusters and TreeKnit MCCs. As mentioned above, we dropped all columns with “N” or “-” characters from the HA and HA/NA alignments prior to producing PCA embeddings. We used the original alignments to calculate distance matrices for all other methods, since distance-based methods can ignore N characters in pairwise comparisons. We compared normalized VI values for the HA-only clusters of each method to the corresponding VI values for the HA/NA clusters. Lower VI values in the HA/NA clusters than HA-only clusters indicated better clustering of samples into known reassortment groups.

### 4.13. Assessment of recombination in SARS-CoV-2 populations

To assess the ability of embedding methods to detect recombination in late SARS-CoV-2 populations (2022-2023), we calculated the Euclidean distances in low-dimensional space between the 10 known recombinant lineages and their respective parental lineages described in “Selection of natural virus population data” above. Given that we optimized each method’s parameters to maximize a linear relationship between genetic and Euclidean distance, we expected embeddings to place recombinant lineages between their parental lineages, reflecting the intermediate genetic state of the recombinants. For a recombinant lineage *X* and its parental lineages *A* and *B*, we calculated the average pairwise Euclidean distance, *D*, between samples in *A* and *B*, *A* and *X*, and *B* and *X*. We identified lineages that mapped properly as those for which *D*(*A, X*) *< D*(*A, B*) and *D*(*B, X*) *< D*(*A, B*). We also identified lineages for which the recombinant lineage placed closer to at least one parent than the distance between the parents. Note that we used the original uncollapsed Pango annotations to identify samples in each lineage, as these were the lineage names used to include recombinant samples in the analysis and define known relationships between recombinant and parental lineages.

## Data availability

The entire workflow for our analyses was implemented with Snakemake [86]. We have provided all source code, configuration files, and datasets at https://github.com/blab/cartography. Interactive phylogenetic trees and corresponding embeddings for natural populations are available at https://nextstrain.org/groups/blab/ under the “cartography” keyword. The `pathogen-embed` Python package, available at https://pypi.org/project/pathogen-embed/, provides command line utilities to calculate distance matrices (`pathogen-distance`), calculate embeddings per method (`pathogen-embed`), apply hierarchical clustering to embeddings (`pathogen-cluster`), and find mutations from alignments associated with clusters (`pathogen-cluster-mutations`). Supplementary data are published on Zenodo at https://doi.org/10.5281/zenodo.13381647.

## Author contributions statement

A.B., T.B., and J.H. conceived the experiments. S.N. and J.H. wrote the software, conducted the experiments, analyzed the results, and wrote the manuscript. A.B. and T.B. reviewed and edited the manuscript.

## Acknowledgments

We thank James Hadfield, Katie Kistler, Maya Lewinsohn, Nicola Muller, Louise Moncla, Nidia Trovao, and Michael Zeller for constructive feedback on this project. We gratefully acknowledge the originating and submitting laboratories of seasonal influenza and SARS-CoV-2 sequences from INSDC databases without whom this work would not be possible (Supplementary Table S5).

