## Supplemental Table S1 and Figures for "Dimensionality reduction distills complex evolutionary relationships in seasonal influenza and SARS-CoV-2"

### Supplementary data

**Supplementary Table S 1. Distances between clusters identified per pathogen dataset and method compared to known genetic groups.** Distances are measured by normalized variation of information (VI). Smaller VI values indicate fewer differences between HDBSCAN clusters and known genetic groups. VI of 0 indicates identical clusters and 1 indicates maximally different clusters. Known genetic groups include Nextstrain clades, reassortment groups identified by TreeKnit as Maximally Compatible Clades (MCC), and Pango lineages. Total clusters refers to the number of clusters identified for a given dataset and method not including the "-1" label that HDBSCAN assigns to records that could not be assigned to a cluster. Threshold refers to the minimum distance between initial clusters for HDBSCAN to consider them as distinct clusters. For embedding methods, the threshold represents the Euclidean distance between sequences in each embedding. For the genetic distance method, the threshold represents the number of nucleotide differences between sequences. We identified optimal thresholds per pathogen, genetic group, and method from early influenza and SARS-CoV-2 data and applied these optimal thresholds to the corresponding late datasets for each pathogen. Datasets and methods without a threshold value in this table used the threshold from their corresponding early datasets. Rows appear in ascending order of VI values per method within each combination of pathogen and genetic group type.

| Pathogen Dataset | Genetic Group Type | Method | Total clusters | Variation of Information (VI) | Threshold |
|---|---|---|---|---|---|
| early influenza H3N2 HA (2016-2018) | Nextstrain clade | t-SNE | 15 | 0.09 | 1.0 |
| | | UMAP | 7 | 0.09 | 1.0 |
| | | PCA | 7 | 0.10 | 0.5 |
| | | MDS | 9 | 0.11 | 3.5 |
| | | genetic | 8 | 0.17 | 8.0 |
| late influenza H3N2 HA (2018-2020) | Nextstrain clade | t-SNE | 7 | 0.05 | |
| | | PCA | 5 | 0.07 | |
| | | MDS | 6 | 0.08 | |
| | | UMAP | 8 | 0.09 | |
| | | genetic | 4 | 0.12 | |
| influenza H3N2 reassortment (HA only) | MCC | t-SNE | 15 | 0.11 | |
| | | UMAP | 7 | 0.14 | |
| | | PCA | 6 | 0.18 | |
| | | MDS | 8 | 0.18 | |
| | | genetic | 8 | 0.20 | |
| influenza H3N2 reassortment (HA and NA) | MCC | t-SNE | 17 | 0.06 | |
| | | MDS | 17 | 0.11 | |
| | | UMAP | 8 | 0.11 | |
| | | genetic | 24 | 0.11 | |
| | | PCA | 9 | 0.13 | |
| early SARS-CoV-2 (2020-2022) | Nextstrain clade | t-SNE | 19 | 0.09 | 1.0 |
| | | MDS | 16 | 0.15 | 0.0 |
| | | UMAP | 6 | 0.16 | 0.5 |
| | | genetic | 10 | 0.17 | 15.0 |
| | | PCA | 3 | 0.23 | 0.5 |
| early SARS-CoV-2 (2020-2022) | Pango | t-SNE | 19 | 0.14 | 1.0 |
| | | MDS | 16 | 0.23 | 0.0 |
| | | UMAP | 6 | 0.25 | 0.5 |
| | | genetic | 8 | 0.26 | 15.5 |
| | | PCA | 3 | 0.32 | 0.5 |
| late SARS-CoV-2 (2022-2023) | Nextstrain clade | t-SNE | 66 | 0.09 | |
| | | UMAP | 13 | 0.09 | |
| | | MDS | 18 | 0.16 | |
| | | PCA | 7 | 0.21 | |
| | | genetic | 16 | 0.21 | |
| late SARS-CoV-2 (2022-2023) | Pango | t-SNE | 66 | 0.14 | |
| | | UMAP | 13 | 0.30 | |
| | | MDS | 18 | 0.37 | |
| | | genetic | 15 | 0.43 | |
| | | PCA | 7 | 0.46 | |
| SARS-CoV-2 21J (Delta) only | Pango | t-SNE | 28 | 0.17 | |

**Supplementary Table S 2. Number of clusters (*n_clusters*) identified by HDBSCAN, transitions between clusters in the phylogeny, and excess transitions indicating non-monophyletic groups per pathogen dataset and embedding method.** This table reports the list of specific clusters per dataset and method (*clusters*), the number of transitions between different cluster labels on the phylogeny (*n_cluster_transitions*), and the list of observed transitions for pairs of clusters (*transitions*). The number of excess transitions between clusters (*n_extra_transitions*) reflects the number of times that we observed a transition from one source cluster to another beyond the one expected transition. Embeddings without any excess transitions reflect monophyletic groups in the corresponding pathogen phylogeny. Data available at `https://zenodo.org/records/13381647/files/S2_Table.csv`.

**Supplementary Table S 3. Mutations observed per embedding cluster or Nextstrain clade relative to a reference genome sequence for each pathogen.** Each row reflects the alternate allele identified at a specific position of a given pathogen genome or gene sequence, the pathogen dataset, the genetic group (an embedding cluster or Nextstrain clade), the number of clusters in the genetic group with the observed mutation, and the list of distinct genetic group labels with the mutation. Mutations must have occurred in at least 10 samples of the given dataset with an allele frequency of at least 50% to be reported in the table. Cluster- or clade-specific mutations appear in rows with a *cluster_count* value of 1. Data available at `https://zenodo.org/records/13381647/files/S3_Table.csv`.

**Supplementary Table S 4. Average Euclidean distances between each known recombinant, $X$, and its parental lineages $A$ and $B$ per embedding method.** Distances include average pairwise comparisons between $A$ and $B$ (*distance_A_B*), $A$ and $X$ (*distance_A_X*), and $B$ and $X$ (*distance_B_X*). Additional columns indicate whether each recombinant lineage maps closer to both parental lineages (or at least one) than those parents map to each other (*X_maps_closer_to_both_parentals* and *X_maps_closer_to_any_parental*, respectively). Records with values of “True” in the column *X_maps_closer_to_both_parentals* represent the expected placement of the recombinant lineage between its two parental lineages. Data available at `https://zenodo.org/records/13381647/files/S4_Table.csv`.

**Supplementary Table S 5. Accessions and authors from originating and submitting laboratories of seasonal influenza and SARS-CoV-2 sequences from INSDC databases.** Data available at `https://zenodo.org/records/13381647/files/S5_Table.tsv`.

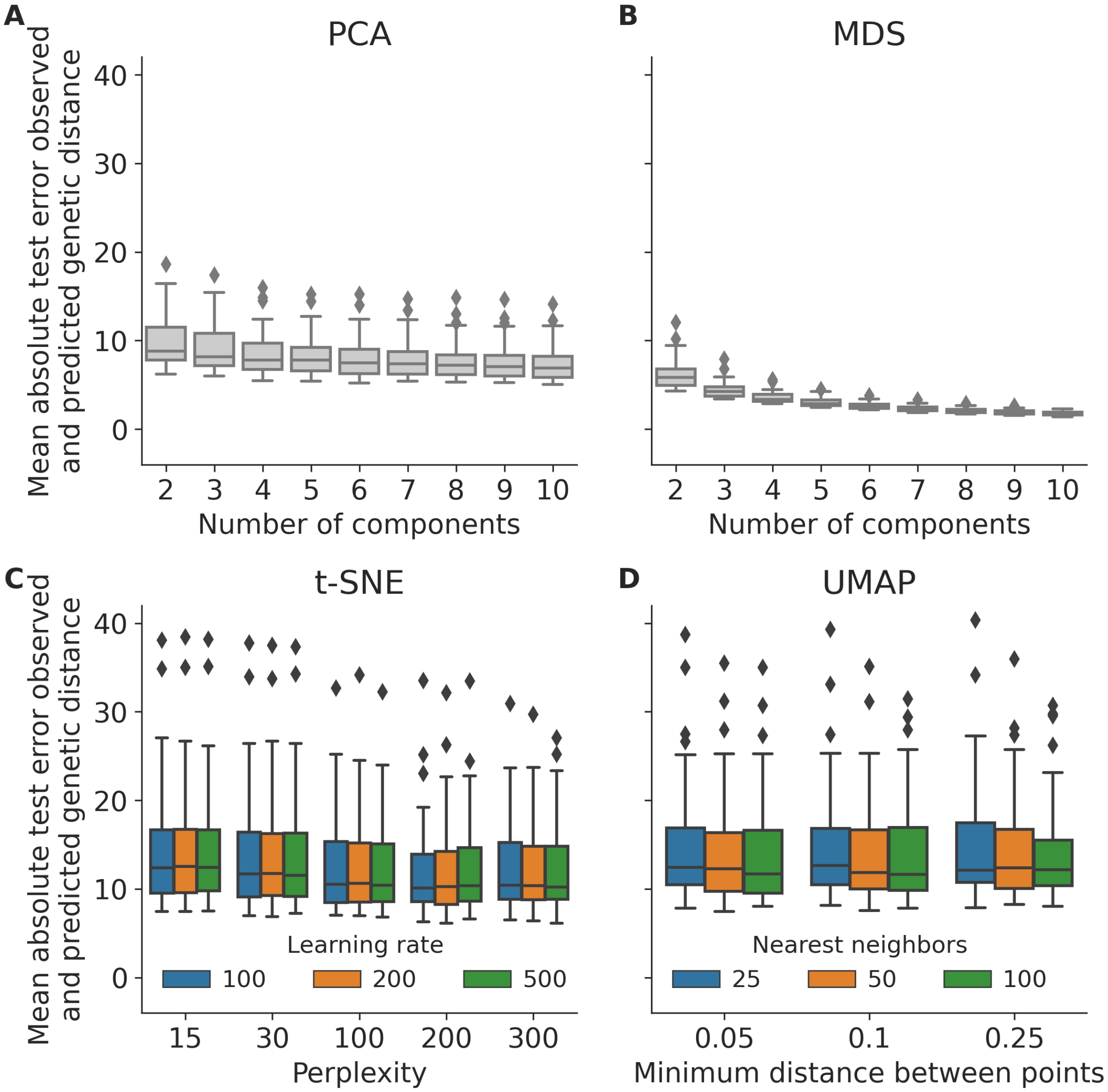


**Supplementary Fig. S 1. Distribution of mean absolute errors (MAE) between observed and predicted pairwise genetic distances per embedding method parameters for simulated influenza-like populations.** For each method and combination of parameters, we created 10 training and test datasets with four years of training data and four years of test data. We created an embedding for each combination of method, parameters, and training/test data, fit a linear model to estimate pairwise genetic distance from pairwise Euclidean distance in the training embedding, estimated the pairwise genetic distance for genomes in the test data based on their Euclidean distances and the linear model fit to the training data, and calculated the mean absolute error (MAE) between estimated and observed genetic distances in the test data. Box plots represent the distribution of MAEs for each combination of method parameters across all training/test datasets. We identified optimal embedding parameters for t-SNE and UMAP as those that minimized the median MAE. Optimal PCA and MDS parameters were the number of components beyond which the median MAE did not descrease by at least 1 nucleotide.

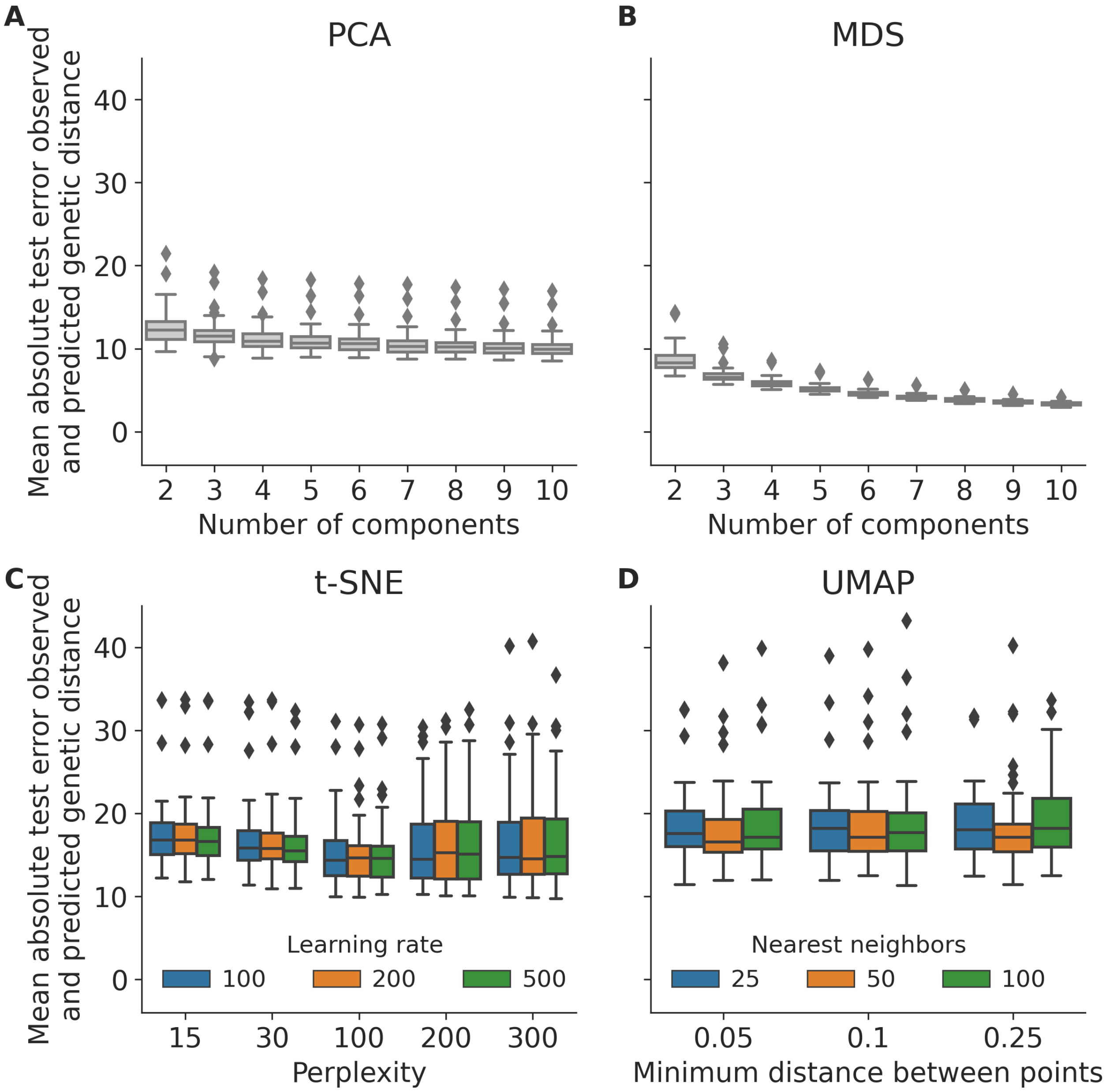


**Supplementary Fig. S 2. Distribution of mean absolute errors (MAE) between observed and predicted pairwise genetic distances per embedding method parameters for simulated coronavirus-like populations.** For each method and combination of parameters, we created 10 training and test datasets with four years of training data and four years of test data. We created an embedding for each combination of method, parameters, and training/test data, fit a linear model to estimate pairwise genetic distance from pairwise Euclidean distance in the training embedding, estimated the pairwise genetic distance for genomes in the test data based on their Euclidean distances and the linear model fit to the training data, and calculated the mean absolute error (MAE) between estimated and observed genetic distances in the test data. Box plots represent the distribution of MAEs for each combination of method parameters across all training/test datasets. We identified optimal embedding parameters for t-SNE and UMAP as those that minimized the median MAE. Optimal PCA and MDS parameters were the number of components beyond which the median MAE did not descrease by at least 1 nucleotide.

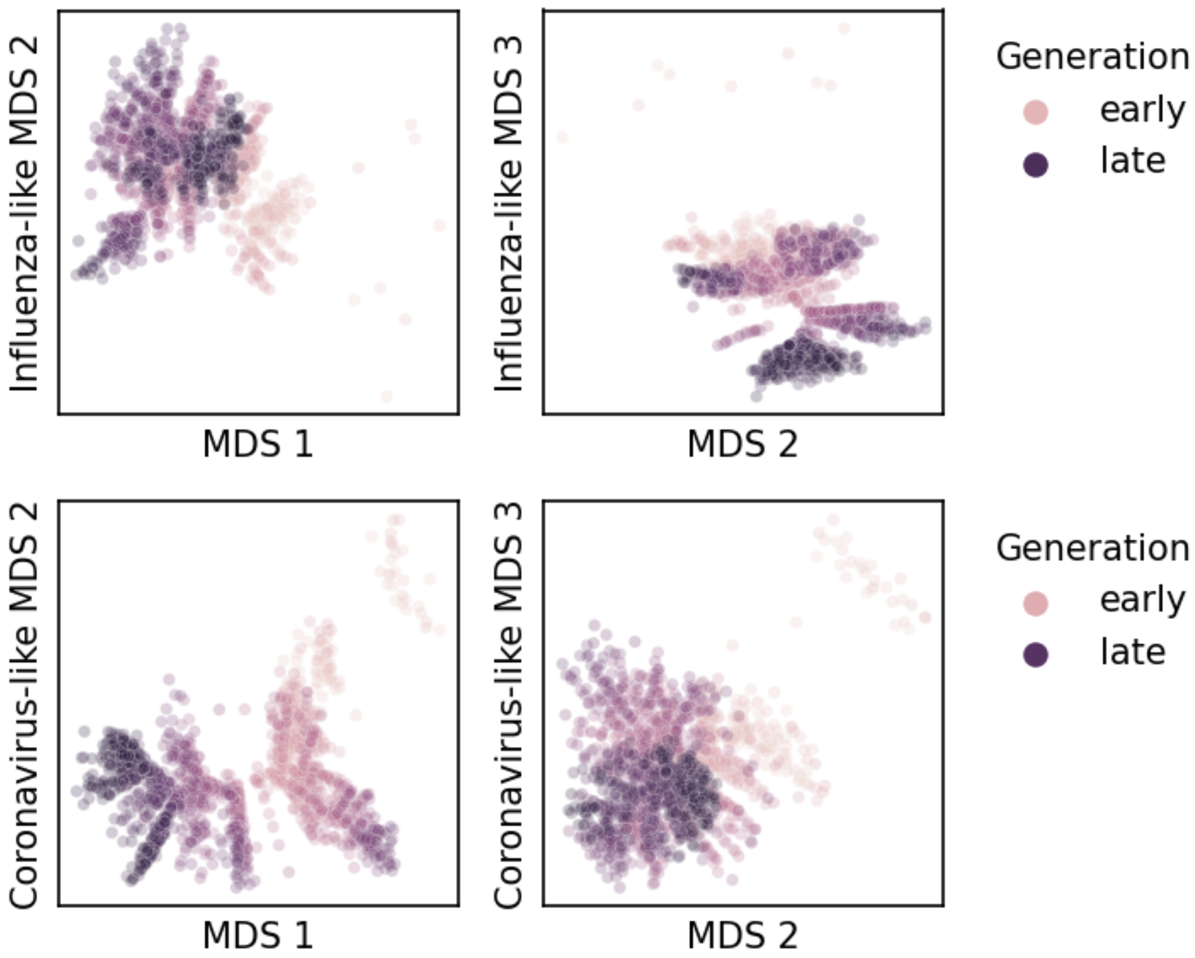


**Supplementary Fig. S 3. Representative MDS embeddings for simulated populations using optimal parameters per pathogen (rows) and showing all three components.** Each panel shows the embedding for sequences from the first four years of a single replicate population for the corresponding pathogen type. Each point represents a simulated viral sequence colored by its generation with darker values representing later generations.

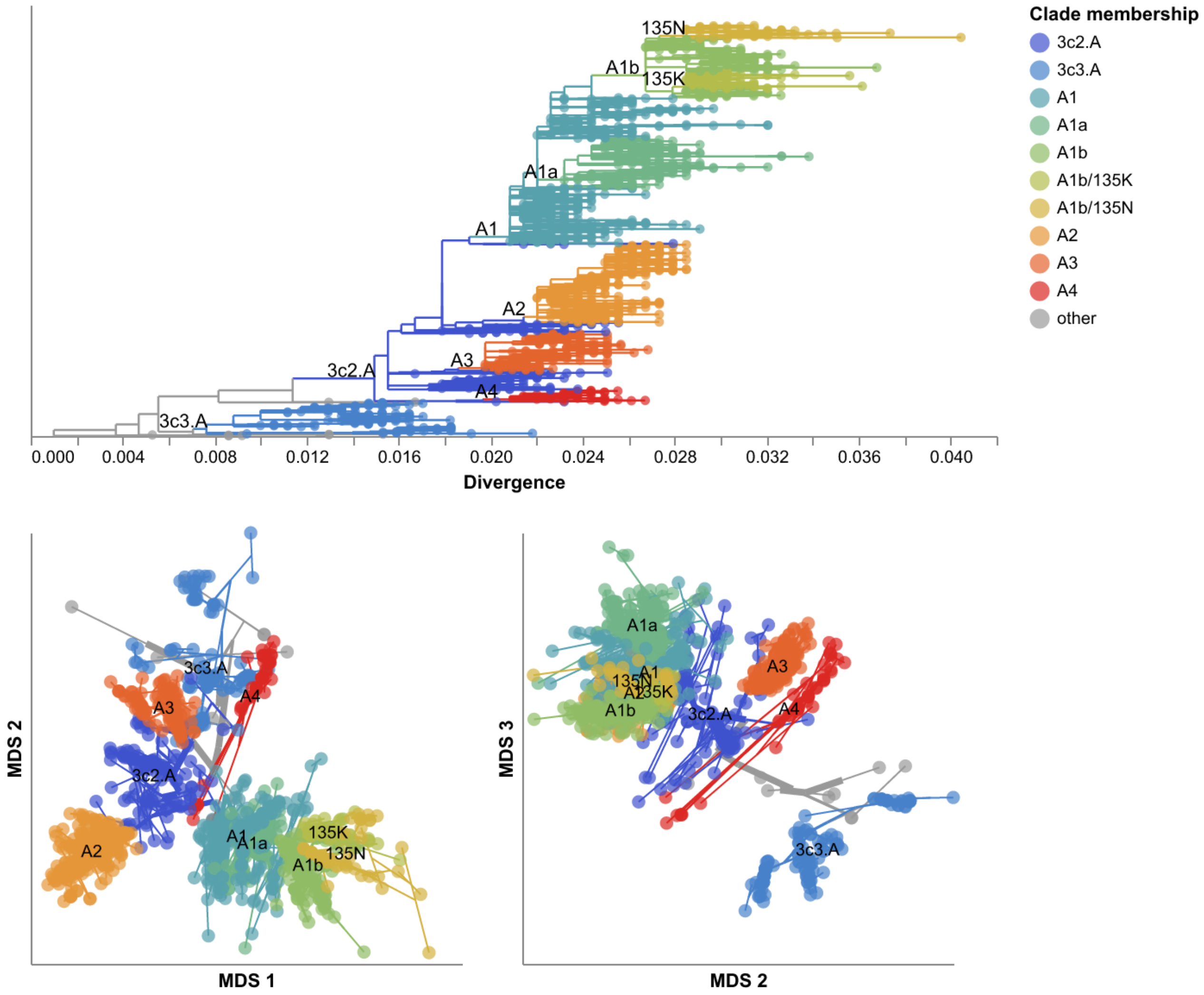


**Supplementary Fig. S 4. MDS embeddings for early (2016–2018) influenza H3N2 HA sequences showing all three components.** Line segments in each embedding reflect phylogenetic relationships with internal node positions calculated from the mean positions of their immediate descendants in each dimension (see Methods). Line colors represent the clade membership of the most ancestral node in the pair of nodes connected by the segment. Line thickness in the embeddings scales by the square root of the number of leaves descending from a given node in the phylogeny. Clade labels appear in the tree at the earliest ancestral node of the tree for each clade. Clade labels appear in each embedding at the average position on the x and y axis for sequences in a given clade.

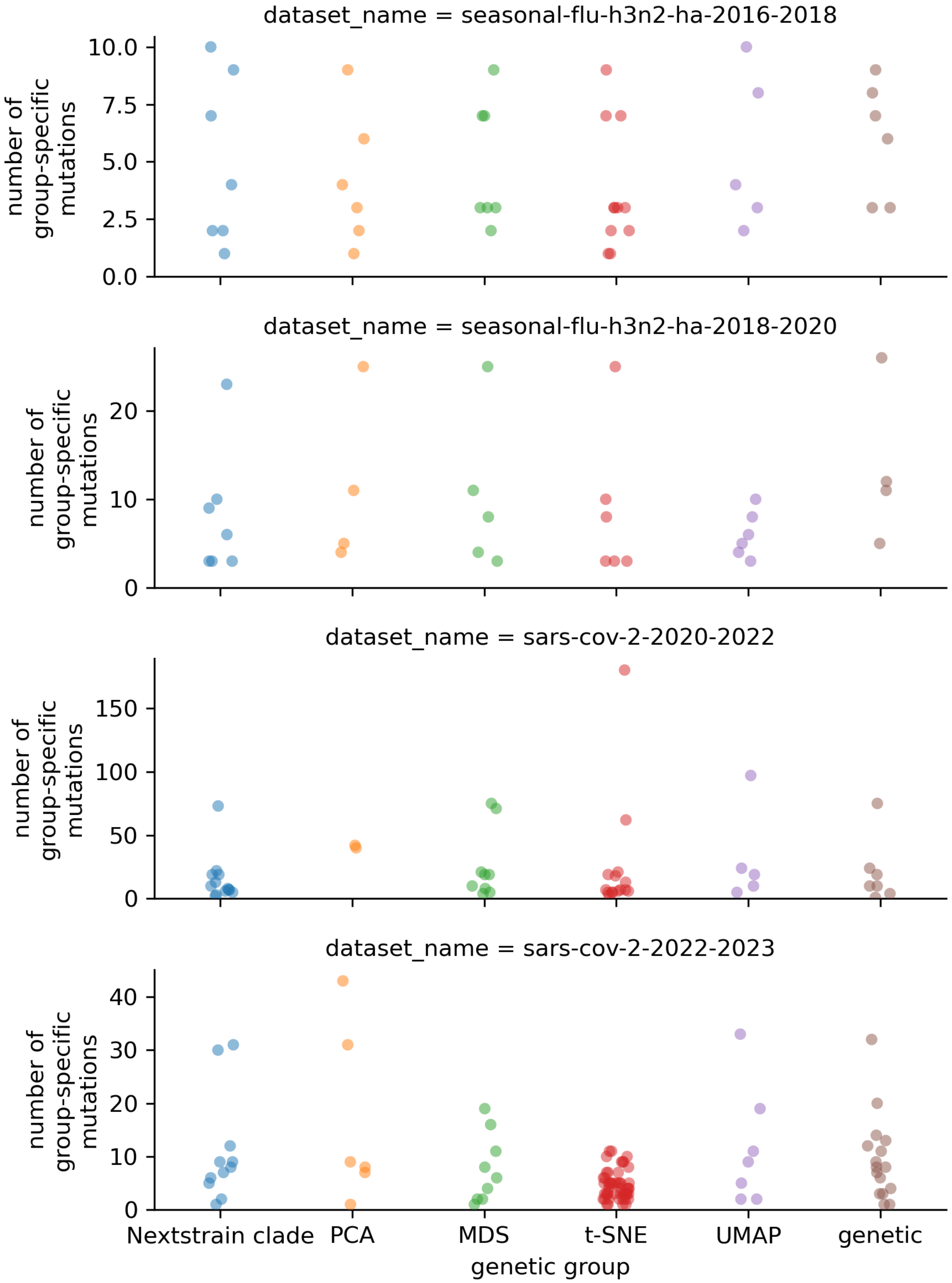


**Supplementary Fig. S 5. Number of group-specific mutations per pathogen dataset (rows) and genetic group type (x-axis).** Each point represents an individual Nextstrain clade or embedding cluster and the number of nucleotide mutations that were specific to that group. These counts come from filtering Supplementary Table S3 to mutations with a cluster count of 1, grouping the remaining records by dataset name and cluster column, and counting the number of records per group.

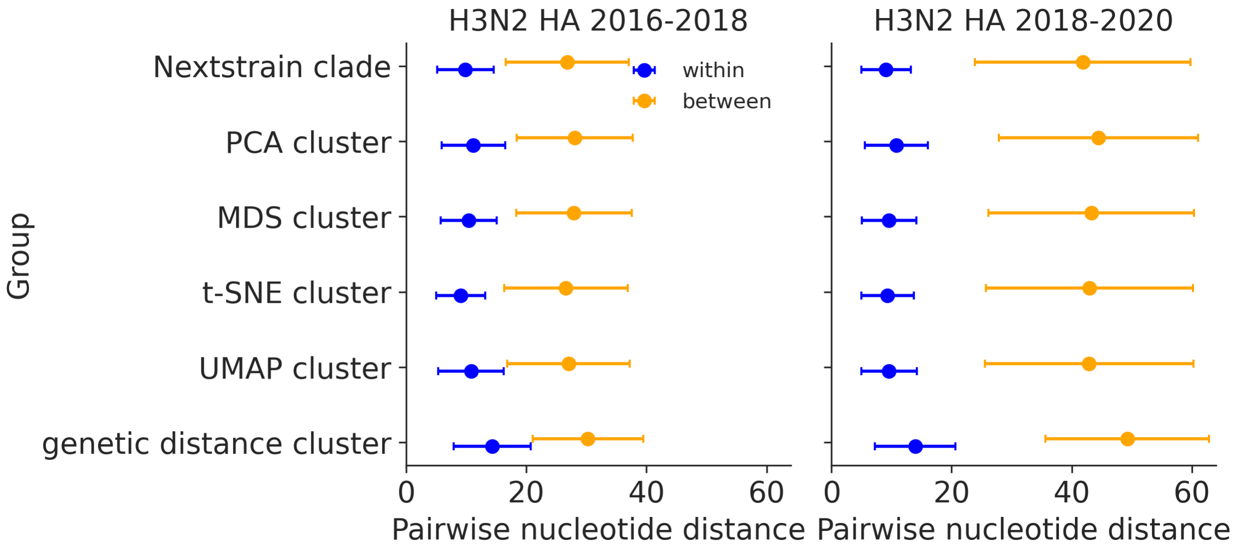


**Supplementary Fig. S 6. Pairwise nucleotide distances for early (2016–2018, left) and late (2018–2020, right) influenza H3N2 HA sequences within and between genetic groups defined by Nextstrain clades and clusters from PCA, MDS, t-SNE, and UMAP embeddings and clusters from pairwise genetic distances.** Each point represents the mean nucleotide distance for pairs of sequences within or between the genetic group in each row. Error bars represent the corresponding standard deviation.

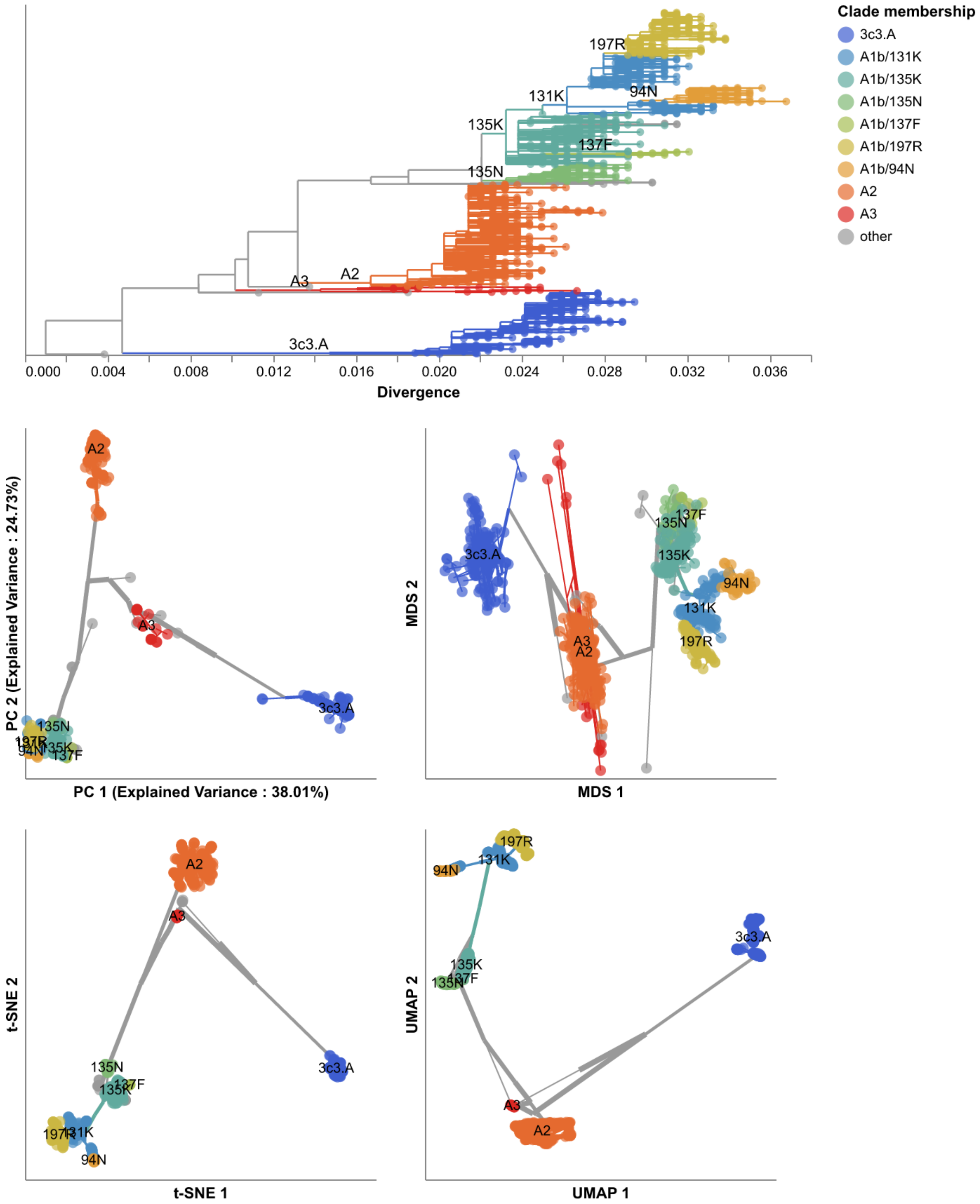


**Supplementary Fig. S 7. Phylogeny of late (2018–2020) influenza H3N2 HA sequences plotted by nucleotide substitutions per site on the x-axis (top) and low-dimensional embeddings of the same sequences by PCA (middle left), MDS (middle right), t-SNE (bottom left), and UMAP (bottom right).** Tips in the tree and embeddings are colored by their Nextstrain clade assignment. Line segments in each embedding reflect phylogenetic relationships with internal node positions calculated from the mean positions of their immediate descendants in each dimension (see Methods). Line colors represent the clade membership of the most ancestral node in the pair of nodes connected by the segment. Line thickness in the embeddings scales by the square root of the number of leaves descending from a given node in the phylogeny. Clade labels appear in the tree at the earliest ancestral node of the tree for each clade. Clade labels appear in each embedding at the average position on the x and y axis for sequences in a given clade.

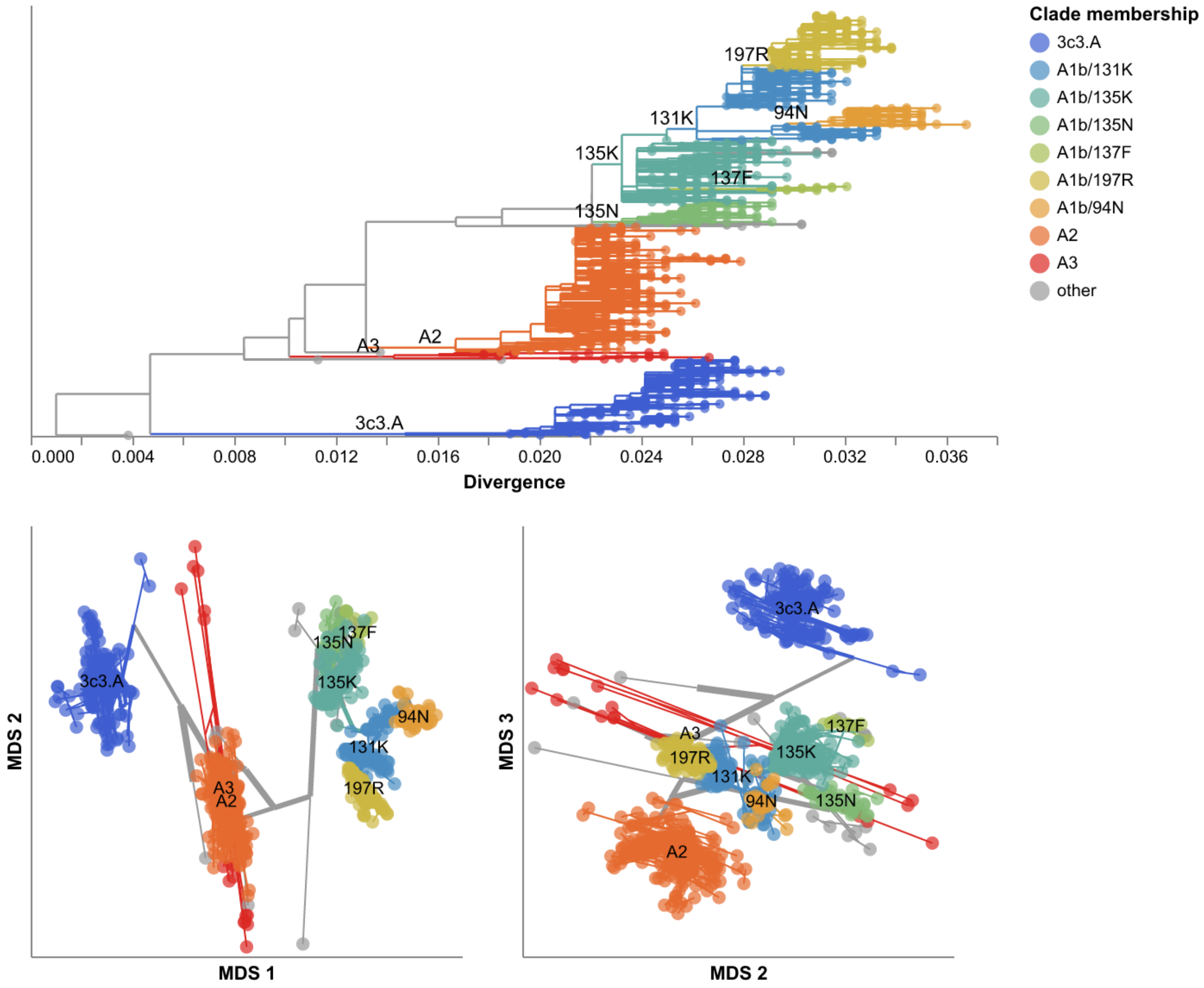


**Supplementary Fig. S 8. MDS embeddings for late (2018–2020) influenza H3N2 HA sequences showing all three components.** Line segments in each embedding reflect phylogenetic relationships with internal node positions calculated from the mean positions of their immediate descendants in each dimension (see Methods). Line colors represent the clade membership of the most ancestral node in the pair of nodes connected by the segment. Line thickness in the embeddings scales by the square root of the number of leaves descending from a given node in the phylogeny. Clade labels appear in the tree at the earliest ancestral node of the tree for each clade. Clade labels appear in each embedding at the average position on the x and y axis for sequences in a given clade.

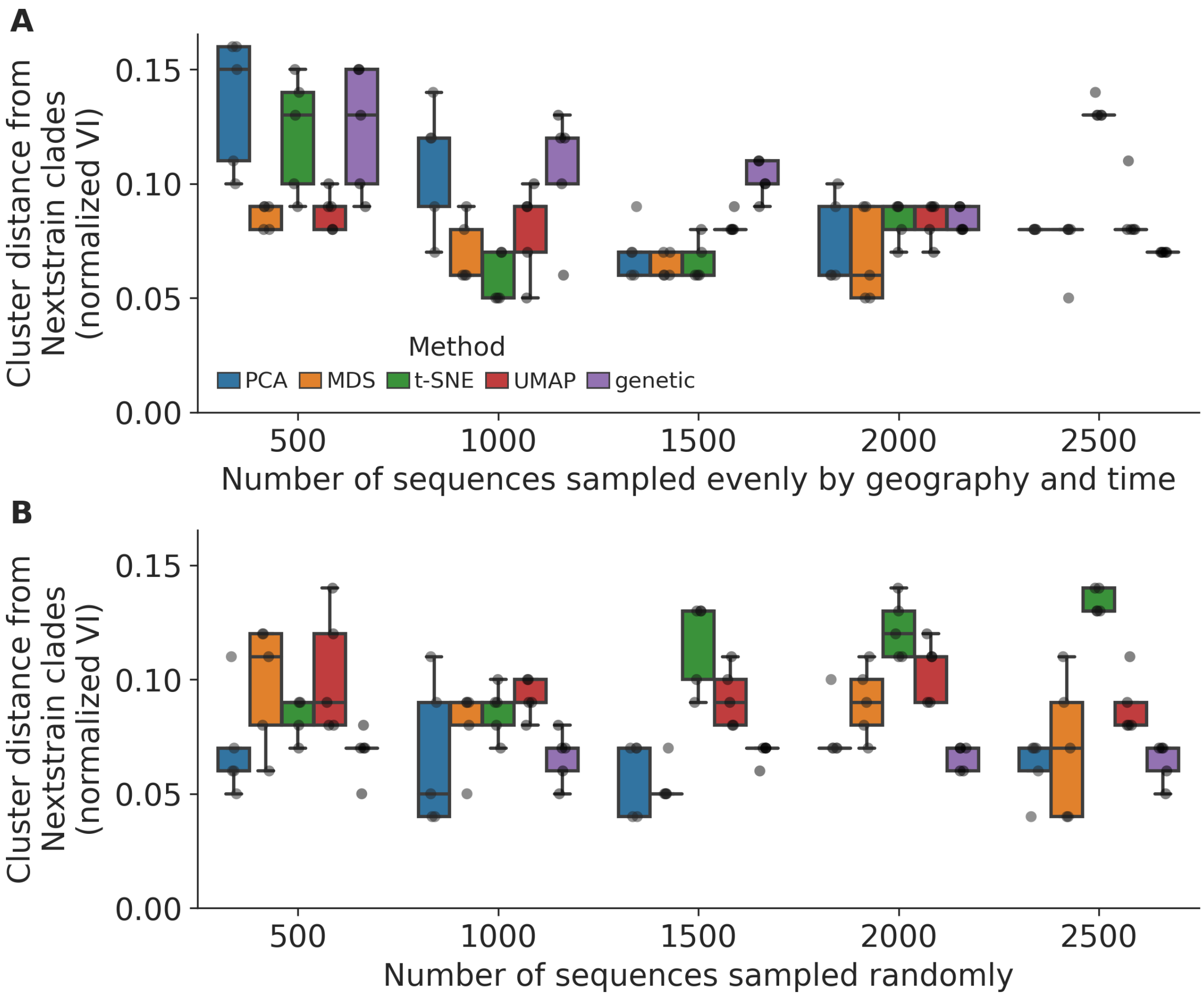


**Supplementary Fig. S 9. Replication of cluster accuracy per embedding method for late (2018–2020) influenza H3N2 HA sequences across different sampling densities (total sequences sampled) and sampling schemes including A) even geographic and temporal sampling and B) random sampling.** We measured cluster accuracy across five replicates per sampling density and scheme with the normalized VI distance between clusters from a given embedding and Nextstrain clades for the same samples. The even sampling scheme selected sequences evenly across country, year, and month to minimize geographic and temporal bias. The random sampling scheme uniformly sampled from the original dataset, reflecting the geographic and genetic bias in those data.

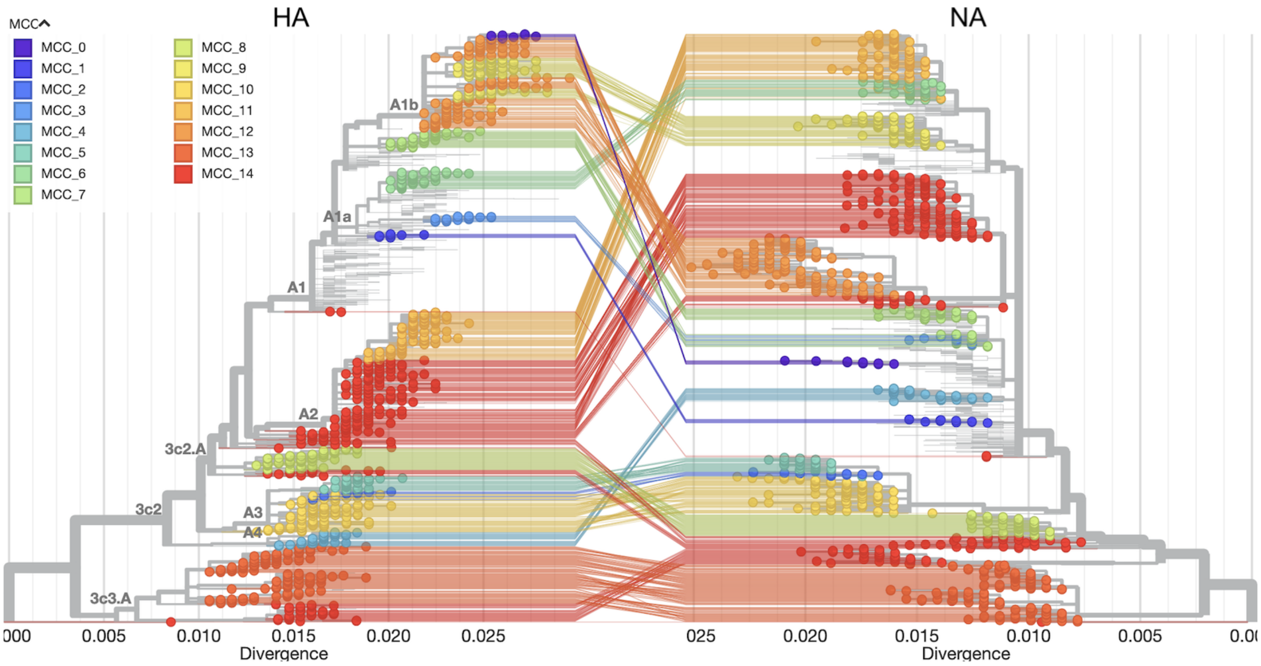


**Supplementary Fig. S 10. Tanglegram view of phylogenetic trees for influenza H3N2 HA (left) and NA (right) with circles representing HA or NA samples, lines connecting the same samples in each tree, and colors showing Maximally Compatible Clades (MCCs) that represent reassortment events identified by TreeKnit.** Samples from MCCs with fewer than 10 sequences appear in gray without circles in the tanglegram. Branch labels in the HA tree show Nextstrain clades to help contextualize placement of each MCC. View an interactive version of this figure on nextstrain.org.

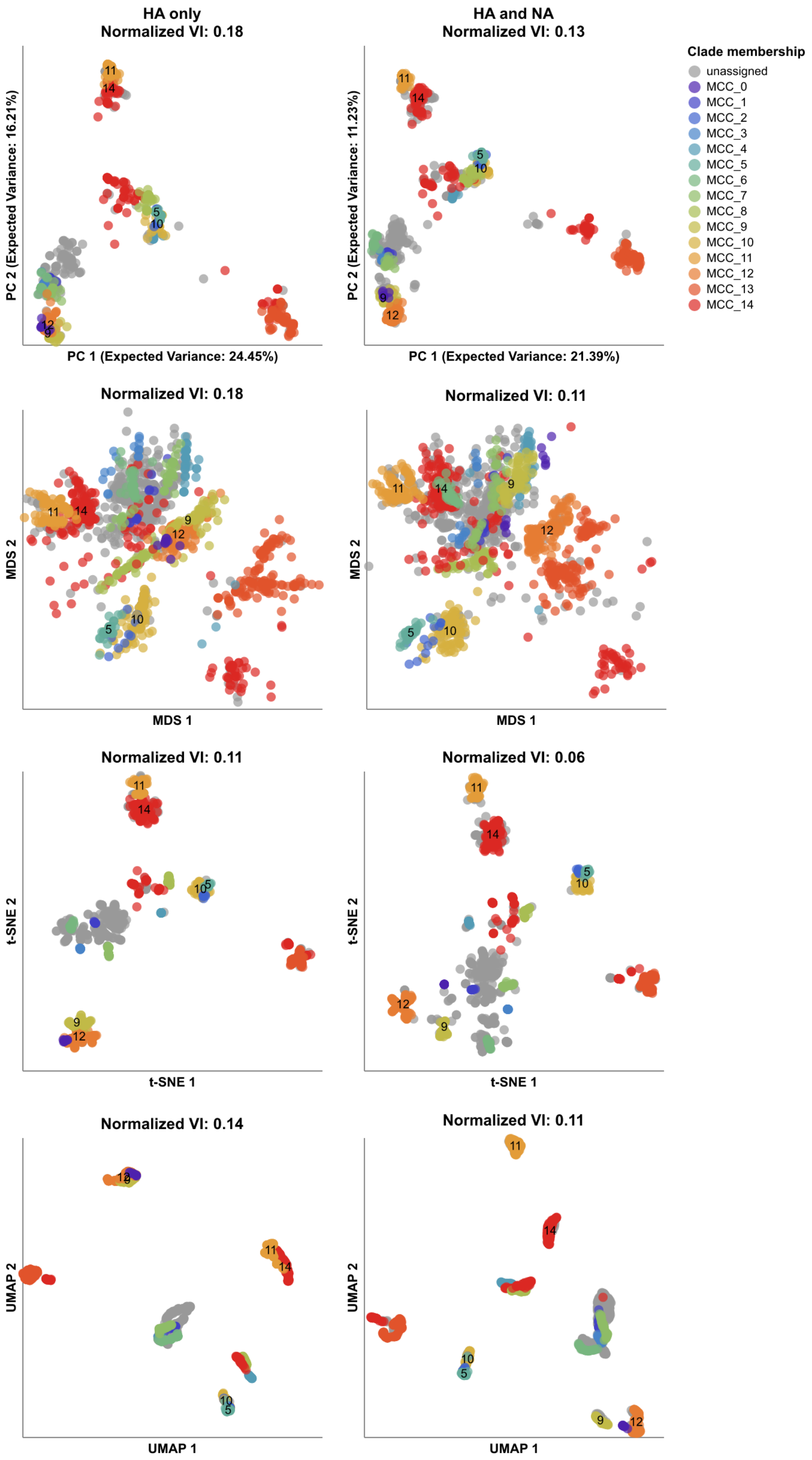


**Supplementary Fig. S 11. Embeddings influenza H3N2 HA-only (left) and combined HA/NA (right) showing the effects of additional NA genetic information on the placement of reassortment events detected by TreeKnit (MCCs).** Sequences from MCCs with fewer than 10 sequences are colored as "unassigned". Normalized VI values quantify the degree to which the combination of HA and NA sequences in an embedding reduces the distance of embedding clusters to TreeKnit reassortment groups represented by MCCs. MCC labels for larger pairs of reassortment events appear in each embedding at the average position on the x and y axis for sequences in a given MCC. MCCs 14 and 11 represent a previously published reassortment event within Nextstrain clade A2 [51]. Labels for MCC 14 represents the sequences from clade A2.

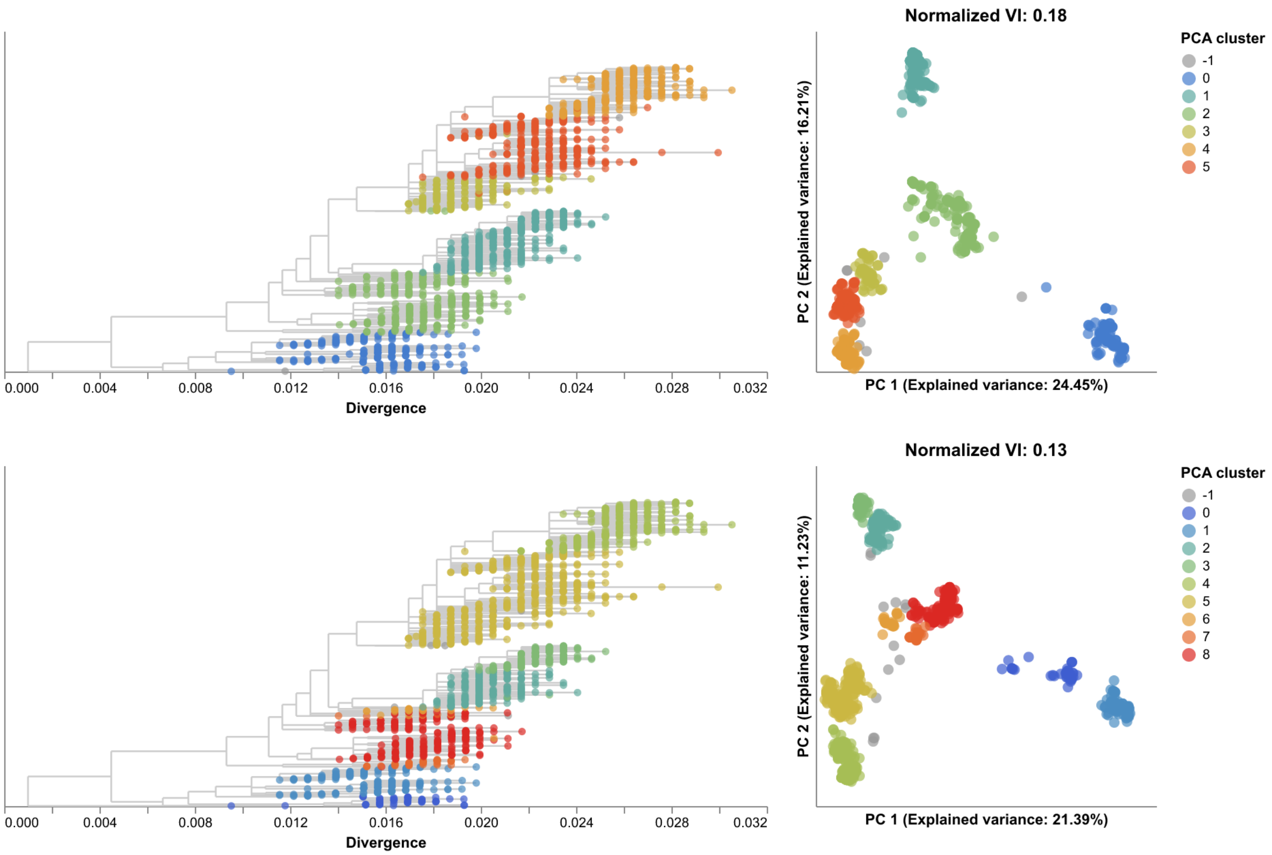


**Supplementary Fig. S 12. PCA embeddings for influenza H3N2 HA sequences only (top row) and HA/NA sequences combined (bottom row) showing the HA trees colored by clusters identified in each embedding (left) and the corresponding embeddings colored by cluster (right).** Normalized VI values quantify the degree to which the combination of HA and NA sequences in an embedding reduces the distance of embedding clusters to TreeKnit reassortment groups represented by MCCs.

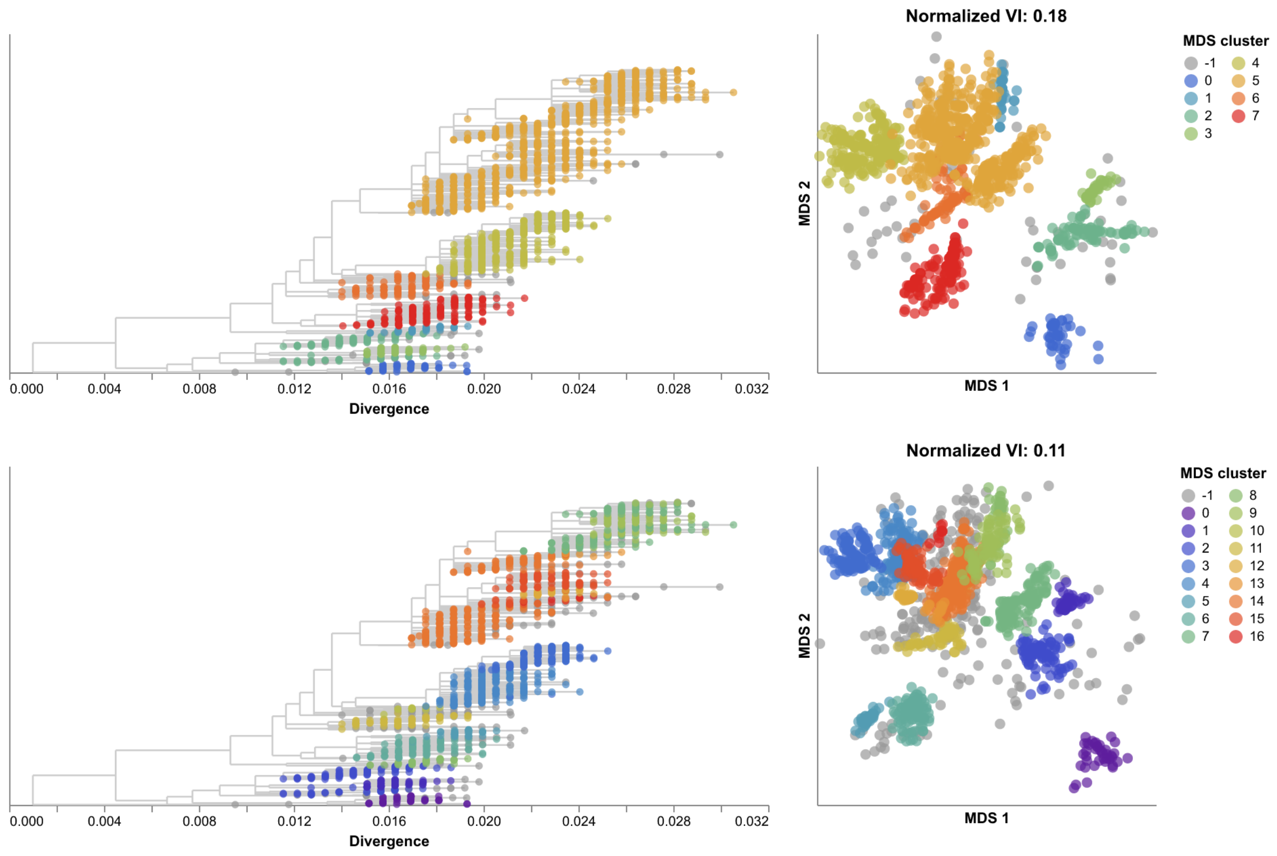


**Supplementary Fig. S 13. MDS embeddings for influenza H3N2 HA sequences only (top row) and HA/NA sequences combined (bottom row) showing the HA trees colored by clusters identified in each embedding (left) and the corresponding embeddings colored by cluster (right).** Normalized VI values quantify the degree to which the combination of HA and NA sequences in an embedding reduces the distance of embedding clusters to TreeKnit reassortment groups represented by MCCs.

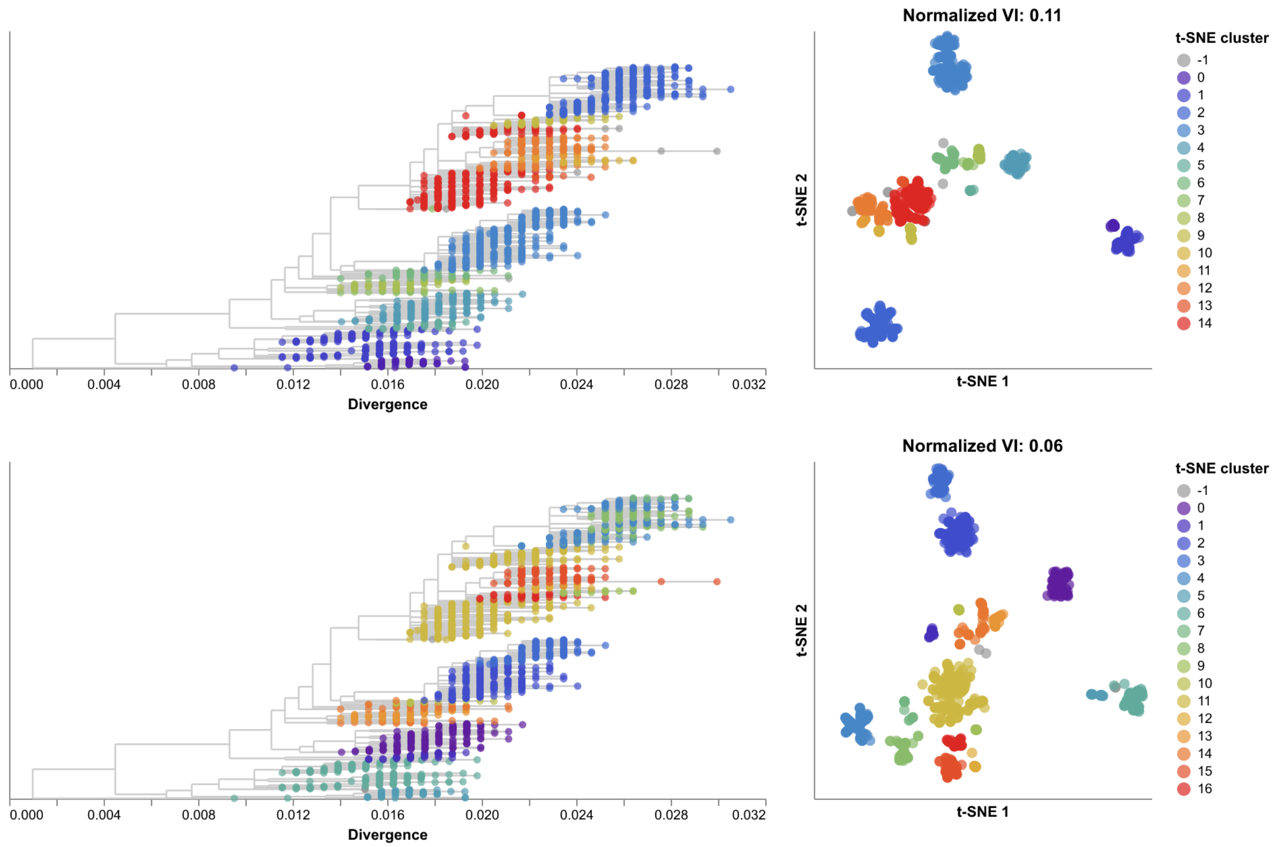


**Supplementary Fig. S 14. t-SNE embeddings for influenza H3N2 HA sequences only (top row) and HA/NA sequences combined (bottom row) showing the HA trees colored by clusters identified in each embedding (left) and the corresponding embeddings colored by cluster (right).** Normalized VI values quantify the degree to which the combination of HA and NA sequences in an embedding reduces the distance of embedding clusters to TreeKnit reassortment groups represented by MCCs.

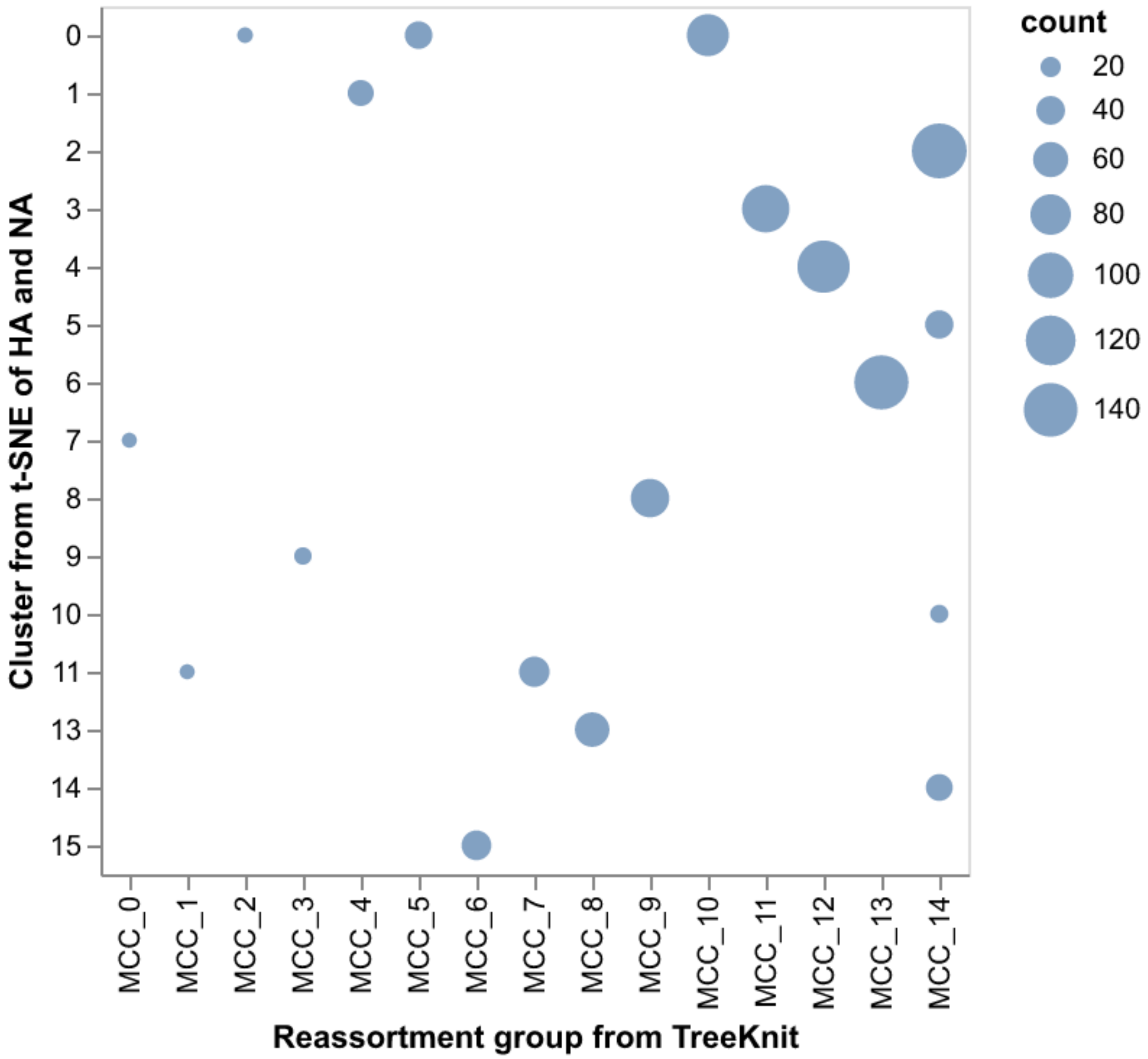


**Supplementary Fig. S 15.** Number of H3N2 strains per combination of reassortment group identified with TreeKnit (MCCs) and cluster identified from joint t-SNE embeddings of HA and NA where the count was at least 10 strains.

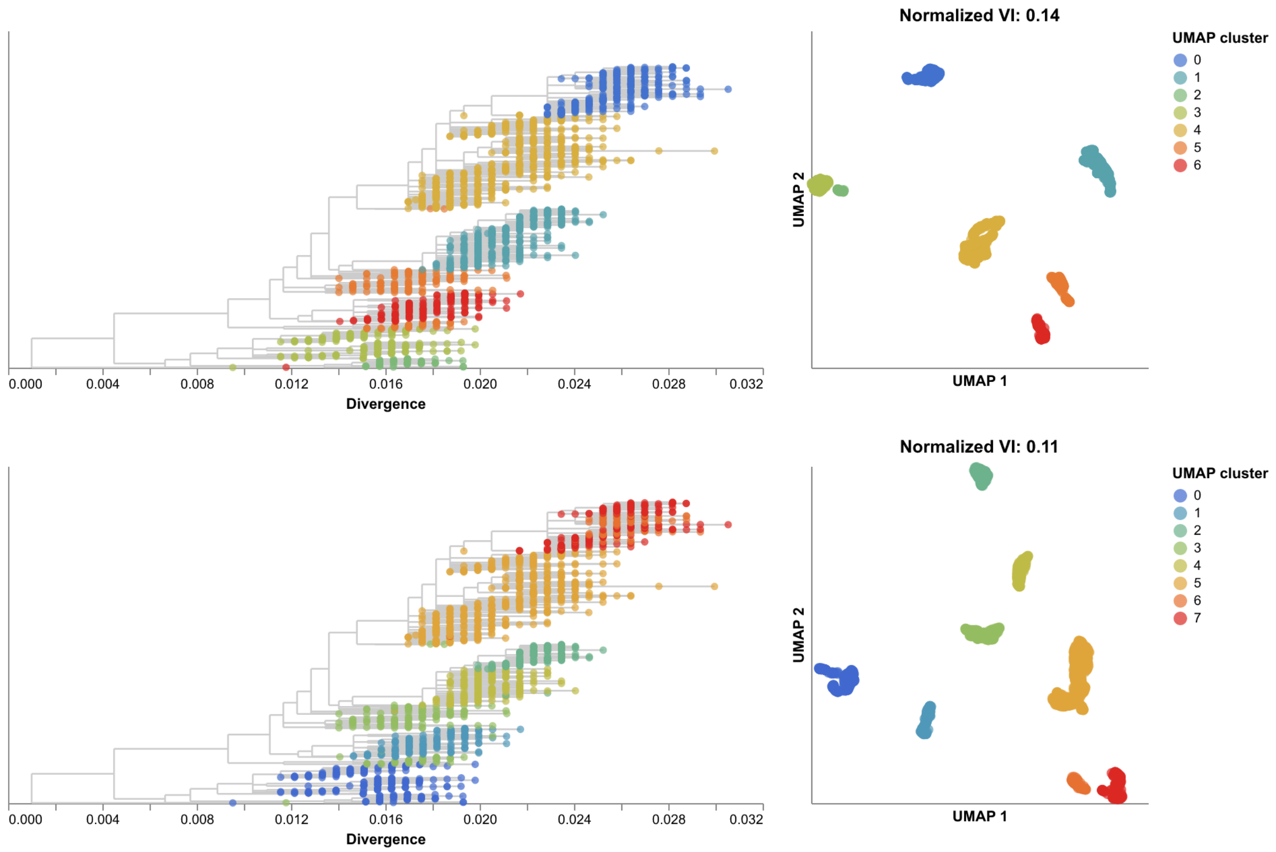


**Supplementary Fig. S 16. UMAP embeddings for influenza H3N2 HA sequences only (top row) and HA/NA sequences combined (bottom row) showing the HA trees colored by clusters identified in each embedding (left) and the corresponding embeddings colored by cluster (right).** Normalized VI values quantify the degree to which the combination of HA and NA sequences in an embedding reduces the distance of embedding clusters to TreeKnit reassortment groups represented by MCCs.

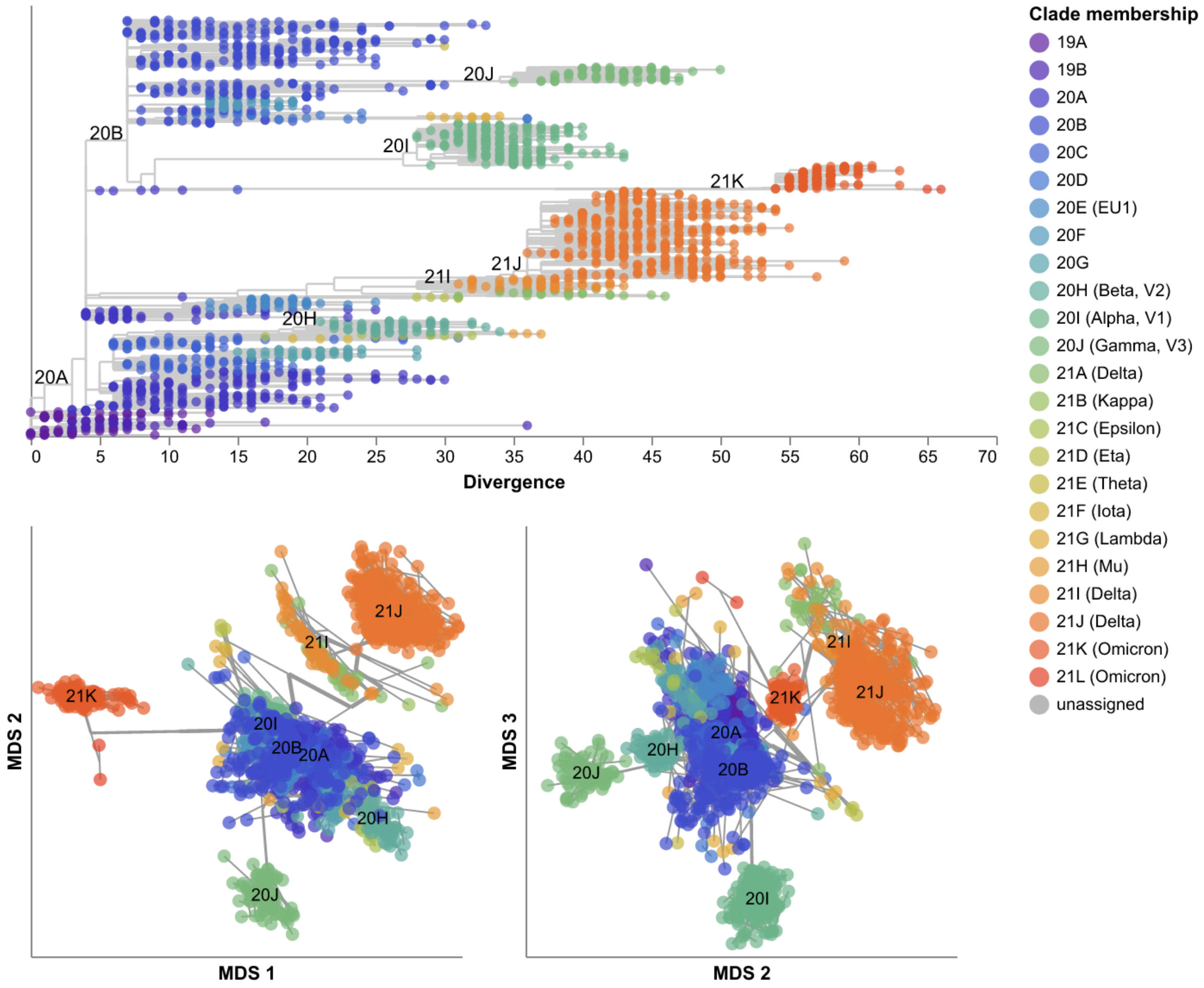


**Supplementary Fig. S 17. MDS embeddings for early SARS-CoV-2 sequences showing all three components.** Line segments in each embedding reflect phylogenetic relationships with internal node positions calculated from the mean positions of their immediate descendants in each dimension (see Methods). Line thickness in the embeddings scales by the square root of the number of leaves descending from a given node in the phylogeny. Clade labels in the tree and embeddings highlight larger clades.

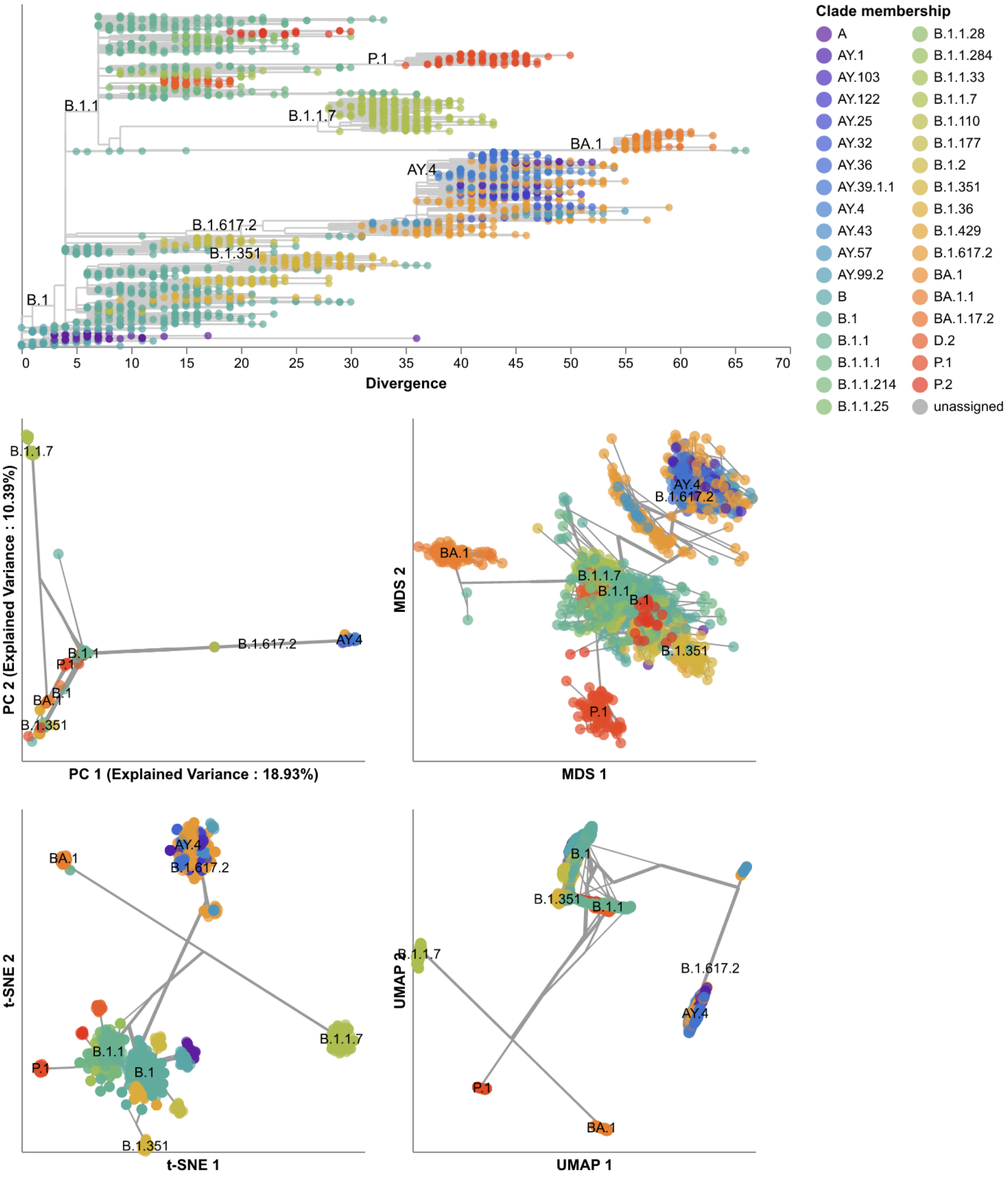


**Supplementary Fig. S 18. Phylogeny of early (2020–2022) SARS-CoV-2 sequences plotted by number of nucleotide substitutions from the most recent common ancestor on the x-axis (top) and low-dimensional embeddings of the same sequences by PCA (middle left), MDS (middle right), t-SNE (bottom left), and UMAP (bottom right).** Tips in the tree and embeddings are colored by their Pango lineage assignment. Line segments in each embedding reflect phylogenetic relationships with internal node positions calculated from the mean positions of their immediate descendants in each dimension (see Methods). Line thickness in the embeddings scales by the square root of the number of leaves descending from a given node in the phylogeny. Clade labels in the tree and embeddings highlight larger Pango lineages.

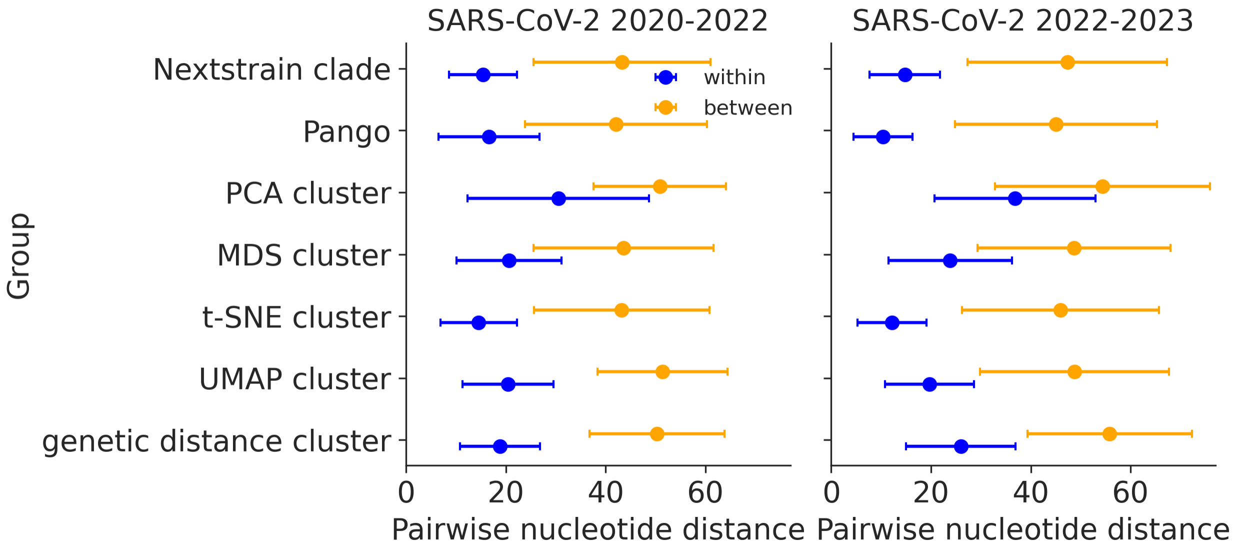


**Supplementary Fig. S 19. Pairwise nucleotide distances for early (2020-2022, left) and late (2022-2023, right) SARS-CoV-2 sequences within and between genetic groups defined by Nextstrain clades, Pango lineages, and clusters from PCA, MDS, t-SNE, and UMAP embeddings and clusters from pairwise genetic distances.** Each point represents the mean nucleotide distance for pairs of sequences within or between the genetic group in each row. Error bars represent the corresponding standard deviation.

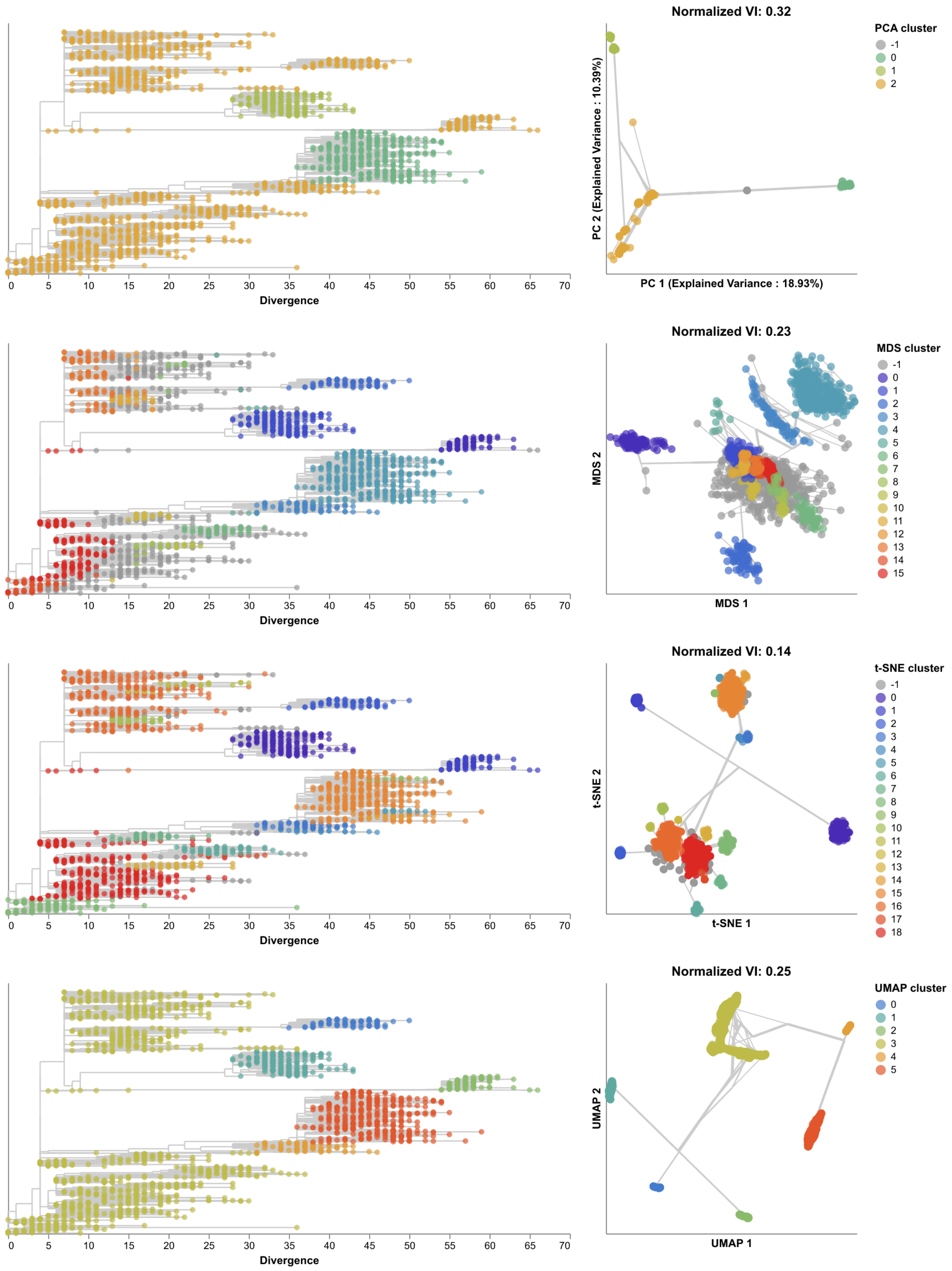


**Supplementary Fig. S 20. Phylogenetic trees (left) and embeddings (right) of early (2020–2022) SARS-CoV-2 sequences colored by HDBSCAN cluster.** Normalized VI values per embedding reflect the distance between clusters and known genetic groups (Pango lineages). Line segments in each embedding reflect phylogenetic relationships with internal node positions calculated from the mean positions of their immediate descendants in each dimension (see Methods). Line thickness in the embeddings scales by the square root of the number of leaves descending from a given node in the phylogeny.

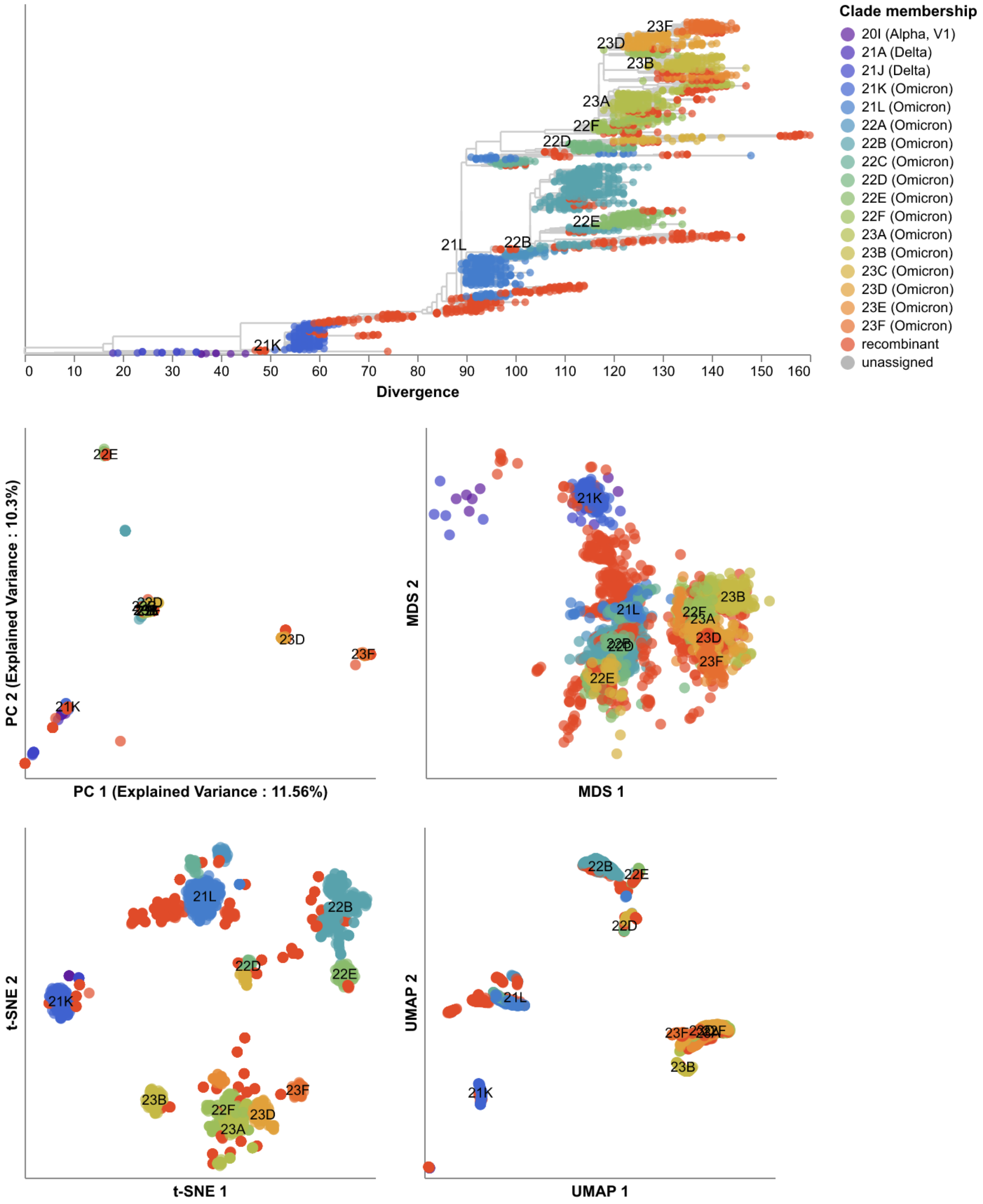


**Supplementary Fig. S 21. Phylogeny of late (2022–2023) SARS-CoV-2 sequences plotted by number of nucleotide substitutions from the most recent common ancestor on the x-axis (top) and low-dimensional embeddings of the same sequences by PCA (middle left), MDS (middle right), t-SNE (bottom left), and UMAP (bottom right).** Tips in the tree and embeddings are colored by their Nextstrain clade assignment. Tips that could not be assigned to a predefined Nextstrain clade due to recombination were colored as "recombinant". Line segments in each embedding reflect phylogenetic relationships with internal node positions calculated from the mean positions of their immediate descendants in each dimension (see Methods). Line thickness in the embeddings scales by the square root of the number of leaves descending from a given node in the phylogeny. Clade labels in the tree and embeddings highlight larger clades.

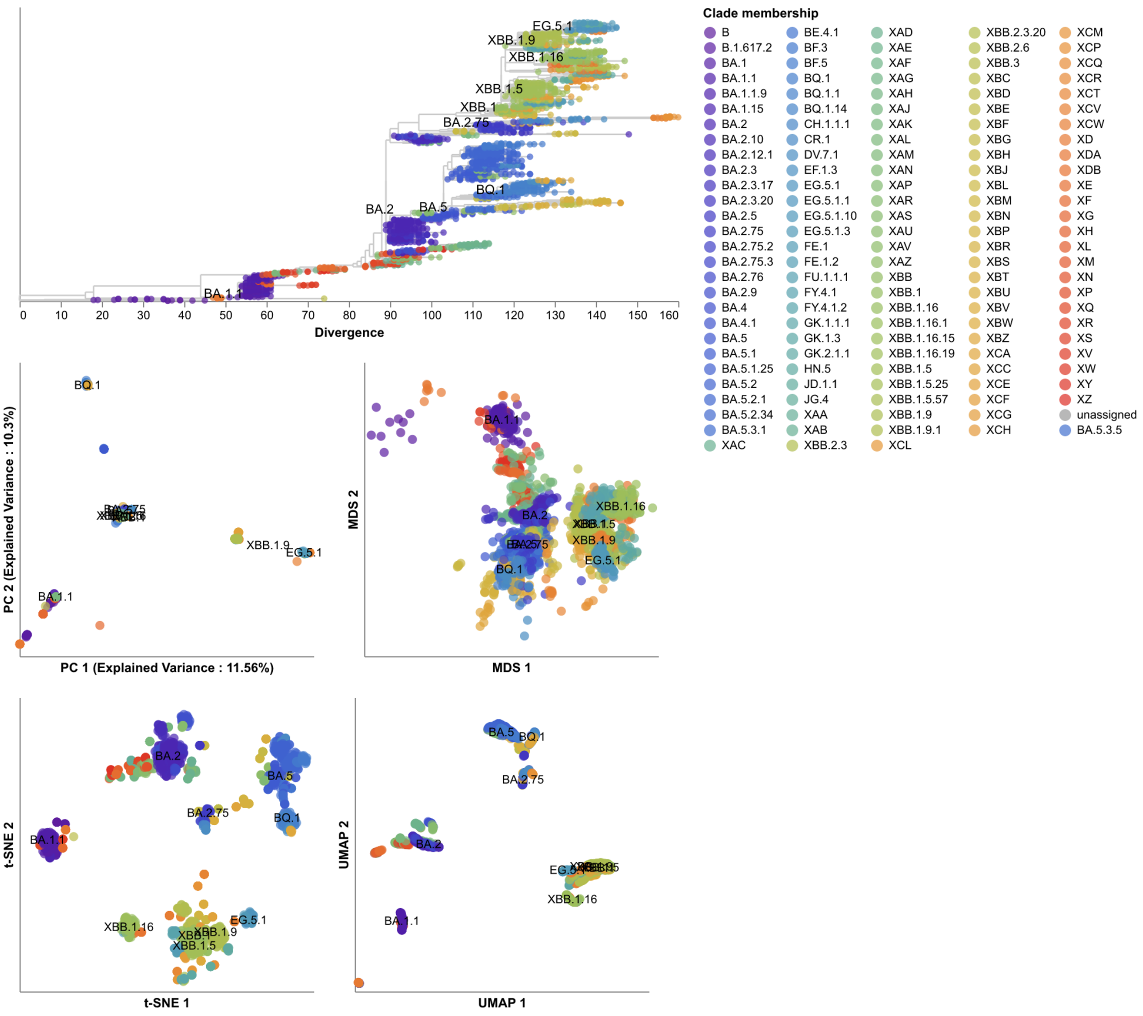


**Supplementary Fig. S 22. Phylogeny of late (2022–2023) SARS-CoV-2 sequences plotted by number of nucleotide substitutions from the most recent common ancestor on the x-axis (top) and low-dimensional embeddings of the same sequences by PCA (middle left), MDS (middle right), t-SNE (bottom left), and UMAP (bottom right).** Tips in the tree and embeddings are colored by their Pango lineage assignment. Line segments in each embedding reflect phylogenetic relationships with internal node positions calculated from the mean positions of their immediate descendants in each dimension (see Methods). Line thickness in the embeddings scales by the square root of the number of leaves descending from a given node in the phylogeny. Clade labels in the tree and embeddings highlight larger Pango lineages.

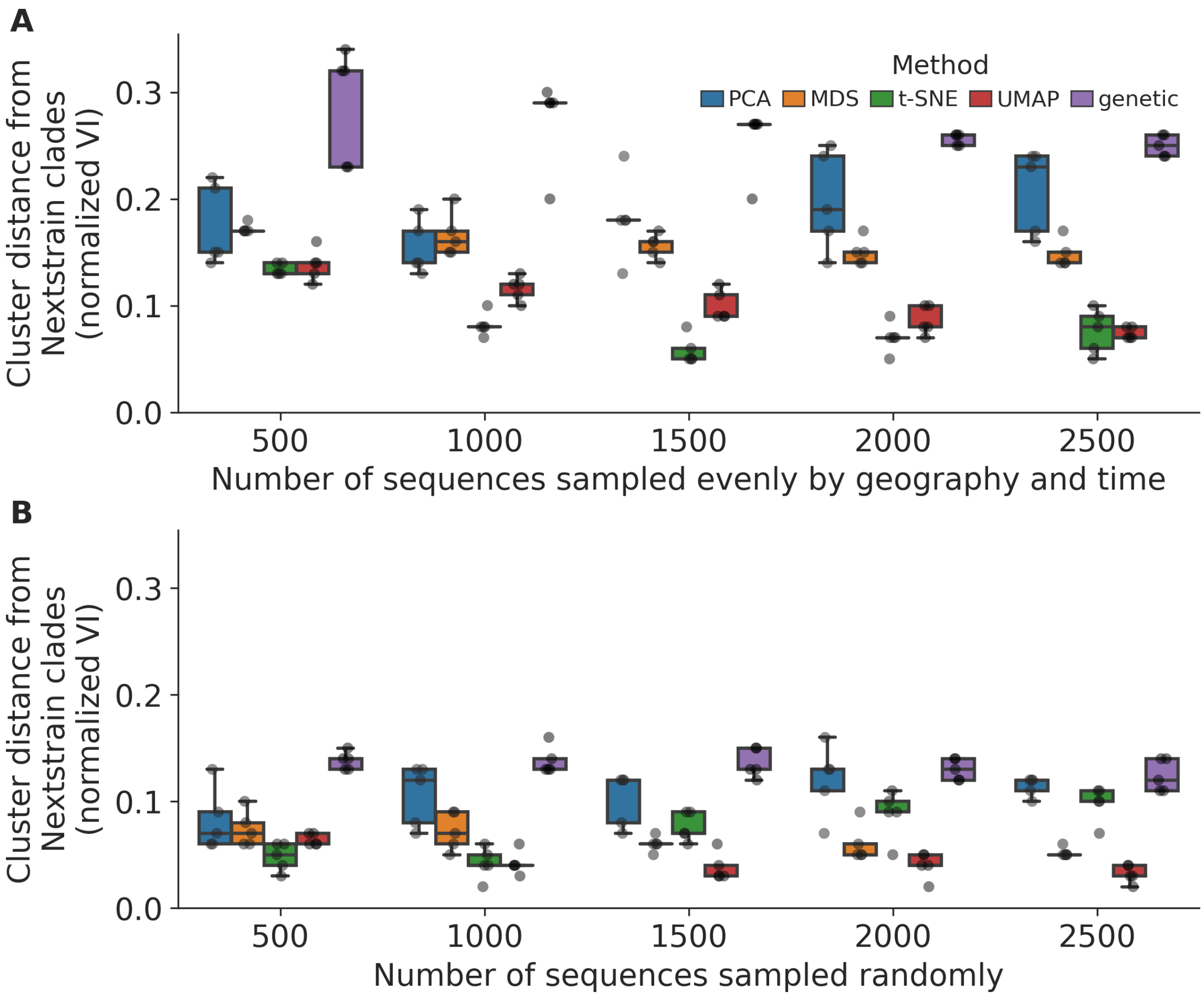


**Supplementary Fig. S 23. Replication of cluster accuracy per embedding method for late (2022–2023) SARS-CoV-2 sequences across different sampling densities (total sequences sampled) and sampling schemes including A) even geographic and temporal sampling and B) random sampling.** We measured cluster accuracy across five replicates per sampling density and scheme with the normalized VI distance between clusters from a given embedding and Nextstrain clades for the same samples. The even sampling scheme selected sequences evenly across region, year, and month to minimize geographic and temporal bias. The random sampling scheme uniformly sampled from the original dataset, reflecting the geographic and genetic bias in those data.

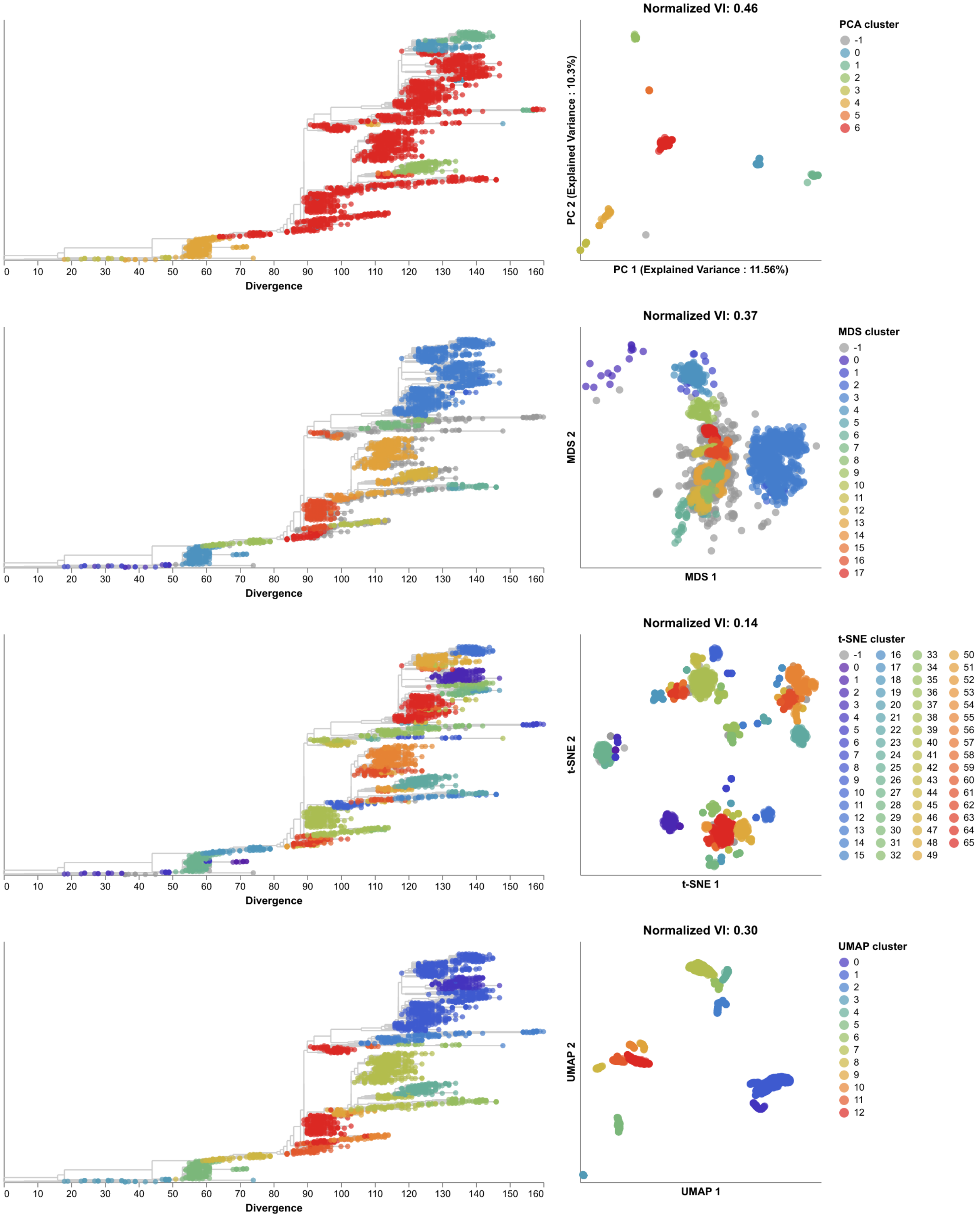


**Supplementary Fig. S 24. Phylogenetic trees (left) and embeddings (right) of late (2022–2023) SARS-CoV-2 sequences colored by HDBSCAN cluster.** Normalized VI values per embedding reflect the distance between clusters and known genetic groups (Pango lineages).

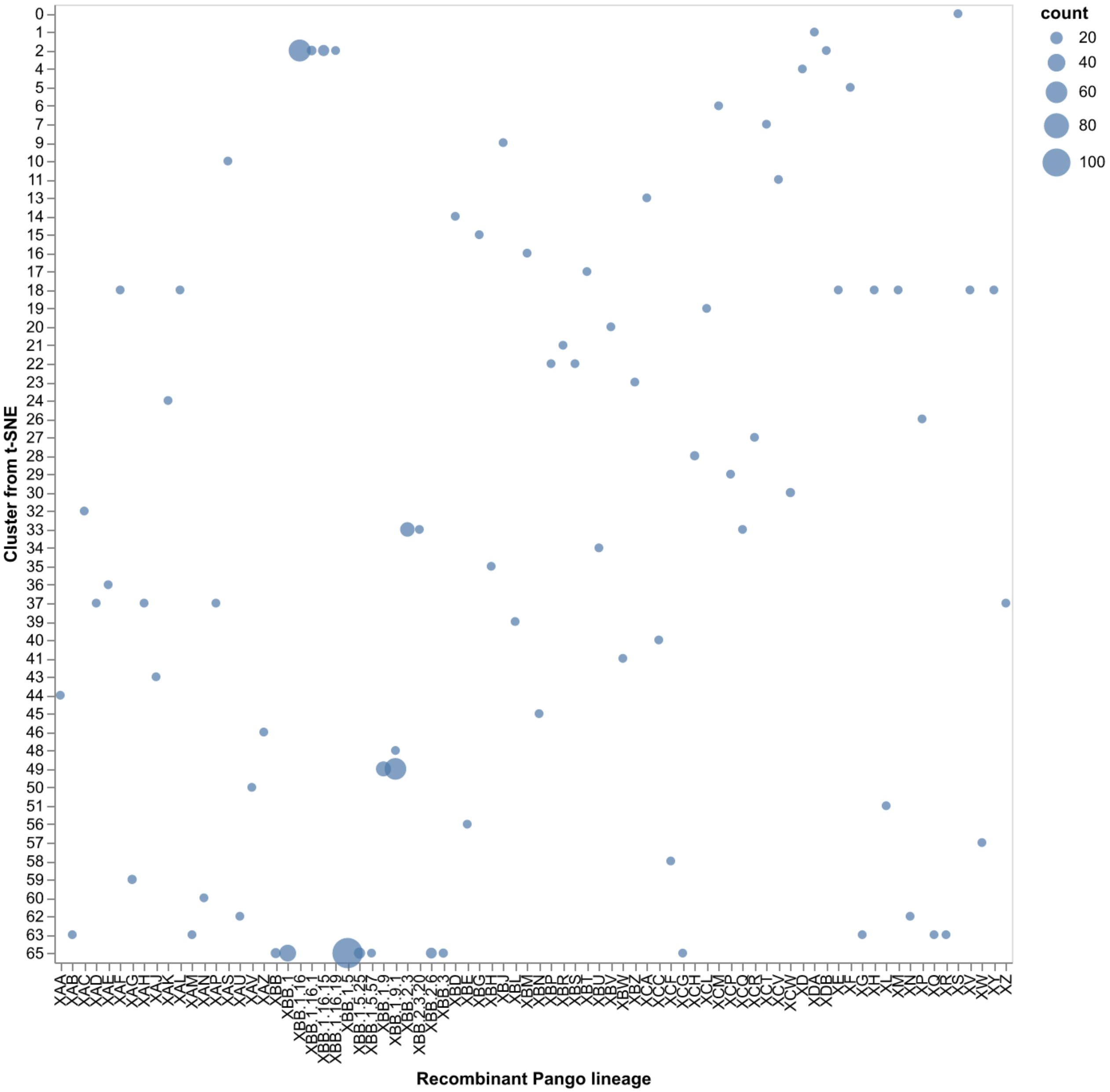


**Supplementary Fig. S 25.** Number of late SARS-CoV-2 strains per combination of recombinant Pango lineage and t-SNE cluster where the count was at least 10 strains.

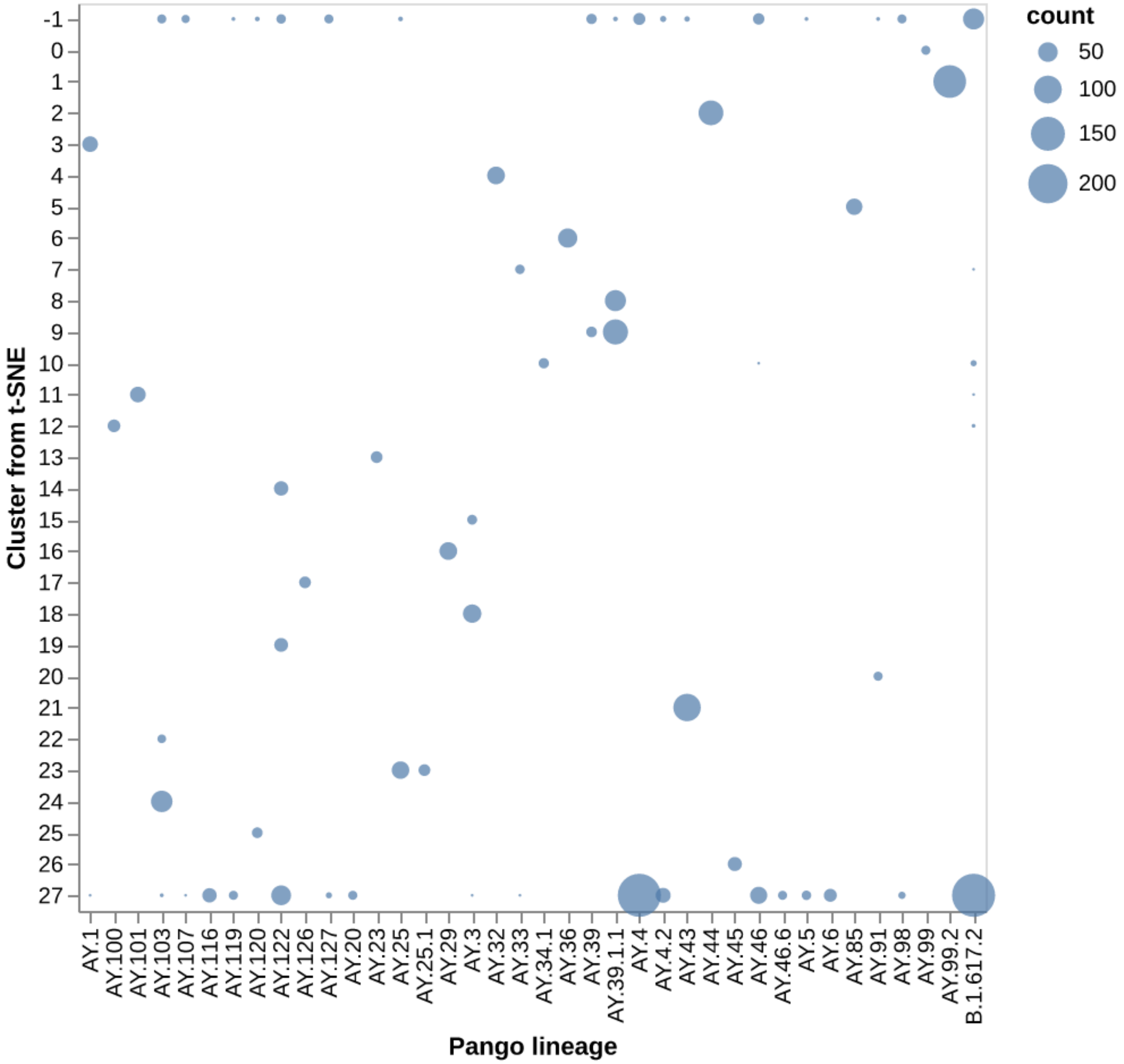


**Supplementary Fig. S 26. Number of SARS-CoV-2 sequences from Nextstrain clade 21J (Delta) per combination of Pango lineage and t-SNE cluster showing the degree to which clusters of sequences from a single Nextstrain clade can capture Pango-resolution genetic groups.** The size of each circle represents the number of sequences associated with each pair of t-SNE cluster and Pango lineage. The cluster label "-1" represents sequences that could not be assigned to a cluster.

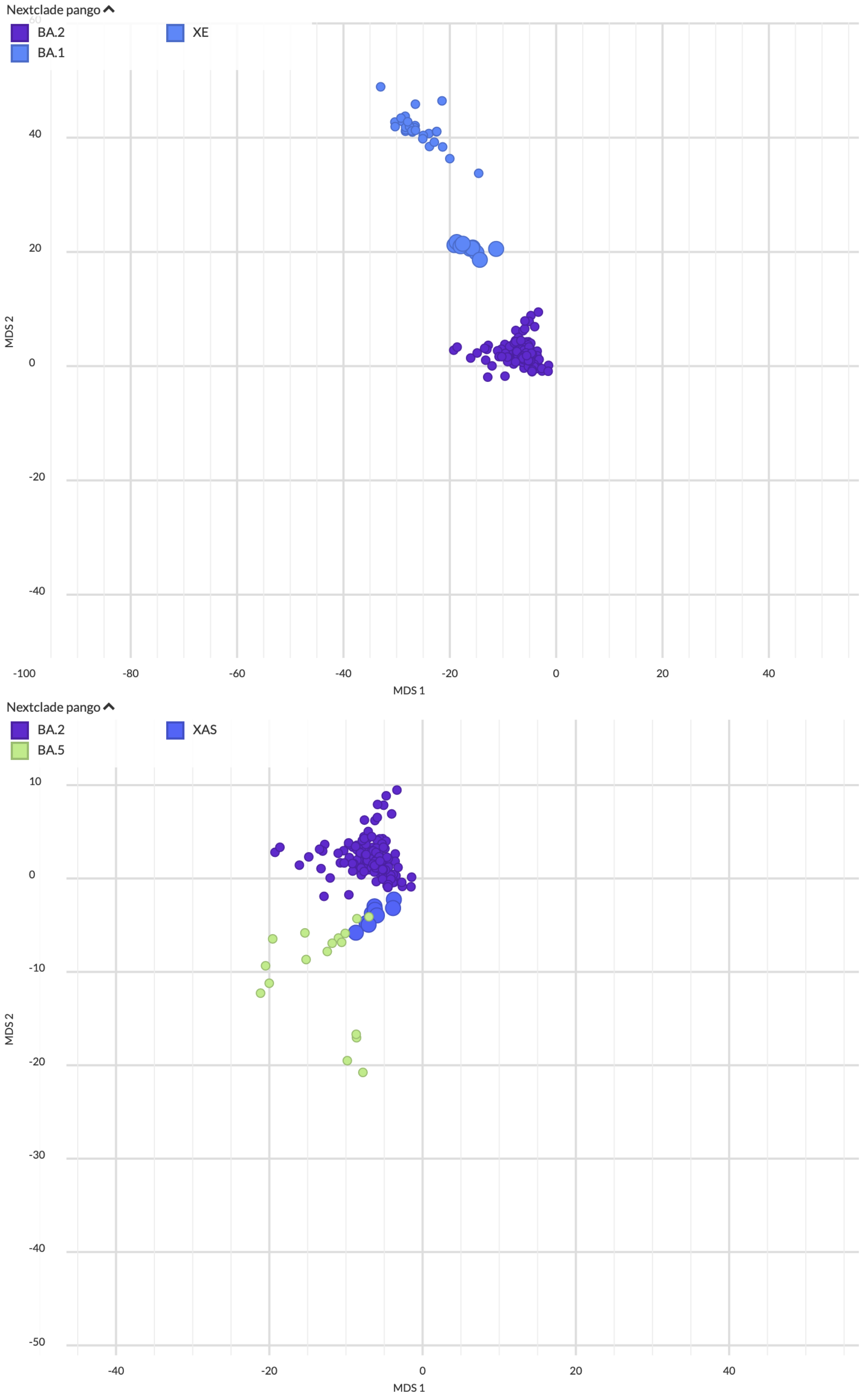


**Supplementary Fig. S 27. Representative MDS embeddings (first two dimensions) of late SARS-CoV-2 samples filtered to known recombinant lineages and their parental lineages including A) XE which descended from BA.1 and BA.2 and B) XAS which descended from BA.2 and BA.5.** Larger circles represent the recombinant lineages in each figure.
